# Structure-Activity Relationship Studies Towards Analogues of Pleconaril as Novel Enterovirus-D68 Capsid-Targeting Antivirals

**DOI:** 10.1101/2025.08.08.668114

**Authors:** David L. Cousins, Ed J. Griffen, Jessica Stacey, Alpha A. Lee, Yuliia Filimonova, Anton Hlavin, Yuliia Holota, Roman Khmil, Mykyta Kordubailo, Oleksii Kostinov, Dmytro Lesyk, Ivan Logvinenko, Mariia Lototska, Viacheslav Lysenko, Anna Pashchenko, Mariia Pavlichenko, Anzhela Rodnichenko, Anton Tkachenko, Brett L. Hurst, Justin G. Julander, Hong Wang, Rebecca Pearl, Jared Benjamin, Randy Diaz-Tapia, Mary E. Gordon, Randy A. Albrecht, Kris White

## Abstract

Non-polio enteroviruses (NPEV) such as enterovirus D68 (EV-D68) that are highly infectious and associated with polio-like neurological complications have caused outbreaks, globally, in recent years. While some clinical and preclinical compounds have shown efficacy against NPEV *in-vitro*, liabilities that caused historical compounds such as pleconaril to fall short of FDA approval still remain. We present herein SAR and SPR studies of analogues of clinical compounds such as pleconaril and vapendavir against EV-D68 as a representative NPEV. Numerous structurally differentiated analogues with EV-D68 antiviral activity and useful ADME properties were discovered, which could serve as starting points for future EV drug discovery campaigns. Screening against a panel of enteroviruses revealed moderately broad-spectrum anti-EV activity of compound 26.

## Introduction

Poliovirus has almost been eradicated through immunisation and other approaches.^1–3^ There are, however, a number of emerging non-polio enteroviruses (NPEV) belonging to the same viral family (*Picornaviridae*) whose outbreaks have been confirmed globally in recent years (Figure 1) and represent a pandemic risk. ^4–13^ These include Enterovirus-D68 (EV-D68), Enterovirus-A71 (EV-A71), Coxsackievirus-A6 (CV-A6), Coxsackievirus-B3 (CV-B3), variants thereof and more.

**Figure 1:**
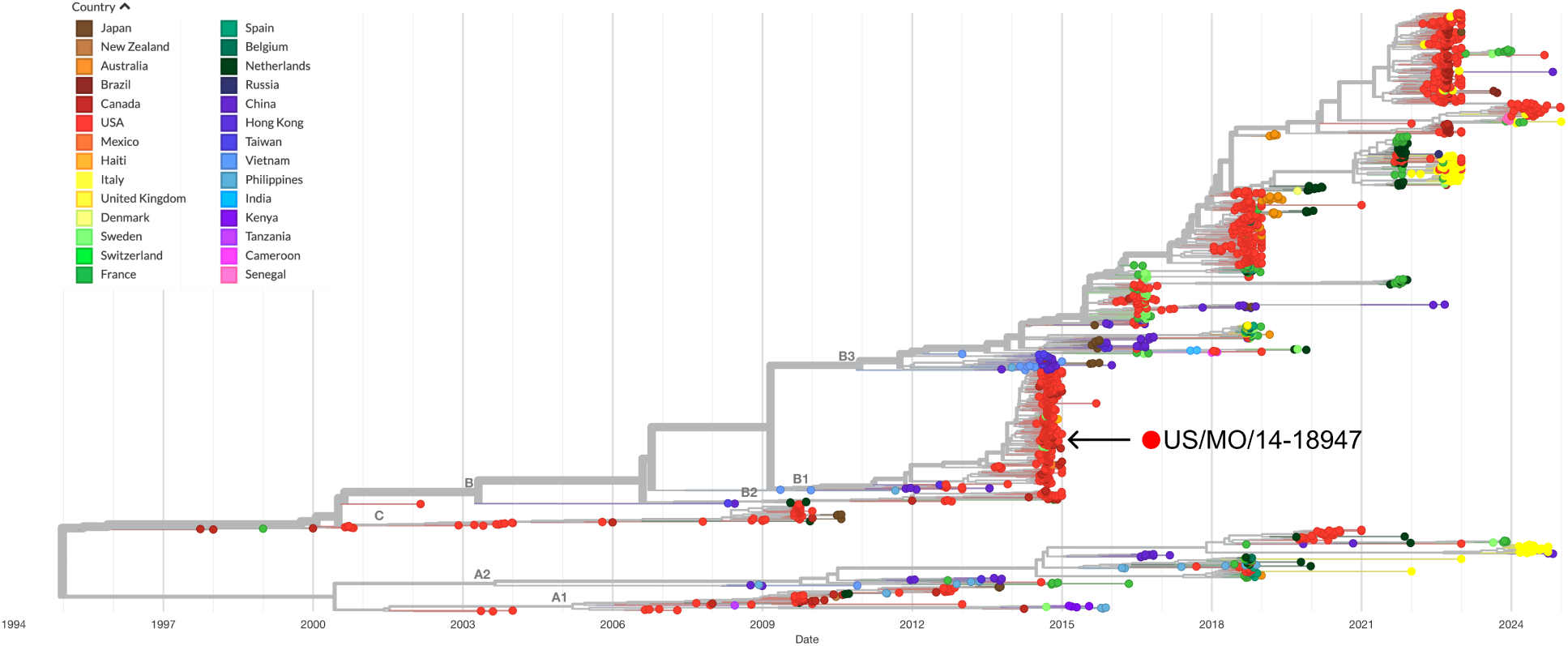
Phylogenetic tree depicting the evolutionary history of EV-D68 for the period 1995-2023, The location of the US/MO/47 strain discovered during the 2014 outbreak of EV-D68 in the US is highlighted. Interactive view generated by NextStrain (https://nextstrain.org/enterovirus/d68/genome)^21,22^

EV-D68 was associated with an outbreak of severe respiratory illness in the United States in 2014.^14,15^ Clinical presentations during the outbreak were typically mild to severe respiratory disease, however, a minority of patients (usually those with näıve or compromised immune systems) developed serious neurological complications, such as acute flaccid paralysis and cranial nerve dysfunction.^16^ There is a growing body of evidence linking EV-D68 infection with these polio-like neurological symptoms.^17–19^ Furthermore, a 2018 study,^20^ compared the infectivity of historical strains of EV-D68 with contemporary strains and found that only the more recently discovered US/MO/47 strain (Figure 1) was able to replicate in the SH-SY5Y neuronal cell line, suggesting a changing risk of neurological sequelae after EV-D68 infection. Enterovirus infections have also been linked to the onset of islet cell autoimmunity (Type 1 Diabetes), clinical trials involving anti-EV compounds are ongoing to investigate this link.^23–25^

Though EV-D68 is a non-enveloped virus, it is cold-adapted and pH-sensitive,^31,32^ putting the primary site of viral replication and transmission in the upper respiratory tract. This heightens the risk of rapid airborne transmission and wider spread of the virus. Despite the risk posed by pandemic EV-D68, there are currently no approved drugs, vaccines or monoclonal antibodies available. The closest point to an approved drug capable of treating EV-D68 was pleconaril (**1**, Figure 2A), an orally bioavailable small molecule that shows broad spectrum picornavirus antiviral activity *in-vitro*.^33–35^ Pleconaril was developed as a treatment for the common cold (primarily human rhinoviruses - hRVs), but FDA approval was rejected in 2002 due to a combination of: lack of efficacy, safety concerns and other factors.^36^

**Figure 2:**
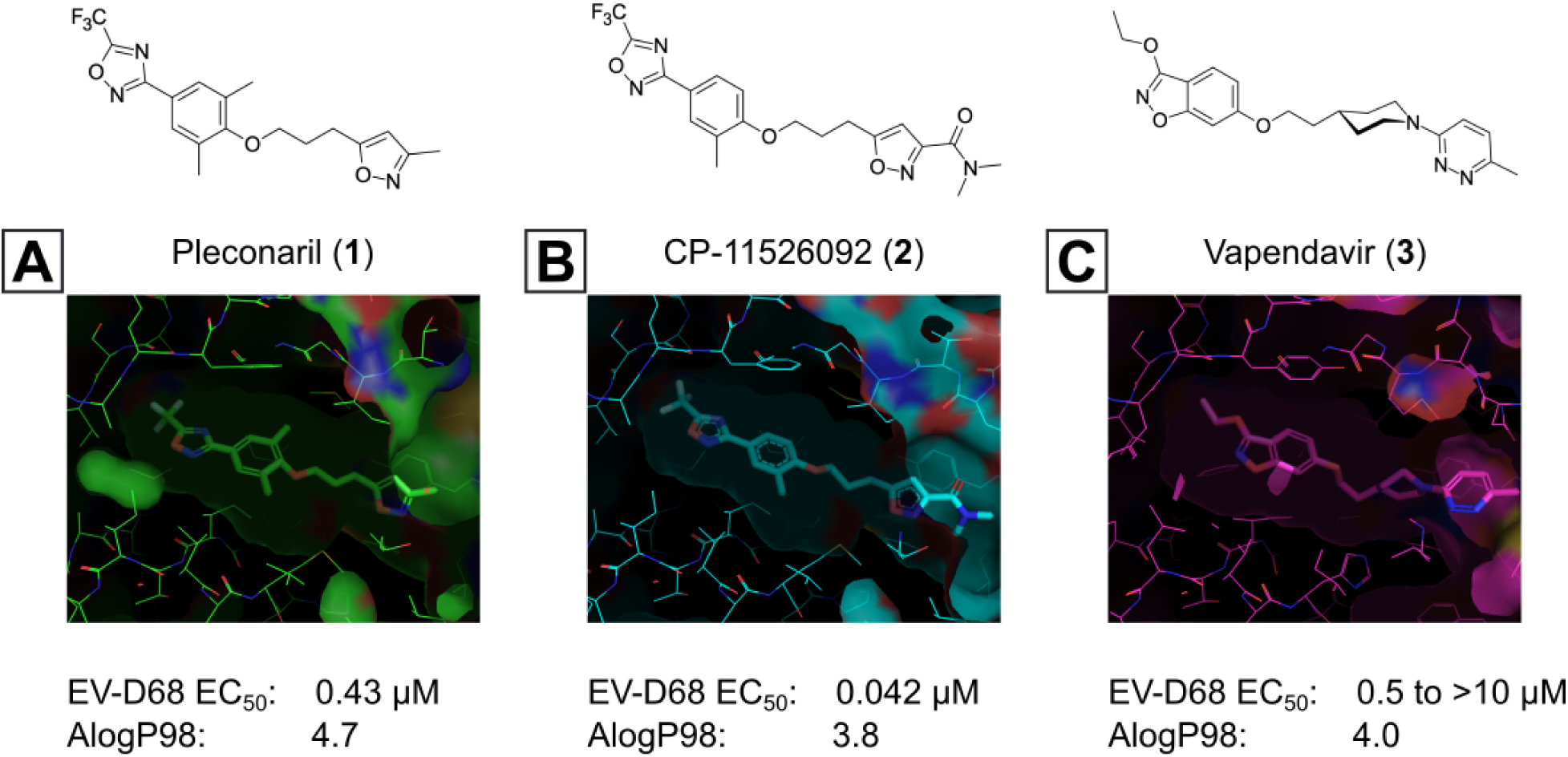
Known capsid binding antiviral compounds. Published EC_50_ values against EV-D68 in-vitro for (A) pleconaril,,^26^ (B) CP-11526092^27^ and (C) vapendavir.^28,29^ Inset structures visualised using PyMOL from left to right: pleconaril (green) bound to the EV-D68 capsid (pdb ID: 4WM7), 11526092 (cyan) bound to the EV-D68 capsid (pdb ID: 7TAF), vapendavir (magenta) bound to the hRV-A2 capsid (pdb ID: 3VDD) AlogP98 = calculated logP. ^30^

The mode of action (MoA) of pleconaril (Figure 3) is mimicry of the natural lipid pocket factor that sits in the hydrophobic cavity of Viral Protein 1 (VP1, one of 4 protomers that make up the icosahedral viral capsid: VP1-4).^26,37,38^ This pocket factor regulates capsid stability and is released when in the acidic environment of the endosome after cell entry, leading to viral uncoating and injection of viral RNA into the host cell. Pleconaril replaces the pocket factor and prevents one or both of: (1) virus-cell attachment and (2) viral uncoating. The cavity of VP1 lies at the bottom of a cleft in the viral capsid, sometimes referred to as the “canyon” (Figure 3B), ^39–41^ which is thought to selectively recognise cellular receptors such as the Intercellular Adhesion Molecule (ICAM-5 and ICAM-1),^42–45^ the human Scavenger Receptor class B member 2 (hSCARB2)^46^ and sialic acid receptors^47–49^ while resisting immune system surveillance. Other molecular targets investigated for anti-EV-D68 drug discovery include key viral proteases, such as 2A, 2C, 3C proteases and the RNA-dependant RNA polymerase (RdRp). All these targets and approaches to anti EV-D68 drug discovery have been reviewed extensively in recent years.^50–53^

**Figure 3:**
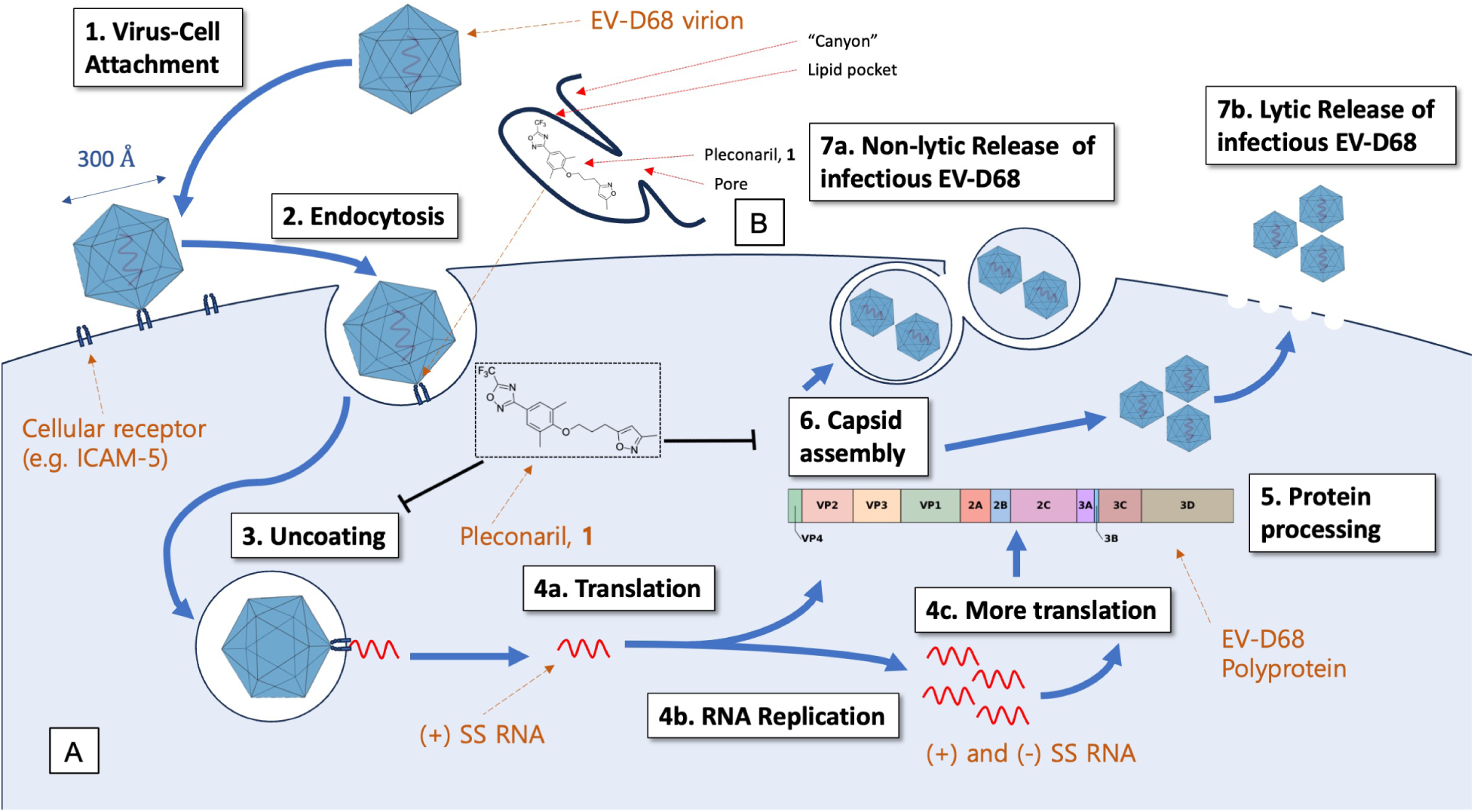
(A) Representation of the EV-D68 replication cycle. (B) Cartoon depicting the topology of the hydrophobic lipid pocket of VP1 with regard to the bound pose of pleconaril, 1 and other capsid-binding antivirals.

Other capsid binders analogous to pleconaril have been investigated recently, such as vapendavir (**3**, Figure 2), which was also developed as a hRV treatment but was similarly halted due to efficacy concerns.^54,55^ The *in-vitro* antiviral activity of vapendavir against EV-D68 is ambiguous, with one source showing activity^29^ and another showing no activity against four strains of EV-D68.^28^ Since the discovery of pleconaril, there have been numerous structure-activity relationship (SAR) studies against NPEV of concern.^27,56–65^

For repurposing pleconaril towards the treatment of emerging NPEV, mutations in the VP1 pocket identified as potentially causing pleconaril resistance should be addressed.^66,67^ Makarov and coworkers have carried out systematic work in recent years,^27,58,61–63,68^ testing close analogues of pleconaril against both pleconaril-sensitive and pleconaril-resistant NPEV strains. Intriguing insights into the SAR of this series have been revealed, including a new ligand-bound cryo-EM structure of the EV-D68 capsid,^27^ resulting in the discovery of the next generation antiviral compound: CP-11526092 (**2**, Figure 2). While representing a step-improvement in broad spectrum NPEV activity from pleconaril, **2** is still highly lipophilic and poorly soluble. The physicochemical property profiles of **1** and **2** are not conducive to an orally bioavailable candidate drug without the need for lipid formulations and special drug delivery systems. This introduces another known risk with the pleconaril series: that of cytochrome P450 (CYP) enzyme induction after repeated dosing. This can lead to drug-drug interactions that are unacceptable for a community use antiviral drug. CYP induction is a well-known side effect of **1**,^69^ and is still present in **2** according to *in-vitro* data.^27^ Neither **1** nor **2** are agonists of the human pregnane-X receptor (PXR), ^27^ which is a common cause of this phenomenon. Further work is required to determine the root cause of CYP induction and what *in-vitro* data are clinically relevant.

The mechanisms of CYP induction are complex and hard to predict due to overlap of nuclear hormone receptor signalling pathways and other factors.^70,71^ However, we speculate that the shared property elements of **1** and **2**, including high lipophilicity (logP), high numbers of rotational bonds (nRotB), high plasma protein binding (PPB) and relatively slow metabolism combine to give a unique CYP induction risk profile. Data describing the CYP induction status of **3** are not in the public domain, to our knowledge.

Thus, our approach to the continued development of the pleconaril series of compounds as EV-D68 antivirals was to find the minimal pharmacophoric elements of antiviral activity and introduce structural novelty to diverge significantly from the established property space (Figure 4). This will be presented here in two parts, firstly, we will present our attempts at rationally editing the present series including combining structural features of literature compounds where structurally and synthetically plausible. Secondly, we will share our results in wider library design and antiviral screening.

**Figure 4:**
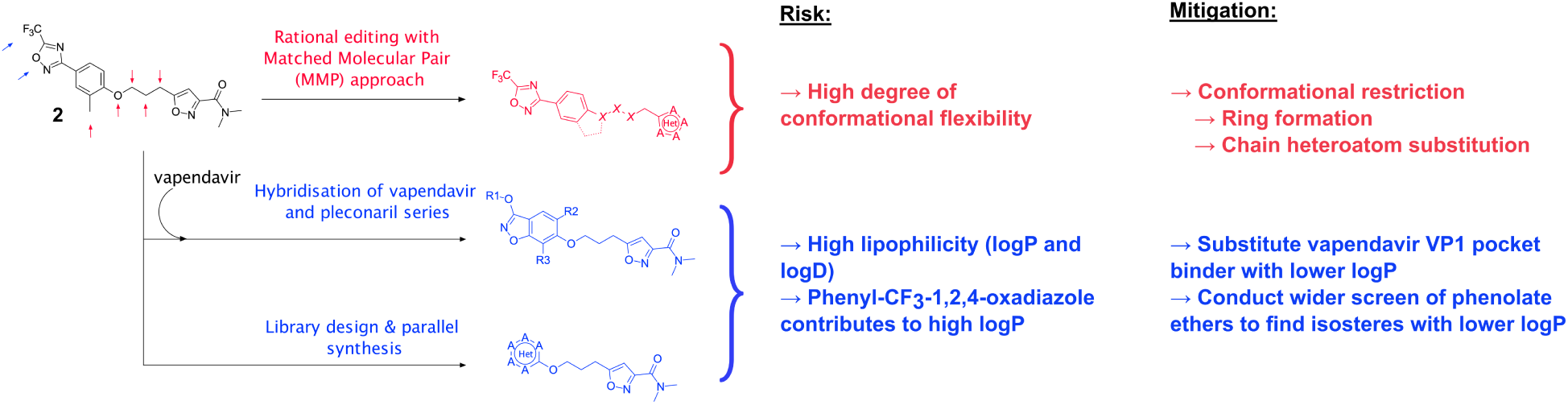
Summary of the medicinal chemistry strategy adopted in this work.

### Synthetic Chemistry

The majority of analogues presented here were made according to Scheme 1, whereby the alcohol **5** was prepared from **4**, and reacted with selected (hetero)phenols under Mitsunobu conditions.^72,73^ The conditions found proved to be generally applicable and were sufficient to rapidly produce analogues **6** in both parallel and singleton fashion.

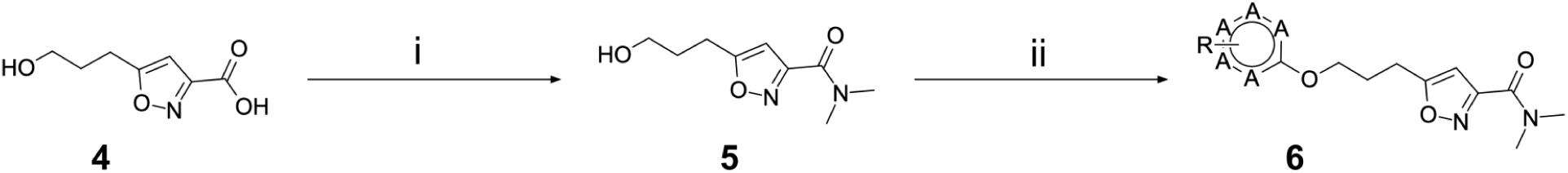

Scheme 1: Reagents and conditions: (i) dimethylamine, HATU, DIPEA, MeCN, RT, 12h (ii) phenol, PPh_3_ or PBu_3_, DEAD or DIAD or ADDP, THF, RT 16h

In contrast, the preparation of ether analogues **10** (Scheme 2) of 2-methyl-4-[5-(trifluoromethyl)-1,2,4-oxadiazol-3-yl]phenol proved to be less generally applicable, primarily due to the chemical sensitivity of the CF_3_-oxadiazole ring towards basic workup procedures. It is for this reason that cyclised analogues **14** (Scheme 3) were prepared according to Scheme 3, where the CF_3_-oxadiazole was constructed in the very last step from the corresponding amidoxime and TFAA.

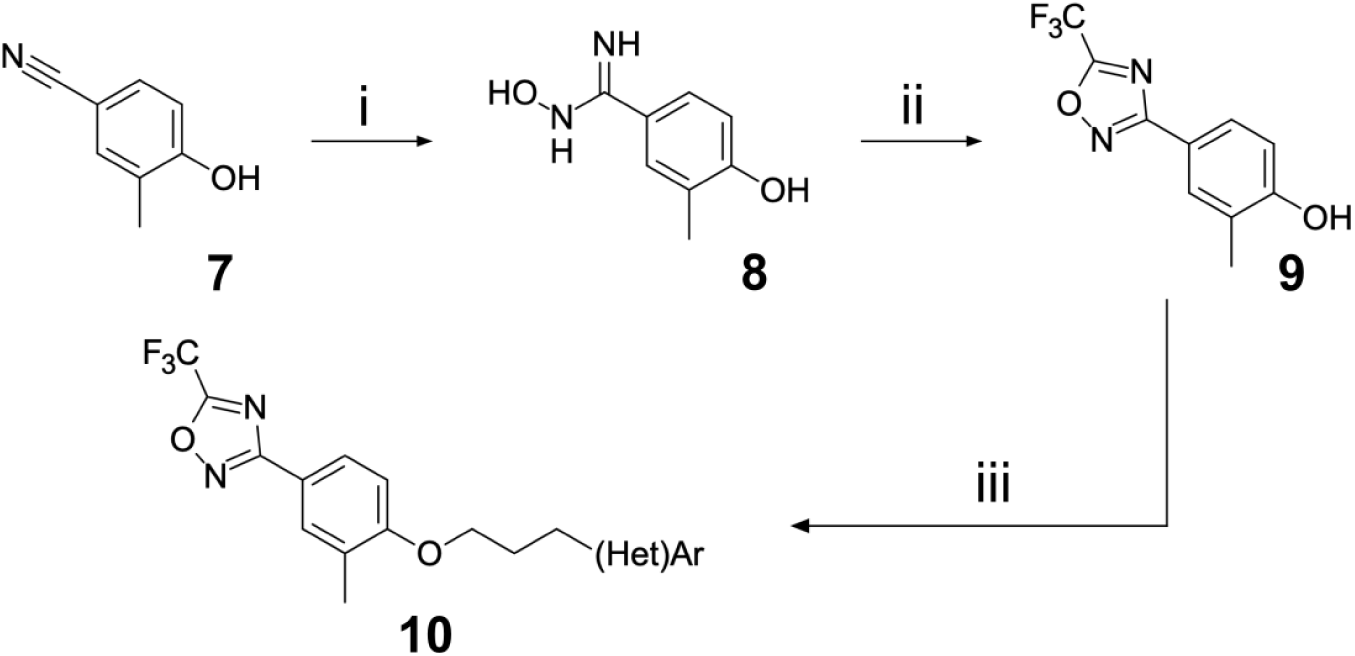

Scheme 2: Reagents and conditions: (i) hydroxylamine hydrochloride, K_2_CO_3_, EtOH (absolute), reflux, 16h (ii) TFAA, pyridine, 80 °C, 2h (iii) 3-(hetero)arylpropyl alcohol, PPh_3_, DEAD, THF, RT, 12h

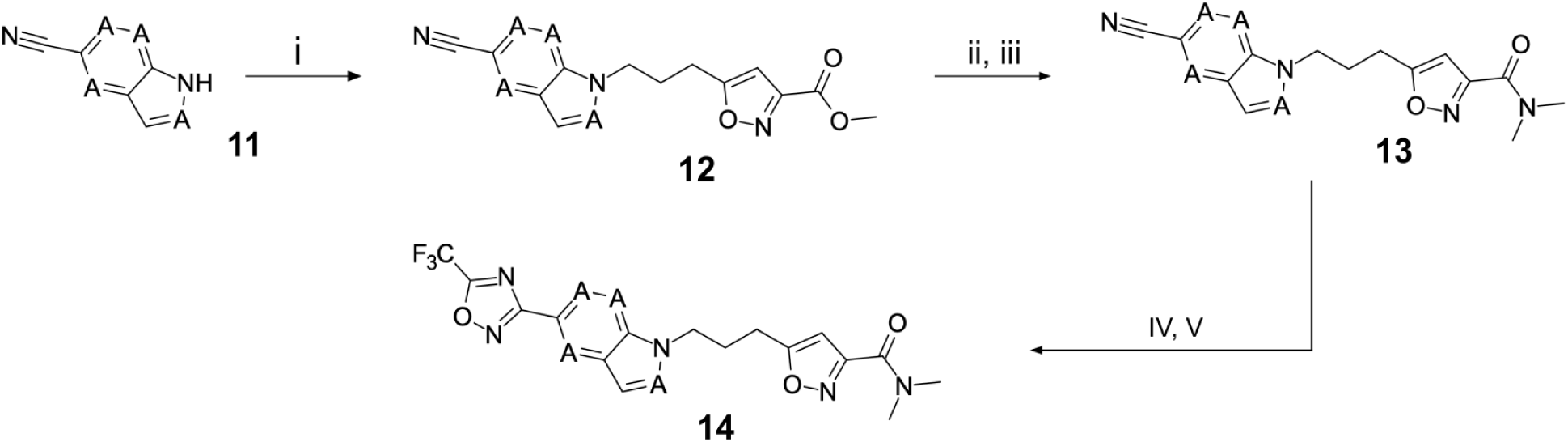

Scheme 3: : Reagents and conditions: (i) methyl 5-(3-bromopropyl)-1,2-oxazole-3-carboxylate, NaH, DMF, 0 to 50 °C, 16 h (ii) LiOH, THF/H_2_O, RT, 16h (iii) dimethylamine, HATU, DIPEA, RT, 16h (iv) hydroxylamine hydrochloride, NaHCO_3_, EtOH, 70 °C, 16h (v) TFAA, THF, RT, 16h

Regarding the preparation of vapendavir-pleconaril hybrid compounds **19** (Scheme 4), a general route that proved successful can be seen in Scheme 4. 2,4-difluoro benzoic acids **15** proved to be very useful double electrophiles in both the synthesis of benzisoxazoles **17** and the final phenolate ester compounds **19**. Other schemes to obtain similar products were attempted, such as in Scheme 5, where 3-chloro-6-fluoro-1,2-benzoxazole **20** was reacted sequentially with the appropriate alcohol under S_N_Ar conditions. Interestingly, this scheme resulted in regioisomeric products **21a** and **21b** (isolated in 1:1 ratio), upon testing, it was found that an example of the originally undesired isomer **21b** (see compound **51**, Table 5) was also active against EV-D68.

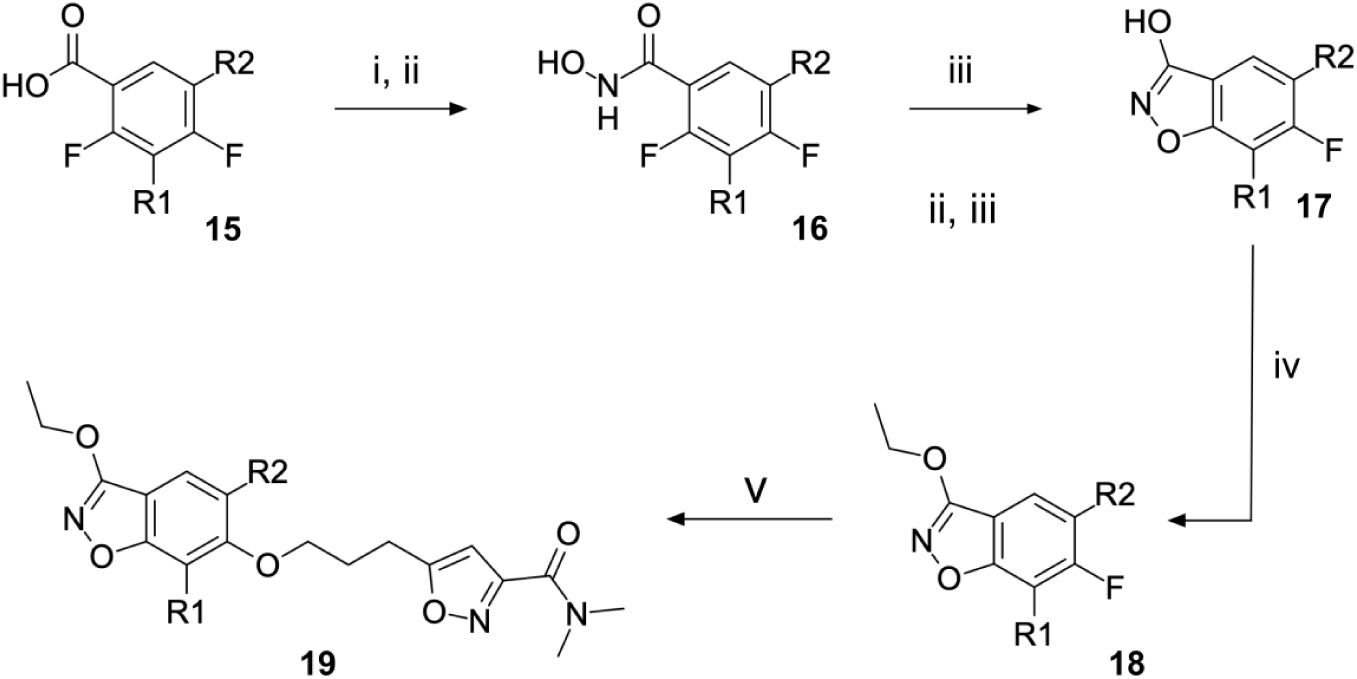

Scheme 4: Reagents and conditions: (i) hydroxylamine hydrochloride, oxalyl chloride (ii) K_2_CO_3_, EtOAc/H_2_O, 0 °C to RT, 16 h (iii) KOH, ^n^BuOH, 120 °C, 4 h (iv) ethyl iodide, Ag_2_O, CHCl_3_, reflux, 16 h (v) 5-(3-hydroxypropyl)-N,N-dimethyl-1,2-oxazole-3-carboxamide, NaH, DMF, 100 °C, 16h

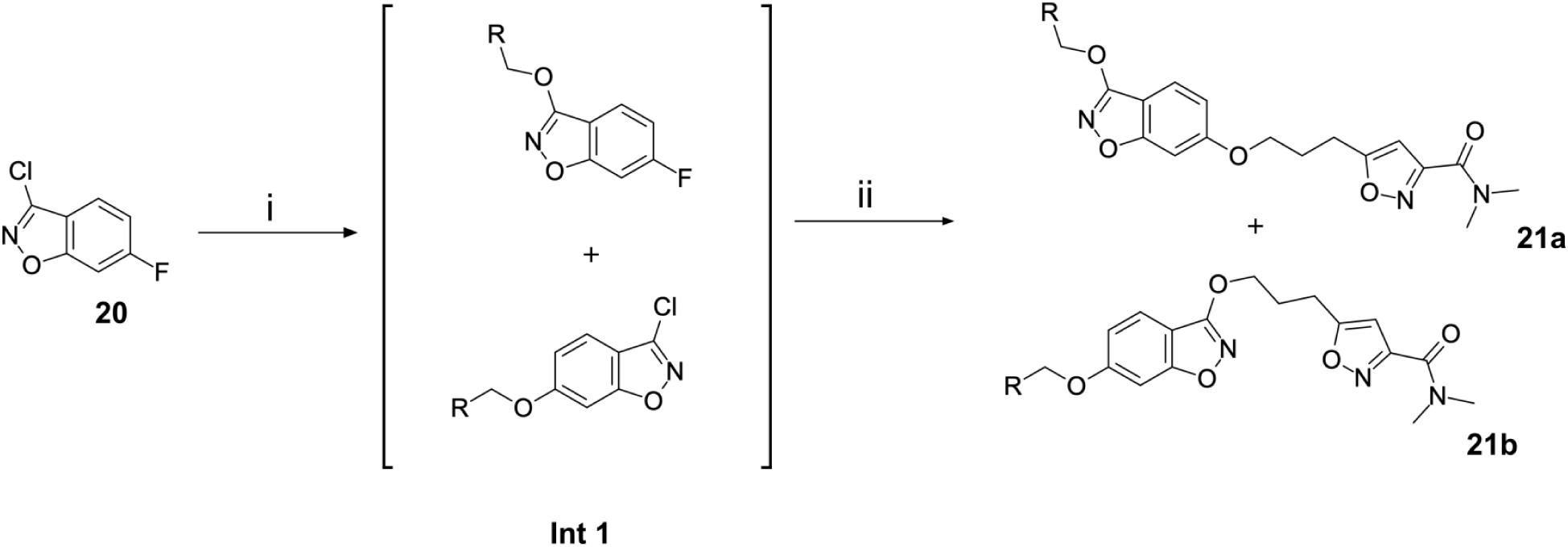

Scheme 5: Reagents and conditions: (i) aliphatic primary alcohol, NaH, DMF, 0 °C to RT, 16 h (ii) 5-(3-hydroxypropyl)-N,N-dimethyl-1,2-oxazole-3-carboxamide, NaH, DMF, 0 °C to RT, 16h

## Results and discussion

### *In-Vitro* Antiviral and ADME Characterisation of Known EV Capsid-Binding Antivirals

Our first round of testing was on the key clinical and preclinical capsid-binding anti-EV compounds found in the literature, on which we would base our studies. We initially screened these compounds in our EV-D68 antiviral assay and a suite of ADME tests, including determination of: logD, kinetic aqueous solubility (KSOL), human and mouse microsomal stability (HLM and MLM, respectively) and MDCK-MDR1 A-*>*B permeability. Our results for **1**, **2** and **3** can be seen in Table 1 (for a full table of data including additional literature capsid binders, see **Table S1** in the supporting information). Obtaining antiviral EC_50_ values against EV-D68 confirmed the sub-*µ*M potency of compounds **1**-**3**, with **1** having the highest potency at *<*29 nM. Close analogue of pleconaril: **2** and the clinical antiviral **3** were similarly potent in our hands, with EC_50_ values of 43 nM and 170 nM, respectively.

**Table 1:**
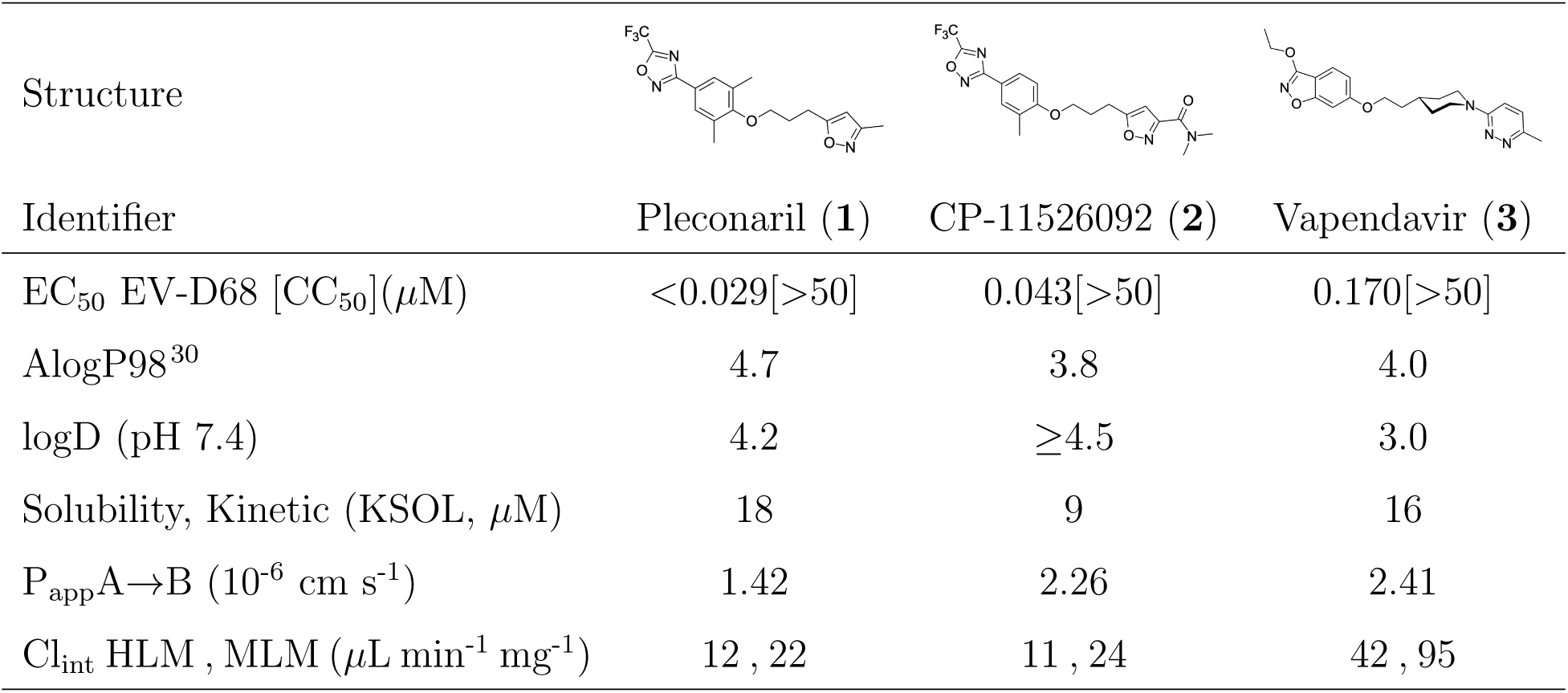
Measured EV-D68 antiviral activity (RD cells) and *in-vitro* ADME profile of relevant known capsid binders from the literature. EC_50_ and CC_50_ are reported as the geometric mean. P_app_ = Apparent Permeability (MDCKII-MDR1 cell system), Cl_int_ = Intrinsic Clearance, HLM = Human Liver Microsomes, MLM = Mouse Liver Microsomes. See the Supporting Information for further details.

| Structure |  |  |  |
| --- | --- | --- | --- |
| Identifier | Pleconaril (**1**) | CP-11526092 (**2**) | Vapendavir (**3**) |
| $EC_{50}$ EV-D68 [$CC_{50}$](μM) | <0.029[>50] | 0.043[>50] | 0.170[>50] |
| AlogP98<sup>30</sup> | 4.7 | 3.8 | 4.0 |
| logD (pH 7.4) | 4.2 | $\geq 4.5$ | 3.0 |
| Solubility, Kinetic (KSOL, μM) | 18 | 9 | 16 |
| $P_{app}A \rightarrow B$ ($10^{-6}$ cm s<sup>-1</sup>) | 1.42 | 2.26 | 2.41 |
| $Cl_{int}$ HLM , MLM (μL min<sup>-1</sup> mg<sup>-1</sup>) | 12 , 22 | 11 , 24 | 42 , 95 |

The ADME properties of the published compounds were measured (Table 1), these compounds generally exhibit: high lipophilicity, high permeability and low aqueous solubility with varying levels of intrinsic microsomal clearance. **1** is known to have useful PK properties,^35,74^ although the oral bioavailability (%F) is heavily dependent on whether the patient is in a fed or fasted state. The high %F of **1** is in spite of its high lipophilicity (measured logD = 4.20), due to a number of factors including: high tissue penetration^75^ and a “global protective” effect from P450 metabolism provided by the CF_3_ group attached to the oxadiazole ring.^76^

### Structural Editing of Known EV Capsid-Binding Antivirals

Building on the extensive SAR studies by Makarov and coworkers cited above, we began our investigations by modifying compound **2**. Moving the oxygen atom of the ether linker along the chain by one (**22**, Table 2) and two (**23**) atoms resulted in a drop in logD and a jump in the measured kinetic aqueous solubility. Both **22** and **23** displayed slightly reduced anti-EV-D68 potency compared to **2**, with EC_50_ values of 115 nM and 216 nM respectively. Moving to an ethanolamine linker (**24**) resulted in much diminished antiviral activity. Al-though it wasn’t experimentally determined, the cell permeability was expected to be lower than **2** due to the introduction of the aniline NH into the chain. ^77^ To probe the effects of conformational restriction by scaffold-hopping, (aza)indole-linked compounds **25-30** (Table 3) were prepared according to Scheme 3. The intermediate nitrile **25** was tested and found to have an EC_50_ of 2.87 *µ*M, significantly less potent against EV-D68 than **2**, but appreciably active. The first indole oxadiazole derivative **26** was found to be very potent against EV-D68, with an EC_50_ value of 17 nM. While the high antiviral potency of **26** against EV-D68 was a promising result, the measured ADME properties suggested limited improvement from **2** in terms of lipophilicity (logD = ≥4.5) and aqueous solubility (KSOL = 8 *µ*M).

**Table 2:**
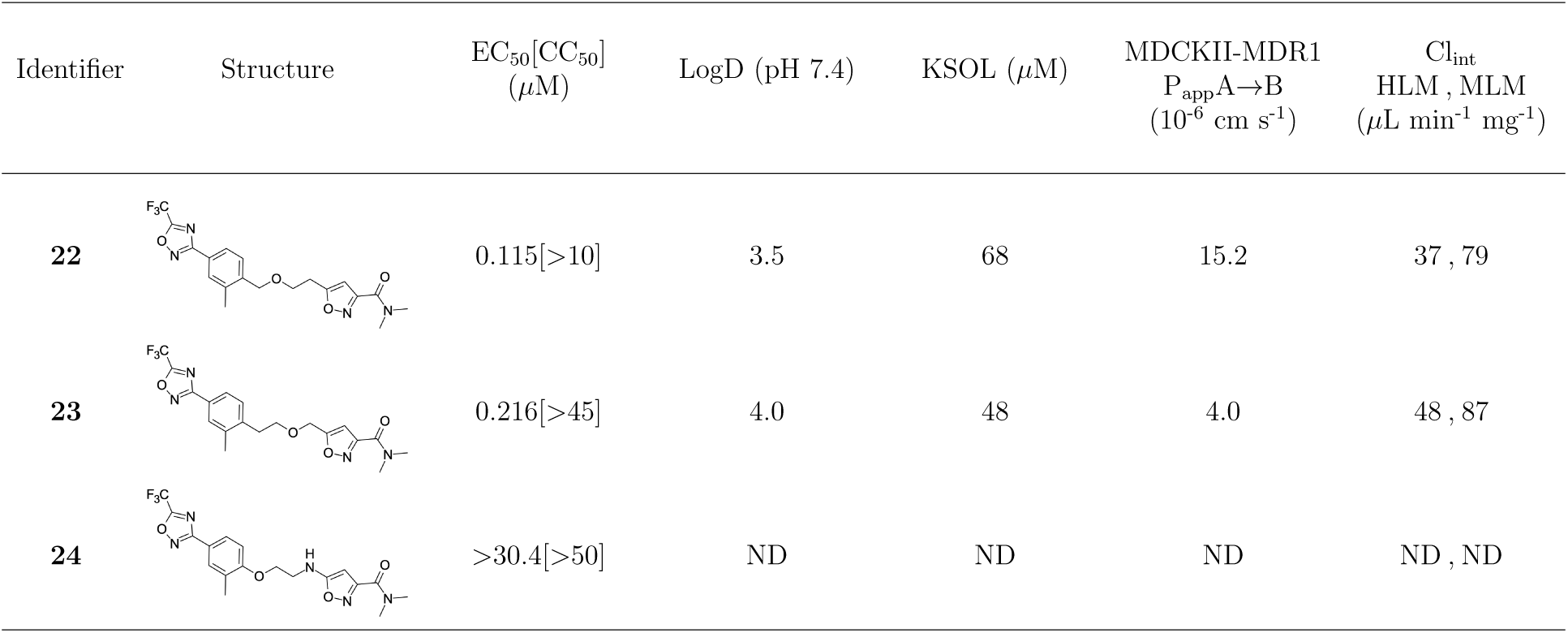
Measured EV-D68 antiviral activity (RD cells) and and *in-vitro* ADME profile of linker-variant compounds from Compound **2**. See the Supporting Information for further details.

| Identifier | Structure | EC<sub>50</sub>[CC<sub>50</sub>]<br>(μM) | LogD (pH 7.4) | KSOL (μM) | MDCKII-MDR1<br>P<sub>app</sub>A→B<br>(10<sup>-6</sup> cm s<sup>-1</sup>) | Cl<sub>int</sub><br>HLM, MLM<br>(μL min<sup>-1</sup> mg<sup>-1</sup>) |
| --- | --- | --- | --- | --- | --- | --- |
| **22** |  | 0.115[>10] | 3.5 | 68 | 15.2 | 37, 79 |
| **23** |  | 0.216[>45] | 4.0 | 48 | 4.0 | 48, 87 |
| **24** |  | >30.4[>50] | ND | ND | ND | ND, ND |

**Table 3:**
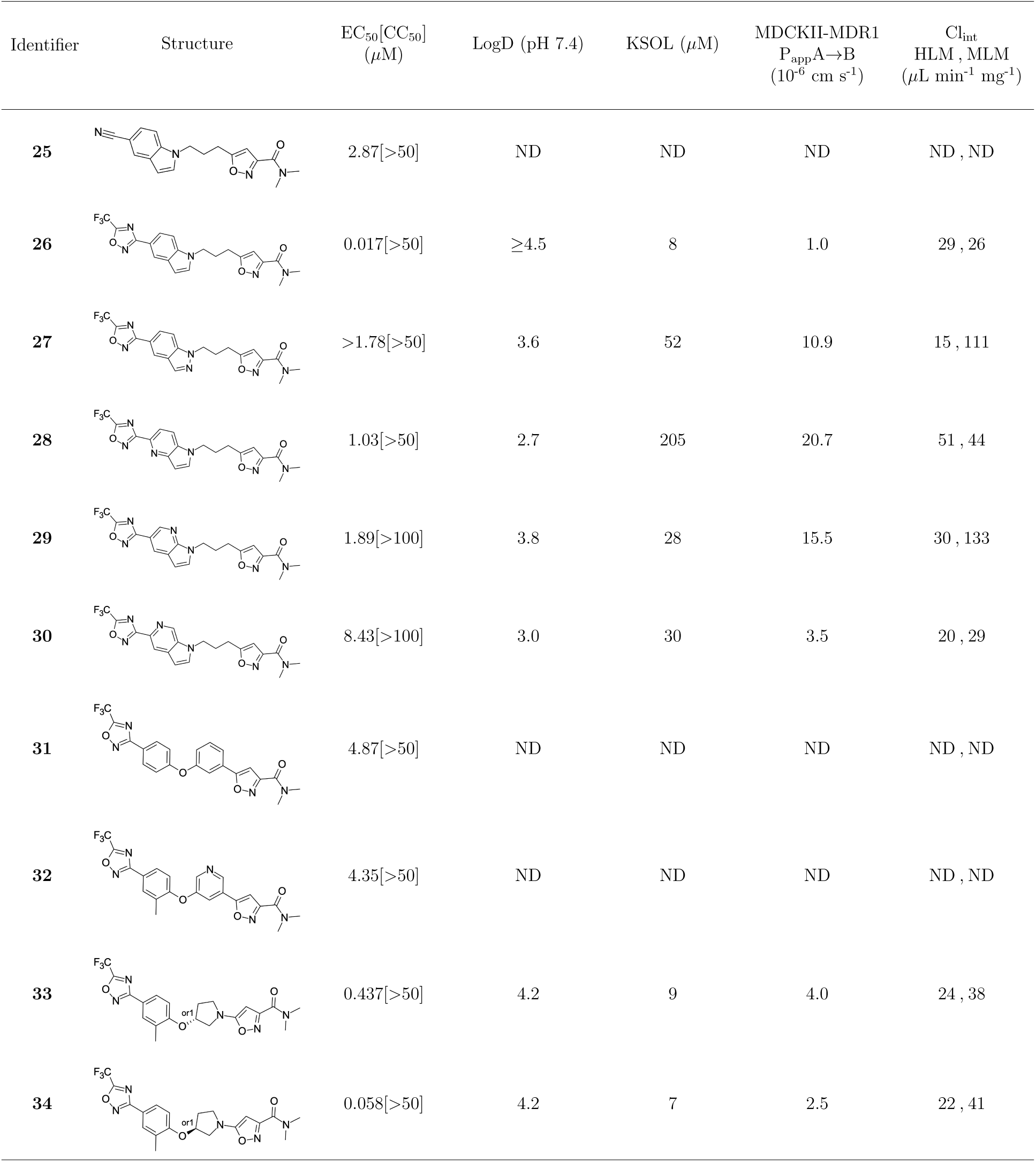
Measured EV-D68 antiviral activity (RD cells) and and *in-vitro* ADME profile of scaffold-hopped designs. See the Supporting Information for further details.

| Identifier | Structure | EC<sub>50</sub>[CC<sub>50</sub>]<br>(μM) | LogD (pH 7.4) | KSOL (μM) | MDCKII-MDR1<br>P<sub>app</sub>A→B<br>(10<sup>-6</sup> cm s<sup>-1</sup>) | Cl<sub>int</sub><br>HLM, MLM<br>(μL min<sup>-1</sup> mg<sup>-1</sup>) |
| --- | --- | --- | --- | --- | --- | --- |
| **25** |  | 2.87[>50] | ND | ND | ND | ND, ND |
| **26** |  | 0.017[>50] | ≥4.5 | 8 | 1.0 | 29, 26 |
| **27** |  | >1.78[>50] | 3.6 | 52 | 10.9 | 15, 111 |
| **28** |  | 1.03[>50] | 2.7 | 205 | 20.7 | 51, 44 |
| **29** |  | 1.89[>100] | 3.8 | 28 | 15.5 | 30, 133 |
| **30** |  | 8.43[>100] | 3.0 | 30 | 3.5 | 20, 29 |
| **31** |  | 4.87[>50] | ND | ND | ND | ND, ND |
| **32** |  | 4.35[>50] | ND | ND | ND | ND, ND |
| **33** |  | 0.437[>50] | 4.2 | 9 | 4.0 | 24, 38 |
| **34** |  | 0.058[>50] | 4.2 | 7 | 2.5 | 22, 41 |

Thus, nitrogen substituted analogues **27**-**30** were also prepared and profiled similarly. All of these aza-substituted analogues dropped the measured logD and increased KSOL relative to **2**, as expected, but this was found to be at the cost of EV-D68 antiviral potency. Benzene and pyridine linker replacements for the central propyl chain were investigated in **31** and **32**, respectively. Neither of these provided potency gains, nor were expected to offer any benefit in terms of ADME properties so were deprioritised from further testing. Finally, pyrollidine-linked enantiomers **33** and **34** were prepared. Both enantiomers were found to have submicromolar EC_50_ values, but **34** was 10-fold more potent than **33**. Intrigued by the departure from closer analogues of **2** described above: represented by these rigid pyrollidine-linked compounds, we further profiled **33** and **34** in ADME assays. Compared to **2**, they maintained a similar profile, with high lipophilicity (logD = 4.2), low kinetic solubility (*<*10 *µ*M), high permeability and moderate to low microsomal clearance. While the properties of **34** as a whole were not sufficient for progression, 2 rotatable bonds were eliminated from **2** through this change to a pyrollidine linker, making the compound significantly more conformationally rigid. This novel chemotype is a highly attractive core for further derivatisation and property optimisation.

We then turned our attention to the VP1 pore-binding (Figure 3B) region, by preparing analogues of **2** according to Scheme 2, using readily available (heteroaryl)propyl alcohol building blocks (Table 4). 1,3,4-oxadiazoles **35**-**38** were all found to be active against EV-D68. Unsubstituted azole **35** was potent, with an EC_50_ of 41 nM while also offering a lower logD and higher solubility than **2**. Methyl substituted analogue **36** lost some EV-D68 potency (EC_50_ = 932 nM), but further improved the solubility (KSOL = 78 *µ*M). **36**, however, was found to have moderate microsomal clearance for both human and mouse (MLM/HLM Cl_int_ = 75/17 *µ*L min^-1^ mg^-1^ protein). Interestingly, **39** (the triazole matched pair of **36**) had much lower microsomal clearance values (MLM/HLM Cl_int_ = 15/*<*10 *µ*L min^-1^ mg^-1^ protein) and was significantly more potent against EV-D68, but it did display a lower solubility (KSOL = 18 *µ*M) compared to **36**. Fluoroalkyl substitution of the oxadiazole was explored, first with difluoromethyl derivative**, 37**, which showed high antiviral potency but very low solubility (KSOL = 6 *µ*M). Trifluoromethyl derivative **38** was found to be much less potent (EV-D68 EC_50_ = 39.9 *µ*M) and was not characterised further.

**Table 4:**
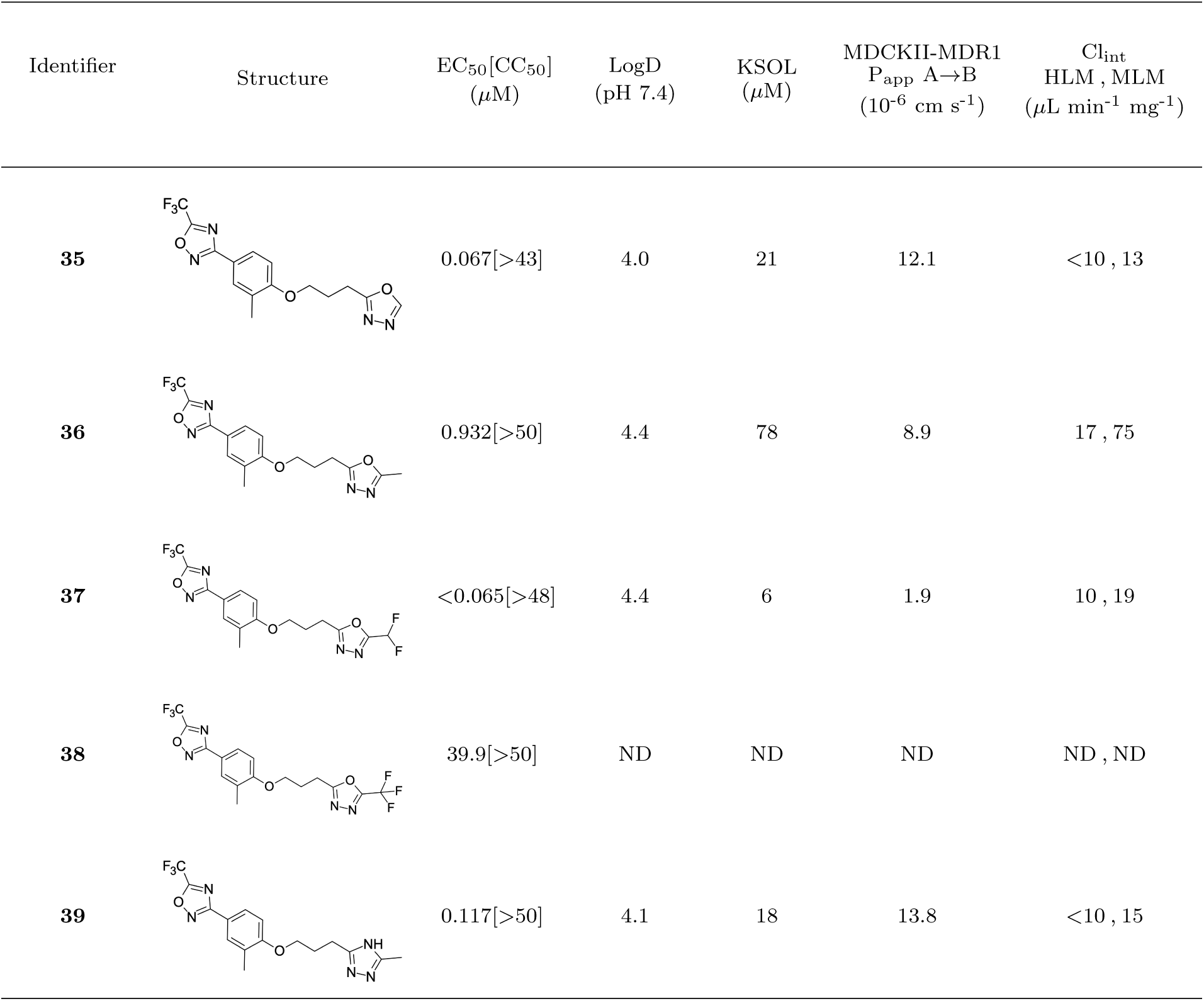
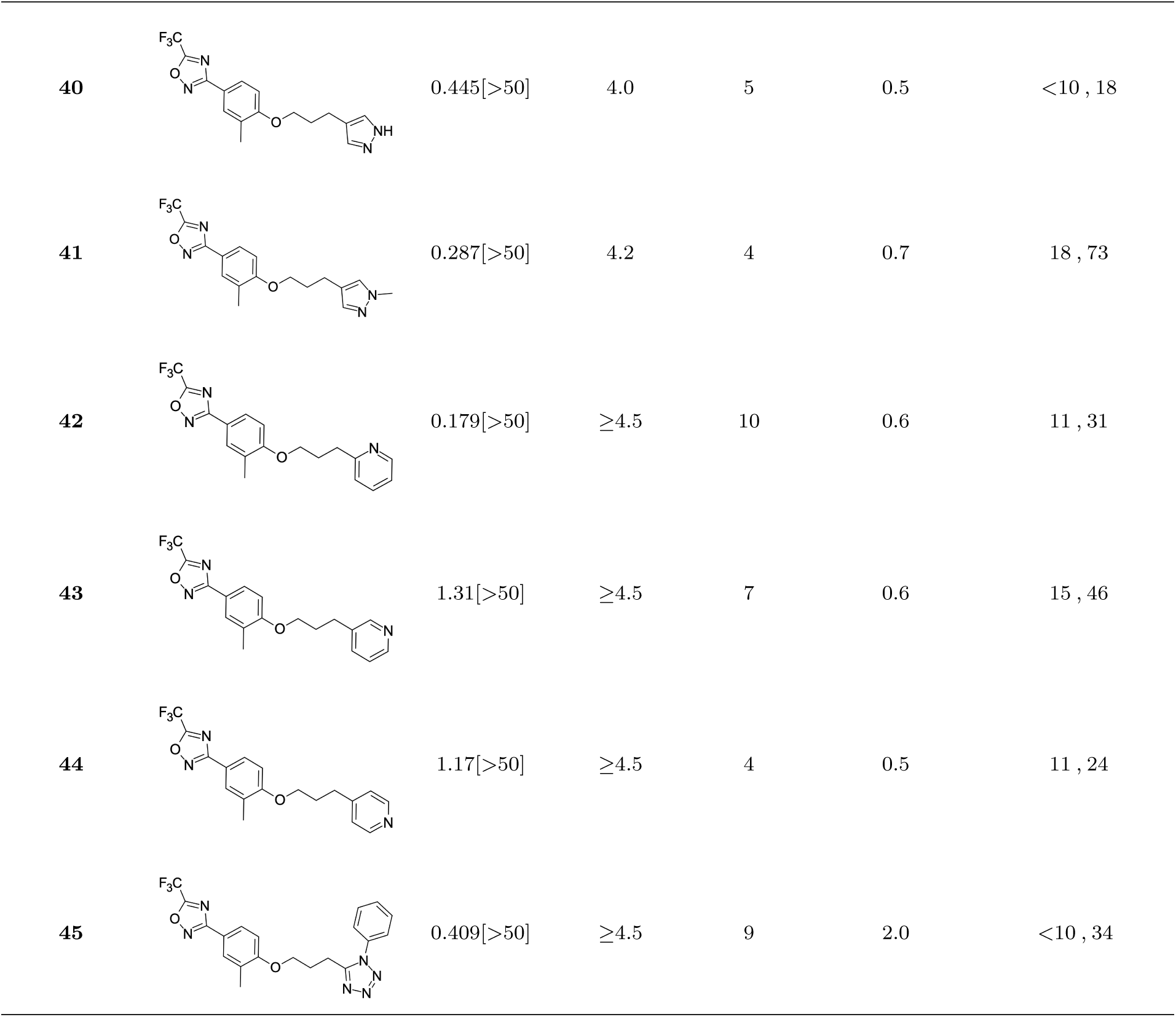
Measured EV-D68 antiviral activity (RD cells) and and *in-vitro* ADME profile of compounds with alternative capsid VP1 pore-binding groups. See the Supporting Information for further details.

| Identifier | Structure | $EC_{50}[CC_{50}]$<br>(μM) | LogD<br>(pH 7.4) | KSOL<br>(μM) | MDCKII-MDR1<br>$P_{app}$ A→B<br>($10^{-6}$ $cm$ $s^{-1}$) | $Cl_{int}$<br>HLM, MLM<br>(μL $min^{-1}$ $mg^{-1}$) |
| --- | --- | --- | --- | --- | --- | --- |
| **35** |  | 0.067[>43] | 4.0 | 21 | 12.1 | <10, 13 |
| **36** |  | 0.932[>50] | 4.4 | 78 | 8.9 | 17, 75 |
| **37** |  | <0.065[>48] | 4.4 | 6 | 1.9 | 10, 19 |
| **38** |  | 39.9[>50] | ND | ND | ND | ND, ND |
| **39** |  | 0.117[>50] | 4.1 | 18 | 13.8 | <10, 15 |

Table 4: (Continued)
| Identifier | Structure | EC<sub>50</sub>[CC<sub>50</sub>]<br>(μM) | LogD<br>(pH 7.4) | KSOL<br>(μM) | MDCKII-MDR1<br>P<sub>app</sub> A→B<br>(10<sup>-6</sup> cm s<sup>-1</sup>) | Cl<sub>int</sub><br>HLM, MLM<br>(μL min<sup>-1</sup> mg<sup>-1</sup>) |
| --- | --- | --- | --- | --- | --- | --- |
| 40 |  | 0.445[>50] | 4.0 | 5 | 0.5 | <10, 18 |
| 41 |  | 0.287[>50] | 4.2 | 4 | 0.7 | 18, 73 |
| 42 |  | 0.179[>50] | ≥4.5 | 10 | 0.6 | 11, 31 |
| 43 |  | 1.31[>50] | ≥4.5 | 7 | 0.6 | 15, 46 |
| 44 |  | 1.17[>50] | ≥4.5 | 4 | 0.5 | 11, 24 |
| 45 |  | 0.409[>50] | ≥4.5 | 9 | 2.0 | <10, 34 |

**Table 5:**
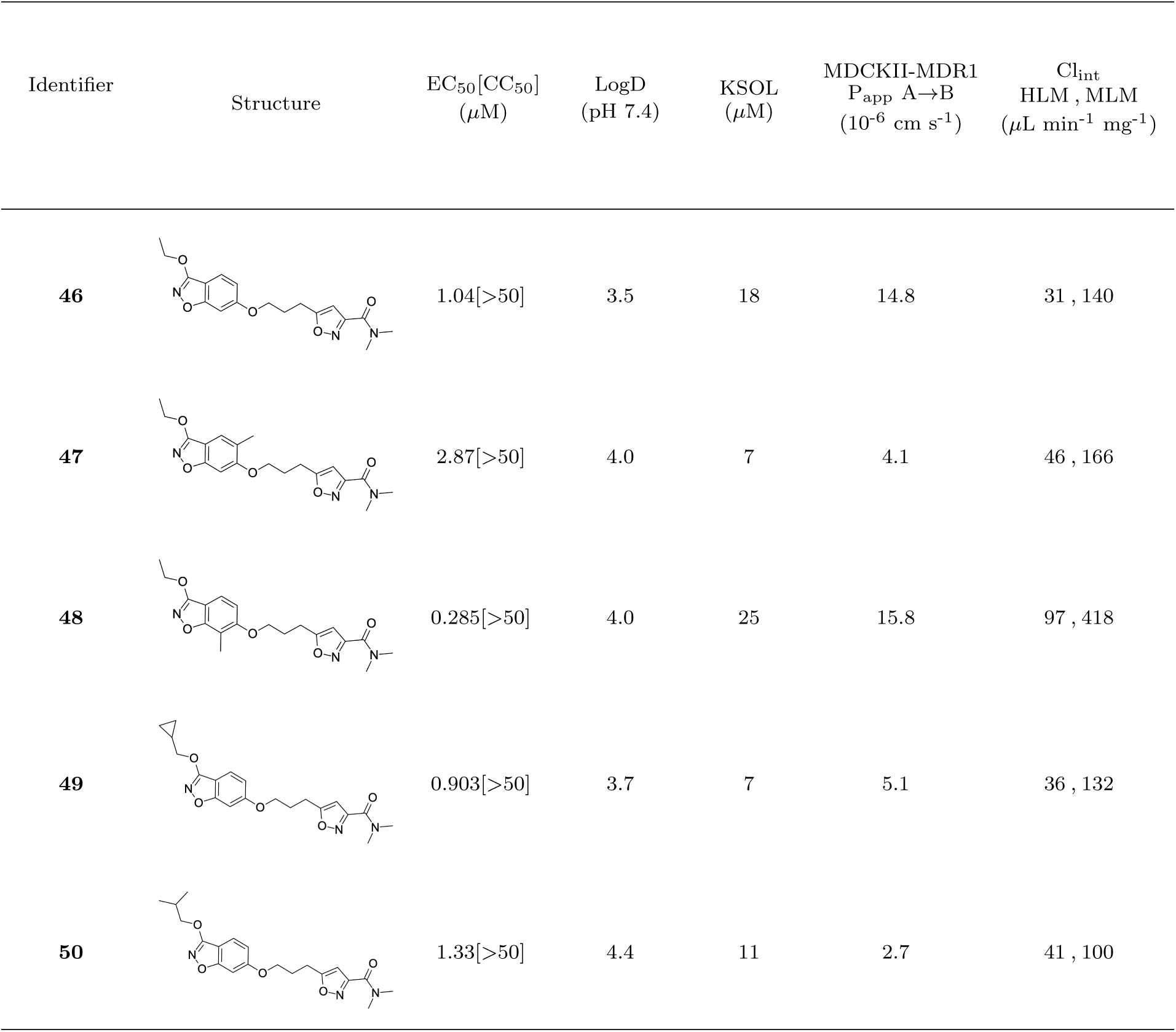
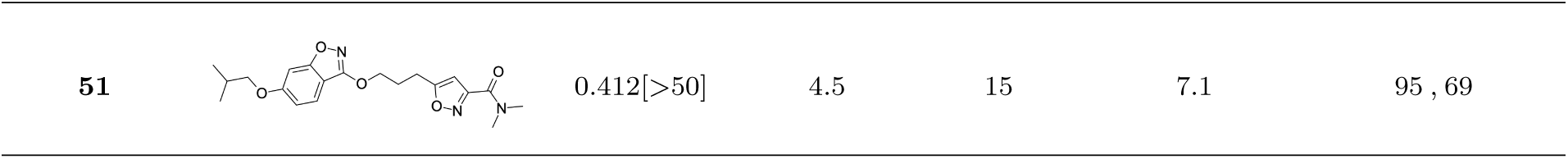
Measured EV-D68 antiviral activity (RD cells) and and *in-vitro* ADME profile of compounds representing structural mergers between compounds **2** and **3**. See the Supporting Information for further details.

| Identifier | Structure | $EC_{50}[CC_{50}]$<br>($\mu M$) | LogD<br>(pH 7.4) | KSOL<br>($\mu M$) | MDCKII-MDR1<br>$P_{app} A \rightarrow B$<br>($10^{-6} cm s^{-1}$) | $Cl_{int}$<br>HLM, MLM<br>($\mu L \min^{-1} mg^{-1}$) |
| --- | --- | --- | --- | --- | --- | --- |
| **46** |  | 1.04[>50] | 3.5 | 18 | 14.8 | 31, 140 |
| **47** |  | 2.87[>50] | 4.0 | 7 | 4.1 | 46, 166 |
| **48** |  | 0.285[>50] | 4.0 | 25 | 15.8 | 97, 418 |
| **49** |  | 0.903[>50] | 3.7 | 7 | 5.1 | 36, 132 |
| **50** |  | 1.33[>50] | 4.4 | 11 | 2.7 | 41, 100 |

Table 5: (Continued)
| Identifier | Structure | EC<sub>50</sub>[CC<sub>50</sub>]<br>(μM) | LogD<br>(pH 7.4) | KSOL<br>(μM) | MDCKII-MDR1<br>P<sub>app</sub> A→B<br>(10<sup>-6</sup> cm s<sup>-1</sup>) | Cl<sub>int</sub><br>HLM, MLM<br>(μL min<sup>-1</sup> mg<sup>-1</sup>) |
| --- | --- | --- | --- | --- | --- | --- |
| **51** |  | 0.412[>50] | 4.5 | 15 | 7.1 | 95, 69 |

Pyrazole (**40**, **41**) and pyridine (**42**-**44**) isomers were found to have useful, but variable levels of EV-D68 potency, these could be considered as useful isosteres for the isoxazole pore-binding groups found in **1** and **2**, for later SAR/SPR combinations. Finally, the phenyl tetrazole compound **45** was found to have useful anti-EV-D68 potency, with an EC_50_ of 55 nM. This tetrazole functionality bears some resemblance to the EV-D68 antiviral R856932 recently discovered by Wang *et al.*^60^ whose capsid-binding MoA was investigated extensively and is expected to bind the pore region of VP1 as we also expect of the tetrazole of **45**.

To explore SAR and SPR mergers between the pleconaril (**1**) and vapendavir (**2**) compound series, novel hybrid compounds **46**-**50** (Table 5) were prepared according to Schemes 4 and 5. The simplest benzisoxazole combination, **46**, was found to have modest potency against EV-D68 (EC_50_ = 1.04 *µ*M), much lower lipophilicity than **2** (logD = 3.5) and moderate human and mouse microsomal clearance HLM/MLM Cl_int_ = 31/140 *µ*L min^-1^ mg^-1^ protein).

Methyl substitution in the 5-position of the benzisoxazole, giving compound **47**, was detrimental to EV-D68 potency and tested ADME properties. Interestingly, 7-methyl analogue **48** was found to be much more potent (EC_50_ = 152 nM) than **47** while also having slightly higher solubility (KSOL = 25 *µ*M). The comparison of potency between isolipophilic compounds **47** and **48** is reminiscent of a “magic methyl” effect,^78,79^ where a significant and non-obvious change in activity is yielded from methyl substitution. With the context of methylation SAR from the work of Makarov et al.^58,63^ and alignment of published capsid structures of **2** (pdb ID = 7TAF) and **3** (pdb ID = 3VDD) (alignment not shown, see **Figures S1 and S2** in the supporting information), we can infer that the optimal location to place the single methyl group on the benzisoxazole would be the 7-position, as confirmed by the activity of **48**. Regrettably, **48** displayed metabolic instability, with HLM and MLM Cl_int_ values of 97 and 418 mL min^-1^ mg^-1^ protein, respectively. Despite having a lower logD than **2**, benzisoxazole compounds **46**, **47** and **48** displayed much lower microsomal stability. This could suggest a structural metabolic vulnerability of the alkoxy benzisoxazole itself, or it could signify a return to a more conventional relationship between lipophilicity and metabolic stability from the anomalous profile of **1** and **2**.

Alternative alkyl chain termini were investigated in compounds **49** (cyclopropyl methyl) and **50** (isobutyl), these compounds were similar in potency to the ethyl parent **46** and didn’t offer any improvement in ADME properties. As outlined in Scheme 5, sequential addition of alcohols to the bifunctional reagent **20** resulted in isomeric structures **21a** and **21b**.

One such example of isomer **21b**, compound **51** (Table 5), was tested and found to have an EV-D68 EC_50_ of just 412 nM. This intriguing result encouraged us to profile **51** further, revealing broadly similar ADME properties to its isomer **50**, lower MLM clearance (MLM Cl_int_ = 69 mL min^-1^ mg^-1^ protein) but interestingly, slightly higher HLM clearance (HLM Cl_int_ = 95 mL min^-1^ mg^-1^ protein).

As high lipophilicity is one of the key risks with most of these compounds, a key metric for ranking was the Lipophilic Ligand Efficiency (LLE), ^80,81^ or in our case of cellular potency measurements: Lipophilic Cellular Efficiency (LipCellE). LipCellE is defined as the compound’s measured pEC_50_ value minus its calculated lipophilicity (AlogP98). Figure 5 shows an overview of EV-D68 cell potency (pEC_50_) for the compounds discussed in the previous section, against their AlogP98 values and coloured by KSOL.

**Figure 5:**
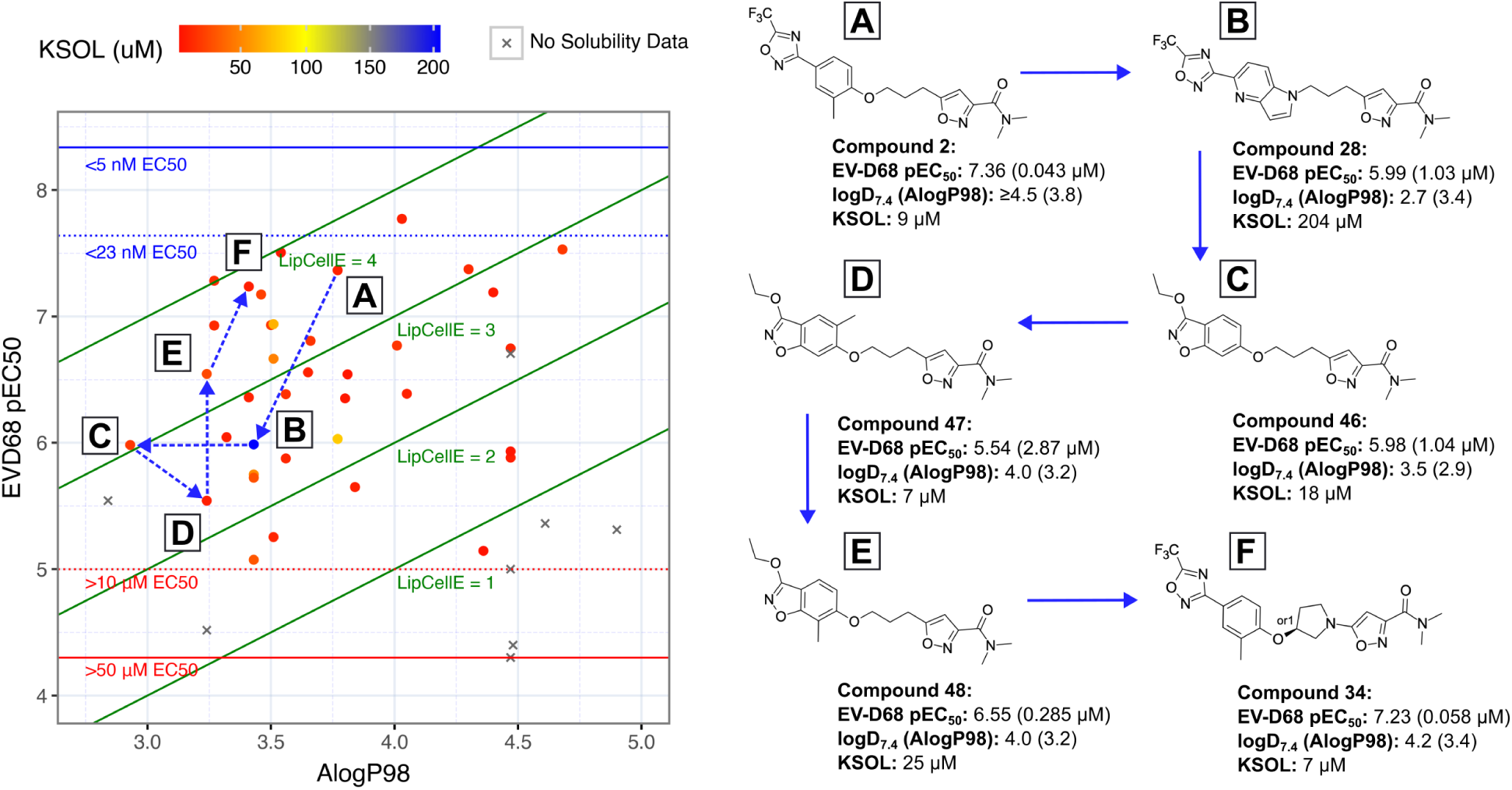
Plot of EV-D68 antiviral potency (pEC50 = -log10(EC50)) vs calculated lipophilicity (AlogP98) for compounds designed via rational editing of known antivirals. Points are coloured based on their measured kinetic solubility values (KSOL / *µ*M). Lipophilic Cellular Efficiency (LipCellE) displayed as normalisation factors of cellular potency against the lipophilicity of the compound (LipCellE = cellular pEC50 - AlogP98)

Figure 5, A shows the position of **2**, highlighting that although it is highly lipophilic, its high potency places the LipCellE at *>*3 indicating that the lipophilic groups on **2** add potency more efficiently than many others in this set (I.e. compounds with LipCellE*<*3). Moving to Figure 5, <u>B</u>, shows us compound **28**, which displays excellent aqueous KSOL and a lower logP than **2**, but the loss in potency is enough to lower the LipCellE value to below 3. Almost equipotent to **28** is vapendavir-hybrid compound **46** (Figure 5, <u>C</u>), with a lower lipophilicity, reinstating the higher LipCellE. 5’-methylation of the benzisoxazole of **46** gives us **47** (Figure 5, <u>D</u>), this was a move that increased the lipophilicity and lowered the potency which is the most direct way to lower LipCellE. In contrast, 7’-methylation in compound **48** (Figure 5, <u>E</u>) provided a 19-fold increase in potency from **47** for the same lipophilicity, providing a drastic increase in LipCellE. Finally, moving to the highly constrained pyrollidine **34** (Figure 5, <u>F</u>) resulted in yet higher potency along with only a slight increase in lipophilicity, further increasing the LipCellE, just above that of **2**.

### Design and Synthesis of Pleconaril Analogue Libraries

During the course of our synthetic work, we discovered an efficient set of Mitsunobu coupling conditions for the production of phenol ethers (Scheme 1). These conditions were robust enough to be employed in parallel synthesis, so we designed a library of compounds based on available phenol building blocks (BBs) aimed at exploring the VP1 pocket more thoroughly. We began this process by characterising a set of ~4000 phenols (Figure 6) with calculated properties, including: 3D shape descriptors (Normalised Principal Moment of Inertia, NPMI1, NPMI2)^82^ and pharmacophoric descriptors (identity and topological bond count from reacting group of pharmacophoric features). A summary of this full characterisation can be seen in Figure 6B, with the NPMI plot showing the 3D shape distribution of the total BB set lying largely on the “Rod-Disk” axis, as expected for a phenol library. The pharmacophoric feature map displayed good coverage of features among the total set, with the exception of acidic groups (“Acid”). This is due to carboxylic acids being filtered to avoid alkylation under Mitsunobu conditions.

**Figure 6:**
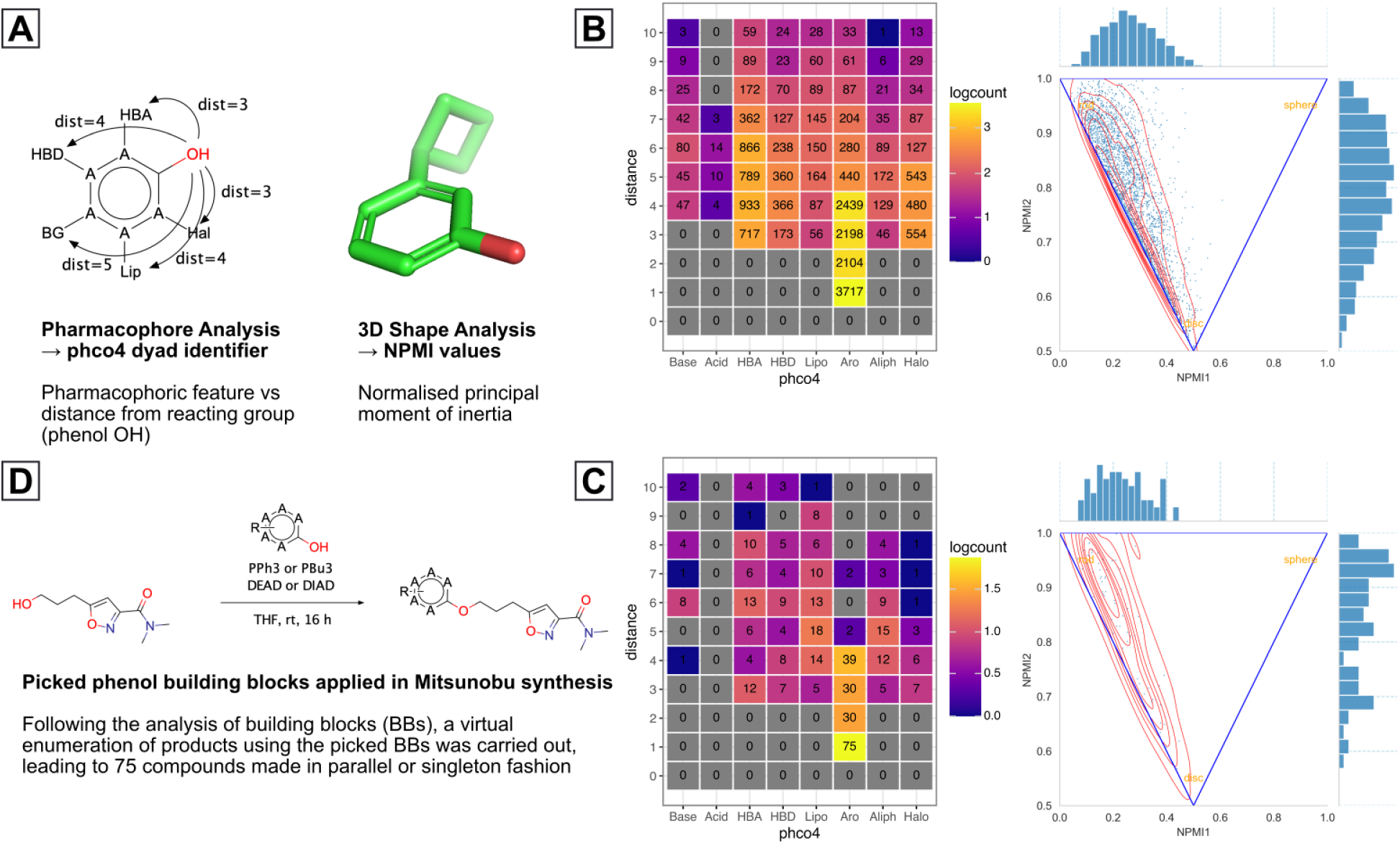
Method for selection of building block (BB) phenols for application in parallel and singleton Mitsunobu reactions, enabling further exploration of the capsid VP1 lipophilic pocket. **A**: Methods through which phenol BB libraries were characterised, **B**(left): Pharmacophore feature heatmap for starting phenol BB library, **B**(right): NPMI^82^ plot for starting phenol BB library, **C**(left): Pharmacophore feature heatmap for selected phenol BBs, **C**(right): NPMI plot for selected phenol BBs, **D**: Scheme for Mitsunobu reactions suitable for parallel synthesis, used to make products. Base = basic group, Acid = Acidic group, HBA = hydrogen bond acceptor, HBD = hydrogen bond donor, Lipo = lipophilic group, Aro = aromatic group, Aliph = aliphatic group, Halo = halogen, phco4 = pharmacophore, NPMI = Normalised principal moment of Inertia

Our phenol picking criteria were centred on whether or not it added diversity in either 3D-shape or pharmacophoric feature coverage (within the confines of chemical space provided by the input phenol BBs), alongside a final ranking based on heavy atom count (HAC) to ensure only the most efficient cluster representatives were taken forward. Out of 123 systematically picked phenols, 80 enumerated ether products were proposed for synthesis, of these, 75 were made and tested. The characteristics of the building blocks associated with these 75 compounds can be seen in Figure 6C. Some gaps in pharmacophoric feature coverage relative to the starting set (Fig. 6A) can be seen, such as “Base 5” and “Aro 6”. This suggests that our selection lacked coverage of basic groups 5 atoms away from the phenol and aromatic groups 6 atoms away. Nevertheless, numerous compounds active against EV-D68 were discovered, a summary of their antiviral pEC_50_ values vs their AlogP98 can be seen in Figure 7.

**Figure 7:**
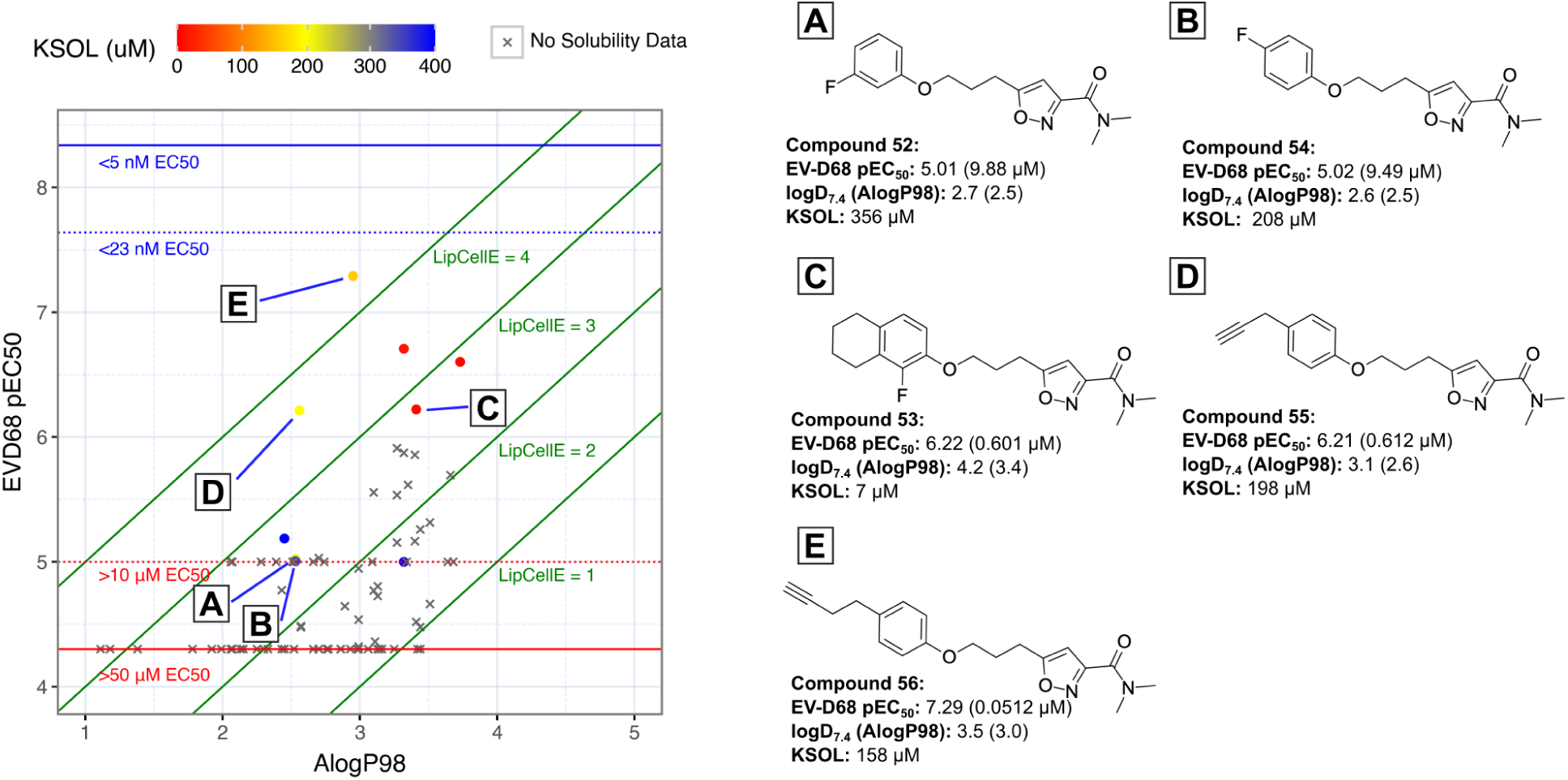
Plot of EV-D68 antiviral potency (pEC50 = -log10(EC50)) vs calculated lipophilicity (AlogP98) for compounds designed via diversity-based library approach. Points are coloured based on their measured kinetic solubility values (KSOL / *µ*M). Lipophilic Cellular Efficiency (LipCellE) displayed as normalisation factors of cellular potency against the lipophilicity of the compound (LipCellE = cellular pEC50 - AlogP98)

The compounds that initially caught our eye as having maintained some EV-D68 antiviral activity, were the fluoroaryls **52**, **53** and **54**. Simple fluorobenzene derivatives **52** and **54** had modest EV-D68 antiviral activity (EC_50_ values of 9.88 *µ*M and 9.49 *µ*M, respectively), but showed reduced lipophilicity and increased KSOL values, indicating that a compromise between potency and lipophilicity could possibly be reached with further development. The more heavily substituted **53** was more potent, but at the expense of higher lipophilicity and lower solubility (logD = 4.2, KSOL = 7 *µ*M). Propargyl derivative **55** was found to have low micromolar potency (EC_50_ = 4.88 *µ*M) and a modest logD value of 3.1, alongside a useful KSOL of 198 *µ*M. Interestingly, the extended homologue of **55**: **56**, was found to have the highest LipCellE out of the entire library and it maintained a lower logD and higher KSOL.

### Assessment of Broad-Spectrum Antiviral Activity Through EV Panel Screening

To test the most promising compounds for broad spectrum anti-enterovirus activity, we profiled **26**, **48** and **56** against a panel of 8 enteroviruses (Table 6).

**Table 6:**
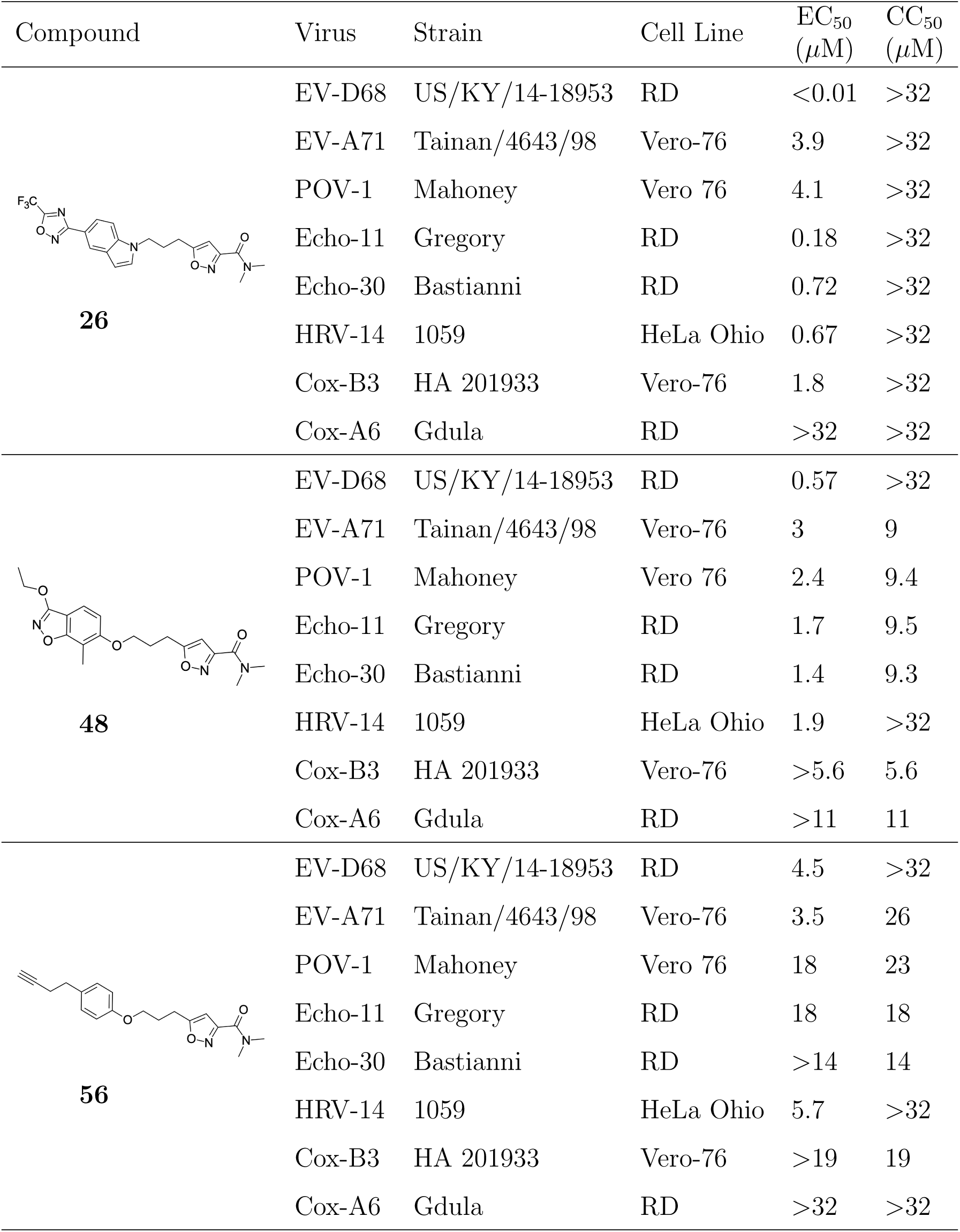
Summary of results from the screening of **26**, **48** and **56** against a panel of 8 enteroviruses. EV = Enterovirus, POV = Poliovirus, Echo = Echovirus, HRV = Human Rhinovirus, Cox = Coxsackievirus. See the supporting information for further details.

| Compound | Virus | Strain | Cell Line | EC<sub>50</sub><br>(μM) | CC<sub>50</sub><br>(μM) |
| --- | --- | --- | --- | --- | --- |
| **26** | EV-D68 | US/KY/14-18953 | RD | <0.01 | >32 |
|  | EV-A71 | Tainan/4643/98 | Vero-76 | 3.9 | >32 |
|  | POV-1 | Mahoney | Vero 76 | 4.1 | >32 |
|  | Echo-11 | Gregory | RD | 0.18 | >32 |
|  | Echo-30 | Bastianni | RD | 0.72 | >32 |
|  | HRV-14 | 1059 | HeLa Ohio | 0.67 | >32 |
|  | Cox-B3 | HA 201933 | Vero-76 | 1.8 | >32 |
|  | Cox-A6 | Gdula | RD | >32 | >32 |
| **48** | EV-D68 | US/KY/14-18953 | RD | 0.57 | >32 |
|  | EV-A71 | Tainan/4643/98 | Vero-76 | 3 | 9 |
|  | POV-1 | Mahoney | Vero 76 | 2.4 | 9.4 |
|  | Echo-11 | Gregory | RD | 1.7 | 9.5 |
|  | Echo-30 | Bastianni | RD | 1.4 | 9.3 |
|  | HRV-14 | 1059 | HeLa Ohio | 1.9 | >32 |
|  | Cox-B3 | HA 201933 | Vero-76 | >5.6 | 5.6 |
|  | Cox-A6 | Gdula | RD | >11 | 11 |
| **56** | EV-D68 | US/KY/14-18953 | RD | 4.5 | >32 |
|  | EV-A71 | Tainan/4643/98 | Vero-76 | 3.5 | 26 |
|  | POV-1 | Mahoney | Vero 76 | 18 | 23 |
|  | Echo-11 | Gregory | RD | 18 | 18 |
|  | Echo-30 | Bastianni | RD | >14 | 14 |
|  | HRV-14 | 1059 | HeLa Ohio | 5.7 | >32 |
|  | Cox-B3 | HA 201933 | Vero-76 | >19 | 19 |
|  | Cox-A6 | Gdula | RD | >32 | >32 |

Compound **26** gave the most promising profile, showing broad spectrum low-micromolar to nanomolar EC_50_ values against 7 out of 8 viral strains tested with CC_50_ values of *>*32 *µ*M across all cell lines. Sub-micromolar EC_50_ values were seen for EV-D68 (US/ KY/14-18953 strain), Echovirus-11 (Echo-11), Echovirus-30 (Echo-30) and Human Rhinovirus-14 (HRV-14). Slightly lower potency was seen for EV-A71 (3.9 *µ*M), Poliovirus-1 (POV-1, 4.1 *µ*M) and Coxsackievirus-B3 (Cox-B3, 1.8 *µ*M). The notable exception to the broad-spectrum activity of **26** was Coxsackievirus-A6 (Cox-A6), for which no antiviral activity was observed.

Moving to the vapendavir hybrid compound **48**, robust activity against EV-D68 (US/KY/14-18953 strain) was observed plus some other low micromolar potencies, however, cytotoxicity was also observed across the cell lines used which complicated the assessment of antiviral activity.

Finally, Mitsunobu library compound **56** displayed low micromolar EC_50_ values against EV-D68 (4.5 *µ*M) and EV-A71 (3.5 *µ*M) and slightly less cytotoxicity throughout the panel than **48**. However, lower antiviral potency for **56** was also observed across the board. This is somewhat unsurprising, given that **56** was discovered as part of the screening campaign against EV-D68 (US/MO/14-18949 strain) (Figure 7) and did not contain as many highly optimised structural features such as those from CP-11526092 and vapendavir in compounds **26** and **48** (respectively).

## Summary and Conclusions

In summary, we have surveyed numerous approaches in the literature towards repurposing analogues of **1** from its original rhinovirus indication. **1** failed to gain FDA approval for its targeted indication, but remains an interesting clinical tool for treating NPEV, which are of potentially greater public health concern. Among known pleconaril analogues, compound **2** was found to have increased potency towards NPEV than pleconaril, however, it resides in a similar property space and still holds the risk of CYP induction, based on *in-vitro* data. The clinical compound **3** occupies a much more favourable property space, but does not offer the NPEV potency of **1** and **2**, according to the literature.

We prepared analogues of the pleconaril and vapendavir series of compounds and tested them against EV-D68 *in-vitro*, as a representative of the NPEV class. We also tested interesting analogues in our tier 1 ADME assays to probe clinical usefulness in the development of orally bioavailable community-use anti-EV-D68 drugs. Compounds were designed by both rationally editing known antivirals in a matched molecular pair approach and by building block-based library generation. Numerous compounds that are significantly differentiated from published examples, in both structure and properties, displayed potent antiviral activity against EV-D68.

We screened 3 compounds against a panel of 8 enteroviruses and identified **26** as a broad-spectrum enterovirus inhibitor. Other compounds **48** and **56** displayed some antiviral activity against NPEV of concern, but also displayed cytotoxicity in a similar concentration window. No antiviral activity was observed against Coxsackievirus-A6 from any of the compounds tested.

In conclusion, the SAR studies undertaken here provide potentially useful data and offer new starting points for those looking to develop potent and orally bioavailable drugs to treat NPEV.

## Acknowledgement

The authors thank Karla Kirkegaard (ORCID: 0000-0001-7628-3770) for engaging discussions.

## Funding

This work was supported by the NIH and NIAID Antiviral Drug Discovery (AViDD) grant (U19AI171399). This work was also supported in part by services provided by the NIAID Preclinical Services Program under Contract No. 75N93019D00021, Task Order No. B11.

The ISMMS BSL-3 Biocontainment Shared Research Resource is supported by subsidies from the ISMMS Dean’s Office and by investigator support through a cost recovery mechanism. Research reported in this publication was supported by NIAID of the NIH under Award Number G20AI174733 (R.A. Albrecht).

## Disclaimers

The content is solely the responsibility of the authors and does not necessarily represent the official views of the National Institutes of Health.

## Disclosures

DLC, EJG and JS are employees of MedChemica Consultancy Ltd. AAL is an employee of PostEra Inc., which has commercial interests in the discovery, development, and commercialisation of therapeutics, including antivirals and results disclosed in this manuscript. YF, AH, YH, RK, MK, OK, DL, IL, ML, VL, AP, MP, AR and AT are employees of Enamine Ltd. BLH, JGJ and HW are employees of Utah State University. RP, JB, RD-T, MEG, RAA and KW are employees of the Icahn School of Medicine at Mount Sinai.

## Supporting Information Available

Experimental procedures and characterisation data for all new compounds, protocols and references for antiviral and ADME assays.

The deposition of all data related to this work into ChEMBL is in progress. In the meantime, data (including raw data and data not presented here) can be made available on reasonable request.

## Notes

https://dx.doi.org/10.17504/protocols.io.yxmvmywybv3p/v1

https://dx.doi.org/10.17504/protocols.io.5qpvo9jddv4o/v1

https://dx.doi.org/10.17504/protocols.io.e6nvw14kdlmk/v1

https://dx.doi.org/10.17504/protocols.io.j8nlk8y41l5r/v1

https://dx.doi.org/10.17504/protocols.io.n2bvjne6ngk5/v1

https://dx.doi.org/10.17504/protocols.io.5qpvokdb9l4o/v1

https://nextstrain.org/enterovirus/d68/genome

### Competing Interest Statement

The authors have declared no competing interest.

### Summary of Updates

No change to ms, correction to supplementary information Scheme S3

