## Supporting Information for "Structure-Activity Relationship Studies Towards Analogues of Pleconaril as Novel Enterovirus-D68 Capsid-Targeting Antivirals"

*https://icahn.mssm.edu/research/global-health*

⊥*The ASAP Discovery Consortium, https://asapdiscovery.org/*

### Contents

**Synthetic Chemistry and Characterisation Data** **S-5**

General Laboratory Conditions . . . . . . . . . . . . . . . . . . . . . . . . . . . . . S-7

Synthesis of Intermediate **9** . . . . . . . . . . . . . . . . . . . . . . . . . . . . . S-9

Synthesis of Compound **2** . . . . . . . . . . . . . . . . . . . . . . . . . . . . . S-10

Synthesis of Compound **22** . . . . . . . . . . . . . . . . . . . . . . . . . . . . . S-11

Synthesis of Compound **23** . . . . . . . . . . . . . . . . . . . . . . . . . . . . . S-14

Synthesis of Compound **24** . . . . . . . . . . . . . . . . . . . . . . . . . . . . . S-17

Synthesis of Compounds **25** and **26** . . . . . . . . . . . . . . . . . . . . . . . . S-20

Synthesis of Compound **27** . . . . . . . . . . . . . . . . . . . . . . . . . . . . . S-24

Synthesis of Compound **28** . . . . . . . . . . . . . . . . . . . . . . . . . . . . . S-28

Synthesis of Compound **29** . . . . . . . . . . . . . . . . . . . . . . . . . . . . . S-28

Synthesis of Compound **30** . . . . . . . . . . . . . . . . . . . . . . . . . . . . . S-29

Synthesis of Compound **31** . . . . . . . . . . . . . . . . . . . . . . . . . . . . . S-29

Synthesis of Compound **32** . . . . . . . . . . . . . . . . . . . . . . . . . . . . . S-33

Synthesis of Compounds **33** and **34** . . . . . . . . . . . . . . . . . . . . . . . . S-37

Synthesis of Compound **35** . . . . . . . . . . . . . . . . . . . . . . . . . . . . . S-40

Synthesis of Compound **36** . . . . . . . . . . . . . . . . . . . . . . . . . . . . . S-42

Synthesis of Compound **37** . . . . . . . . . . . . . . . . . . . . . . . . . . . . . S-42

Synthesis of Compound **38** . . . . . . . . . . . . . . . . . . . . . . . . . . . . . S-44

Synthesis of Compound **39** . . . . . . . . . . . . . . . . . . . . . . . . . . . . . S-46

Synthesis of Compound **40** . . . . . . . . . . . . . . . . . . . . . . . . . . . . . S-48

Synthesis of Compound **41** . . . . . . . . . . . . . . . . . . . . . . . . . . . . . S-49

Synthesis of Compound **42** . . . . . . . . . . . . . . . . . . . . . . . . . . . . . S-50

Synthesis of Compound **43** . . . . . . . . . . . . . . . . . . . . . . . . . . . . . S-50

Synthesis of Compound **44** . . . . . . . . . . . . . . . . . . . . . . . . . . . . . S-51

Synthesis of Compound **45** . . . . . . . . . . . . . . . . . . . . . . . . . . . . . S-52

Synthesis of Compound **46** . . . . . . . . . . . . . . . . . . . . . . . . . . . . . . . S-54

Synthesis of Compound **47** . . . . . . . . . . . . . . . . . . . . . . . . . . . . . . . S-55

Synthesis of Compound **48** . . . . . . . . . . . . . . . . . . . . . . . . . . . . . . . S-58

Synthesis of Compound **49** . . . . . . . . . . . . . . . . . . . . . . . . . . . . . . . S-61

Synthesis of Compounds **50** and **51** . . . . . . . . . . . . . . . . . . . . . . . . . . S-62

**General Procedure A**:

Parallel Synthesis of Phenol Ethers by Mitsunobu Coupling . . . . . . . . . . S-65

Synthesis of Compound **52** . . . . . . . . . . . . . . . . . . . . . . . . . . . . . . . S-65

Synthesis of Compound **53** . . . . . . . . . . . . . . . . . . . . . . . . . . . . . . . S-65

Synthesis of Compound **54** . . . . . . . . . . . . . . . . . . . . . . . . . . . . . . . S-66

Synthesis of Compound **55** . . . . . . . . . . . . . . . . . . . . . . . . . . . . . . . S-66

Synthesis of Compound **56** . . . . . . . . . . . . . . . . . . . . . . . . . . . . . . . S-67

Synthesis of Compound **S-57** (BTA-188) . . . . . . . . . . . . . . . . . . . . . . . . S-68

Synthesis of Compound **S-59** (NLD-22) . . . . . . . . . . . . . . . . . . . . . . . . S-70

Synthesis of Compound **S-60** (OBR-5-340) . . . . . . . . . . . . . . . . . . . . . . S-74

**Methodology for Antiviral and ADME Assays** **S-77**

EV-D68 Antiviral Screening Assay in RD Cells . . . . . . . . . . . . . . . . . . . . S-77

EV Panel Antiviral Screening Assays . . . . . . . . . . . . . . . . . . . . . . . . . . S-78

ADME Assays - LogD . . . . . . . . . . . . . . . . . . . . . . . . . . . . . . . . . . . S-78

ADME Assays - KSOL . . . . . . . . . . . . . . . . . . . . . . . . . . . . . . . . . . . S-78

ADME Assays - MDCK Permeability . . . . . . . . . . . . . . . . . . . . . . . . . . S-79

ADME Assays - Human and Mouse Microsomal Stability . . . . . . . . . . . . . . S-79

**Antiviral and ADME Profiling of Published Antivirals** **S-81**

**Published EV Capsid Structural Data** **S-82**

List of Ligand-Bound Crystal and Cryo-EM Structures of VP1 . . . . . . . . . . . S-82

3D Overlays . . . . . . . . . . . . . . . . . . . . . . . . . . . . . . . . . . . . . . . . S-85

### Synthetic Chemistry and Characterisation Data

#### Abbreviations

| | |
|---|---|
| **ACN** | Acetonitrile |
| **AcOH** | Acetic acid |
| **ADDP** | 1,1-(Azodicarbonyl)dipiperidine |
| **Ar** | Argon |
| **CataCXium A** | Di(1-adamantyl)-n-butyl phosphine |
| **CDI** | 1,1-Carbonyldiimidazole |
| **DCM** | Dichloromethane |
| **DEAD** | Diethyl azodicarboxylate |
| **DIAD** | Diisopropyl azodicarboxylate |
| **DIPEA** | N,N-Diisopropylethylamine |
| **DMAA** | Dimethylacetamide |
| **DMF** | Dimethylformamide |
| **DMSO** | Dimethyl sulfoxide |
| **ESI** | Electrospray ionisation |
| **EtOH** | Ethanol |
| **EtPh** | Ethylbenzene |
| **$Et_2O$** | Diethyl ether |
| **$Et_3N$** | Triethylamine |
| **FCC** | Flash column chromatography |
| **g** | Gram |
| **HATU** | O-(7-azabenzotriazol-1-yl)-N,N,N,N-tetramethyluronium hexafluorophosphate |
| **h** | Hour |
| **$^1H$** | Proton |
| **HCl** | Hydrochloric acid |
| **HPLC** | High performance liquid chromatography |
| **HSQC** | Heteronuclear single quantum correlation |
| **Hz** | Hertz |
| **$K_3PO_4$** | Potassium phosphate tribasic |
| **LCMS** | Liquid chromatography-mass spectrometry |
| **M** | Molar |
| **MeOH** | Methanol |

| | |
|---|---|
| **mg** | Milligram(s) |
| **$MgSO_4$** | Magnesium sulfate |
| **min** | Minute(s) |
| **mL** | Milliliter(s) |
| **$\mu$M** | Microliter(s) |
| **mm** | Millimeter(s) |
| **mmol** | Millimole(s) |
| **$\mu$mol** | Micromole(s) |
| **NaCl** | Sodium chloride |
| **NaH** | NaH |
| **$NaHCO_3$** | Sodium bicarbonate |
| **NaOH** | Sodium hydroxide |
| **$Na_2SO_4$** | Sodium sulfate |
| **$NH_4Cl$** | Ammonium chloride |
| **$NH_4OH$** | Ammonium hydroxide |
| **NOESY** | Nuclear Overhauser effect spectroscopy |
| **PMB** | 4-Methoxybenzyl |
| **PSCBH** | Polymer-supported cyano borohydride |
| **quant** | Quantitative |
| **rt** | Room temperature |
| **RT** | Retention time |
| **SEM-Cl** | 2-(Trimethylsilyl)ethoxymethyl chloride |
| **SFC** | Supercritical fluid chromatography |
| **TFAA** | Trifluoroacetic anhydride |
| **TFA** | Trifluoroacetic acid |
| **THF** | Tetrahydrofuran |
| **TLC** | Thin layer chromatography |
| **UPLC-MS** | Ultra performance liquid chromatography-mass spectrometry |
| % **wt.** | % weight |
| **XPhos Pd G3** | 2-dicyclohexylphosphino-2,4,6-triisopropyl-1,1-biphenyl)[2-(2-amino-1,1-biphenyl)]palladium(II) methanesulfonate |

#### General Laboratory Conditions

**1** (pleconaril) was purchased from MedChemExpress, **3** (vapendavir) and **S-58** (pirodavir) were purchased from AA Blocks Inc., purchased compounds were used as obtained without further purification. All other starting materials were obtained from commercial suppliers and used without further purification unless stated otherwise.

All evaporations were carried out in vacuo with a rotary evaporator. Analytical samples were dried in vacuo (1-5 mmHg) at rt. All references to brine refer to a saturated aqueous solution of NaCl. Unless otherwise indicated, all temperatures are expressed in °C (degrees Centigrade). Most of the reactions were monitored by UPLC-MS (Shimadzu LCMS-2020 Single Quadrupole Liquid Chromatograph Mass Spectrometer or Waters ACQUITY UPLC I-Class PLUS System with Waters SQ Detector 2) or thin-layer chromatography on 0.25mm Merck silica gel plates (60F-254), visualized with UV light. Flash column chromatography was performed on prepacked silica gel cartridges (50 $\mu$M, puriFlash®), Biotage®Sfär KPAmino D (Duo 50 $\mu$M, 28-110 g) and puriFlash®RP-AQ or C18-HP (15 $\mu$M, 120-330 g). Preparative TLC was performed on silica gel plates glass-backed, 1000 $\mu$M, Analtech. Nuclear magnetic resonance (NMR) spectroscopy of samples was carried out on either a Bruker AVANCE DRX 500 or a Varian UNITYplus 400. Proton NMR spectra were recorded at 400 MHz or 500 MHz. 13C NMR spectra were recorded at 100 MHz or 125 MHz. Chemical shifts ($\delta$) are given in parts per million (ppm) and are listed upfield with tetramethylsilane as a reference. Peaks are described as singlets (s), doublets (d), triplets (t), quartets (q), quintets (quint) multiplets (m) and broad (br.). All assignments of NMR spectra were based on 1D NMR data. Raw NMR data (FID) can be provided on request.

#### LCMS Method Details

**Column** Agilent Poroshell 120 SB-C18 4.6x30mm 2.7 $\mu$m

**Column Temperature** 60°C

**Mobile phase** A: water (0.1% formic acid), B: acetonitrile (0.1% formic acid)

**Flow Rate** 3 ml $min^{-1}$

**Gradient** 0.01 min – 1% B, 1.5 min – 100% B, 1.73 min – 100% B

**MS Ionisation Mode** ESI

**MS Scan Range** 83 – 600 m/z or 83 – 1000 m/z

**UV Detection Wavelengths** 215 nm, 254 nm and 280 nm

#### Synthesis of key building block: 2-methyl-4-[5-(trifluoromethyl)-1,2,4-oxadiazol-3-yl]phenol (Compound 9)

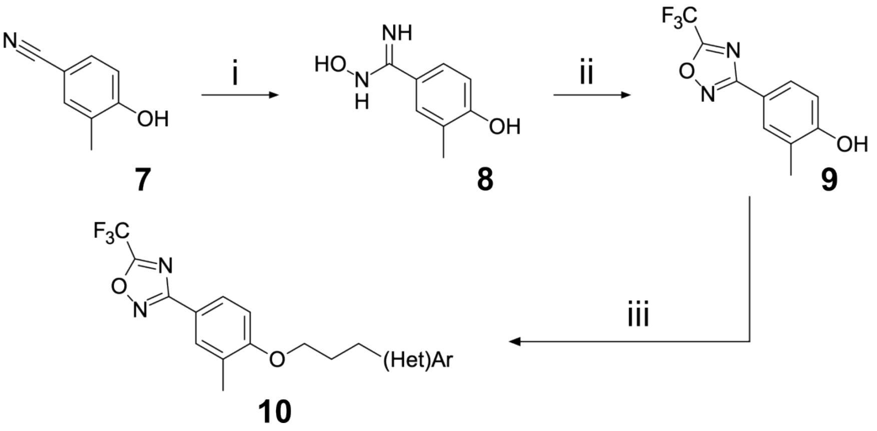

Scheme S1: Reagents and conditions: (i) hydroxylamine hydrochloride, $K_2CO_3$, EtOH (absolute), reflux, 16h (ii) TFAA, pyridine, 80 °C, 2h (iii) 3-(hetero)arylpropyl alcohol, $PPh_3$, DEAD, THF, RT, 12h

#### Step i: Data for N',4-dihydroxy-3-methyl-benzamidine (Compound 8)

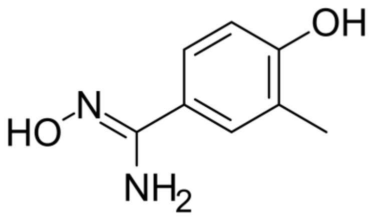

$K_2CO_3$ (51.9 g, 376 mmol) and hydroxylamine hydrochloride (26.1 g, 376 mmol) were added to a stirred solution of 4-hydroxy-3-methyl-benzonitrile (10.0 g, 75.1 mmol) in EtOH (284 mL) at room temperature. The resulting mixture was stirred at reflux overnight. The reaction mixture was filtered hot. The filter cake was washed with hot ethanol (2×100 mL). The combined filtrates were concentrated under reduced pressure to afford N',4-dihydroxy-3-methyl-benzamidine (16 g, 72.21 mmol, 96.15% yield, 75% purity) as a brown solid which was used in the next step without further purification. LCMS(ESI): $[M+H]^+$ m/z: calcd 167.08; found 167.2.

#### Step ii: Data for 2-methyl-4-[5-(trifluoromethyl)-1,2,4-oxadiazol-3-yl]phenol (Compound 9)

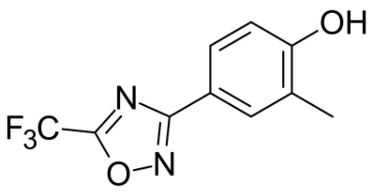


TFAA (30.3 g, 144 mmol, 20 mL) was added dropwise to a stirred solution of N',4-dihydroxy-3-methyl-benzamidine (16.0 g, 72.2 mmol) in pyridine (201 mL) at 85 °C. The resulting mixture was stirred at 85 °C for 1 hr. The reaction mixture was cooled to room temperature and concentrated under reduced pressure. The residue was partitioned between EtOAc (300 mL) and water (100 mL). The organic layer was separated, washed with water (3×100 mL) and brine (100 mL). The combined organic layers were dried over anhydrous sodium sulfate and concentrated under reduced pressure. The residue was subjected to flash column chromatography ($SiO_2$, Hex-EtOAc 4:1) to afford 2-methyl-4-[5-(trifluoromethyl)-1,2,4-oxadiazol-3-yl]phenol (9.50 g, 38.9 mmol, 53.9% yield) as a light-yellow solid. $^1$H NMR (500 MHz, $CDCl_3$) $\delta_H$ 2.34 (s, 3H), 5.26 (s, 1H), 6.91 (d, 1H), 7.82 (d, 1H), 7.92 (s, 1H). LCMS(ESI): $[M+H]^+$ m/z: calcd 245.06; found 245.0.

#### Synthesis of N,N-dimethyl-5-[3-[2-methyl-4-[5-(trifluoromethyl)-1,2,4-oxadiazol-3-yl]phenoxy]propyl]isoxazole-3-carboxamide (Compound 2)

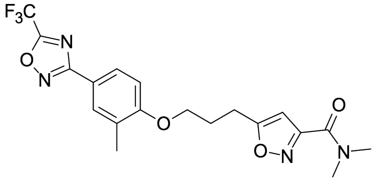


5-(3-Hydroxypropyl)-N,N-dimethyl-isoxazole-3-carboxamide (40.6 mg, 205 $\mu$mol) was mixed with 2-methyl-4-[5-(trifluoromethyl)-1,2,4-oxadiazol-3-yl]phenol (50.0 mg, 205 $\mu$mol) and $PPh_3$ (80.6 mg, 307 $\mu$mol) in dry THF (3.0 mL), after that DEAD (53.5 mg, 307 $\mu$mol) was added dropwise to the solution at 0 °C. The resulting mixture was allowed to warm up to room temperature and stirred overnight. The reaction mixture was subjected to HPLC (0-1-6 min., 40-40-80% water – ACN, flow: 60 mL/min, column: XBridge C18 OBD 100×30 mm, 5 $\mu$m) to afford N,N-dimethyl-5-[3-[2-methyl-4-[5-(trifluoromethyl)-1,2,4-oxadiazol-3-yl]phenoxy]propyl]isoxazole-3-carboxamide (53.3 mg, 126 $\mu$mol, 61.3% yield) as a white solid. $^1$H NMR (500 MHz, dmso) $\delta_H$ 2.14 – 2.20 (m, 2H), 2.22 (s, 3H), 2.98 (s, 3H), 3.01 (t, 2H), 3.04 (s, 3H), 4.14 (t, 2H), 6.53 (s, 1H), 7.13 (d, 1H), 7.84 (d, 1H), 7.86 (dd, 1H). LCMS(ESI): $[M+H]^+$ m/z: calcd 425.16; found 425.0.

#### Synthesis of N,N-dimethyl-5-[2-[[2-methyl-4-[5-(trifluoromethyl) -1,2,4-oxadiazol-3-yl]phenyl]methoxy]ethyl]isoxazole-3-carboxamide (Compound 22)

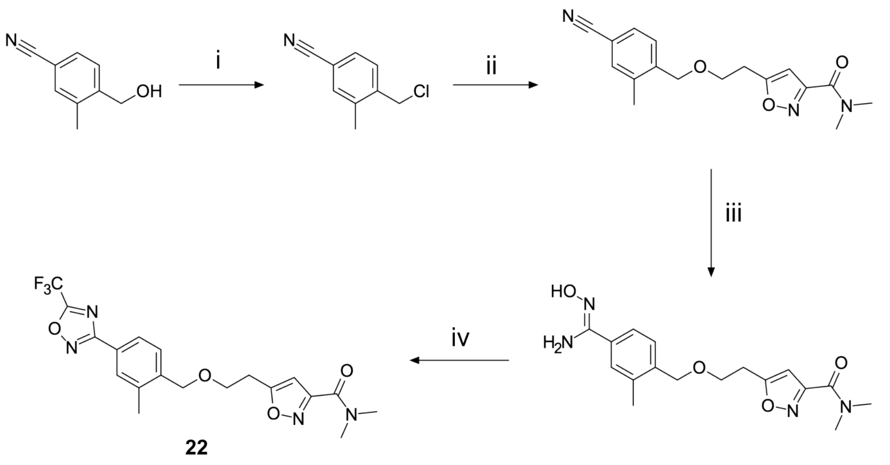


Scheme S2: Reagents and conditions: (i) $SOCl_2$, $CHCl_3$, 50 °C, 3h (ii) 5-(2-hydroxyethyl)-N,N-dimethyl-isoxazole-3-carboxamide, NaH, DMF, 0 °C to RT, 16h (iii) hydroxylamine hydrochloride, $Na_2CO_3$, $H_2O$/MeOH, 60 °C, 16h (iv) TFAA, pyridine, THF, 0 °C to 60 °C, 16h

#### Step i: Data for 4-(chloromethyl)-3-methyl-benzonitrile)

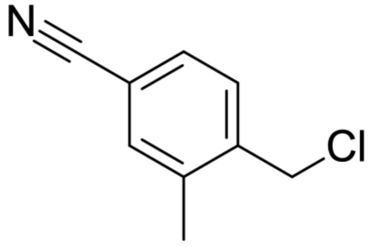

$SOCl_2$ (606 mg, 5.10 mmol, 372 $\mu$L) was added to a stirred solution of 4-(hydroxymethyl)-3-methyl-benzonitrile (500 mg, 3.40 mmol) in $CHCl_3$ (5.0 mL) at room temperature. The resulting mixture was stirred at 50 °C for 3 hr. The reaction mixture was cooled to room temperature and concentrated under reduced pressure to afford 4-(chloromethyl)-3-methyl-benzonitrile (560 mg, 3.38 mmol, 99.5% yield) as a yellow solid which was used in the next step without further purification. $^1$H NMR (400 MHz, CDCl3) $\delta_H$ 2.47 (s, 3H), 4.60 (s, 2H), 7.41 – 7.54 (m, 3H).

#### Step ii: Data for 5-[2-[(4-cyano-2-methyl-phenyl)methoxy]ethyl]-N,N-dimethyl-isoxazole-3-carboxamide

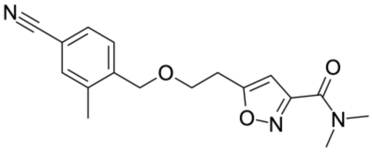

NaH (15.9 mg, 664 $\mu$mol, 60% dispersion in mineral oil) was added at 0 °C to a solution of 5-(2-hydroxyethyl)-N,N-dimethyl-isoxazole-3-carboxamide (111 mg, 604 $\mu$mol) in DMF (3.0 mL). The resulting mixture was stirred at 0 °C for 30 min. 4-(Chloromethyl)-3-methyl-benzonitrile (100 mg, 604 $\mu$mol) was added at 0 °C to the mixture. The resulting mixture was stirred at room temperature for 16 hr. The reaction mixture was concentrated under reduced pressure. The residue was subjected to reverse phase HPLC (0-1.3-6.3 min., 15-15-40% water – ACN, flow: 30 mL/min, column: Chromatorex 18 SMB100-5T 100×19 mm, 5 $\mu$m) to afford 5-[2-[(4-cyano-2-methyl-phenyl)methoxy]ethyl]-N,N-dimethyl-isoxazole-3-carboxamide (7.00 mg, 22.3 $\mu$mol, 3.70% yield) as a yellow oil. LCMS(ESI): $[M+H]^+$ m/z: calcd 314.17; found 314.2.

#### Step iii: Data for 5-[2-[[4-(N-hydroxycarbamimidoyl)-2-methyl-phenyl]methox N,N-dimethyl-isoxazole-3-carboxamide

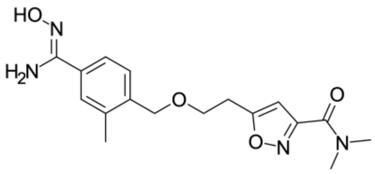

Hydroxylamine hydrochloride (4.66 mg, 67.0 $\mu$mol) and $Na_2CO_3$ (2.37 mg, 22.3 $\mu$mol) were added at room temperature to a stirred solution of 5-[2-[(4-cyano-2-methyl-phenyl)methoxy]ethyl]-N,N-dimethyl-isoxazole-3- carboxamide (7.00 mg, 22.3 $\mu$mol) in water (0.3 mL) and EtOH (0.5 mL). The resulting mixture was stirred at 60 °C for 16 hr. The reaction mixture was cooled to room temperature and concentrated under reduced pressure. The residue was redissolved in DCM (5 mL) and washed with water (3×4 mL). The organic layer was separated, dried over anhydrous sodium sulfate and concentrated under reduced pressure to afford 5-[2-[[4-(N-hydroxycarbamimidoyl)-2-methyl-phenyl]methoxy]ethyl]-N,N-dimethyl-isoxazole-3-carboxamide (7.00 mg, 20.2 $\mu$mol, 90.5% yield) as a white powder which was used in the next step without further purification. LCMS(ESI): $[M+H]^+$ m/z: calcd 347.17; found 347.2.

#### Step iv: Data for N,N-dimethyl-5-[2-[[2-methyl-4-[5-(trifluoromethyl)-1,2,4-oxadiazol-3-yl]phenyl]methoxy]ethyl]isoxazole-3-carboxamide (Compound 22)

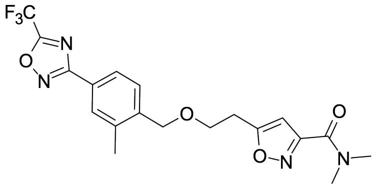

Pyridine (5.70 $\mu$L) and TFAA (10.6 mg, 50.5 $\mu$mol, 7.02 $\mu$L) were added to a stirred solution of 5-[2-[[4-(N-hydroxycarbamimidoyl)-2-methyl-phenyl]methoxy]ethyl]-N,N-dimethyl-isoxazole-3-carboxamide (7.00 mg, 20.2 $\mu$mol) in THF (1.0 mL) at 0 °C. The resulting mixture was stirred at 60 °C for 16 hr. The reaction mixture was cooled to room temperature and concentrated under reduced pressure. The residue was subjected to reverse phase HPLC (0-1.3-5.3 min., 45-45-65% water – ACN, flow: 30 mL/min, column: Chromatorex 18 SMB100-5T 100×19 mm, 5 $\mu$m) to afford N,N-dimethyl-5-[2-[[2-methyl-4-[5-(trifluoromethyl)-1,2,4-oxadiazol-3-yl]phenyl]methoxy]ethyl]isoxazole-3-carboxamide (3.50 mg, 8.25 $\mu$mol, 40.8% yield) as a yellow oil. $^1$H NMR (500 MHz, $CD_3OD$) $\delta_H$ 2.36 (s, 3H), 3.10 (s, 3H), 3.16 (t, 2H), 3.18 (s, 3H), 3.89 (t, 2H), 4.63 (s, 2H), 6.41 (s, 1H), 7.49 (d, 1H), 7.87 – 7.93 (m, 2H). LCMS(ESI): $[M+H]^+$ m/z: calcd 425.16; found 425.2.

#### Synthesis of N,N-dimethyl-5-[2-[2-methyl-4-[5-(trifluoromethyl)-1,2,4-oxadiazol-3-yl]phenyl]ethoxymethyl]isoxazole-3-carboxamide (Compound 23)

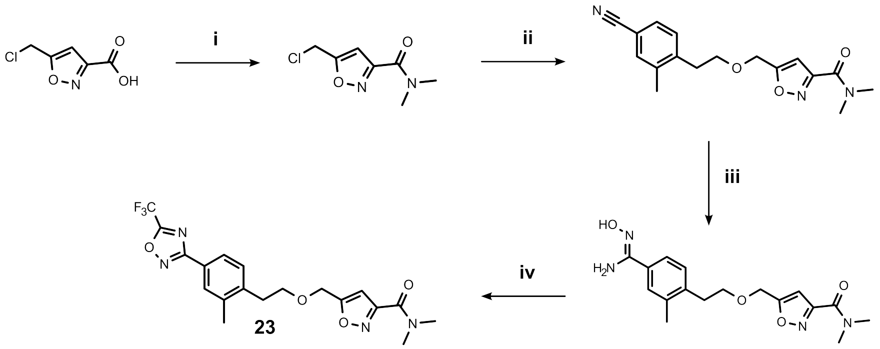

Scheme S3: Reagents and conditions: (i) $Me_2NH$, $(COCl)_2$, DIPEA, THF / DCM, 0 °C to RT, 16h (ii) 4-(2-hydroxyethyl)-3-methyl-benzonitrile, tetrabutylammonium hydrogen sulfate, NaOH, DCM, RT, 16h (iii) hydroxylamine hydrochloride, $NaHCO_3$, $^i$PrOH, RT to reflux, 16h (iv) TFAA, pyridine, 80 °C, 16h

#### Step i: Data for 5-(chloromethyl)-N,N-dimethyl-isoxazole-3-carboxamide

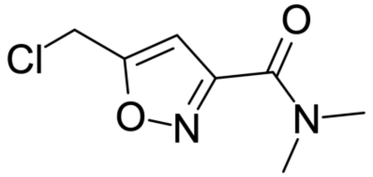

Oxalyl chloride (169 mg, 1.33 mmol, 116 $\mu$L) and a drop of DMF were added to a stirred solution of 5-(chloromethyl)isoxazole-3-carboxylic acid (143 mg, 885 $\mu$mol) in DCM (10 mL) at room temperature. The resulting mixture was stirred at room temperature for 1 hr. The reaction mixture was concentrated under reduced pressure. The residue was redissolved in THF (6.0 mL) and added dropwise to a solution of dimethylamine (79.4 mg, 974 $\mu$mol, 122 $\mu$L, HCl) and DIPEA (137 mg, 1.06 mmol, 185 $\mu$L) in THF (6.0 mL) at 0°C. The resulting mixture was allowed to warm up to room temperature and stirred overnight. The reaction mixture was concentrated under reduced pressure. The residue was partitioned between EtOAc (10 mL) and water (5.0 mL). The aqueous layer was extracted with EtOAc (5.0 mL). The combined organic layers were washed with brine (5.0 mL), dried over anhydrous sodium sulfate and concentrated under reduced pressure to afford 5-(chloromethyl)-N,N-dimethyl-isoxazole-3-carboxamide (150 mg, 740 $\mu$mol, 83.6% yield) as red oil which was used in the next step without further purification. LCMS(ESI): $[M+H]^+$ m/z: calcd 189.05; found 189.0.

#### Step ii: Data for 5-[2-(4-cyano-2-methyl-phenyl)ethoxymethyl]-N,N-dimethyl-isoxazole-3-carboxamide

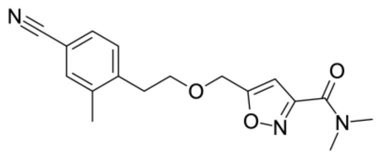

Tetrabutylammonium hydrogen sulfate (31.6 mg, 93.1 $\mu$mol) and 5-(chloromethyl)-N,N-dimethyl-isoxazole-3-carboxamide (105 mg, 558 $\mu$mol) were added to a vigorously stirred mixture of a solution of 4-(2-hydroxyethyl)-3-methyl-benzonitrile (7.50 mg, 465 $\mu$mol) in DCM (10 mL) and aqueous NaOH solution (5.0 mL, 15% wt.) at room temperature. The resulting mixture was stirred at room temperature overnight. The reaction mixture was washed with water (10 mL). The aqueous layer was separated and extracted with DCM (2×10 mL). The combined organic layers were washed with water (10 mL), brine (10 mL), dried over anhydrous sodium sulfate and concentrated under reduced pressure to afford 5-[2-(4-cyano-2-methyl-phenyl)ethoxymethyl]-N,N-dimethyl-isoxazole-3-carboxamide (150 mg, 192 $\mu$mol, 41.2% yield) as a brown oil which was used in the next step without further purification. LCMS(ESI): $[M+H]^+$ m/z: calcd 314.15; found 314.2.

#### Step iii: Data for 5-[2-[4-(N-hydroxycarbamimidoyl)-2-methyl-phenyl] ethoxymethyl]-N,N-dimethyl-isoxazole-3-carboxamide

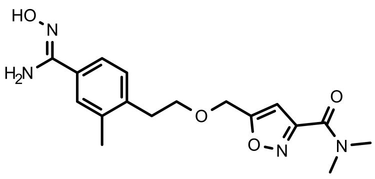

$NaHCO_3$ (96.5 mg, 1.15 mmol) and hydroxylamine hydrochloride (39.9 mg, 574 $\mu$mol) were added to a stirred solution of 5-[2-(4-cyano-2-methyl-phenyl)ethoxymethyl]-N,N-dimethyl-isoxazole-3-carboxamide (150 mg, 192 $\mu$mol) in iPrOH (10.1 mL) at room temperature. The resulting mixture was refluxed overnight. The reaction mixture was cooled to room temperature and filtered. The filter cake was washed with iPrOH (5 mL). The filtrate was concentrated under reduced pressure to afford 5-[2-[4-(N-hydroxycarbamimidoyl)-2-methyl-phenyl]ethoxymethyl]-N,N-dimethyl-isoxazole-3-carboxamide (130 mg, 184 $\mu$mol, 96.1% yield) as a brown solid which was used in the next step without further purification. LCMS(ESI): $[M+H]^+$ m/z: calcd 347.17; found 347.2.

#### Step iv: Synthesis of N,N-dimethyl-5-[2-[2-methyl-4-[5-(trifluoromethyl)-1,2,4-oxadiazol-3-yl]phenyl]ethoxymethyl]isoxazole-3-carboxamide (Compound 23)

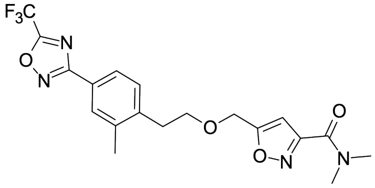

TFAA (116 mg, 552 $\mu$mol, 76.7 $\mu$L) was added to a stirred solution of 5-[2-[4-(N-hydroxycarbamimidoyl)-2-methyl-phenyl]ethoxymethyl]-N,N-dimethyl-isoxazole-3- carboxamide (130 mg, 184 $\mu$mol) in pyridine (4.9 mL). The resulting mixture was stirred at 80 °C overnight. The reaction mixture was cooled to room temperature and concentrated under reduced pressure. The residue was subjected to reverse phase HPLC (0-1-5 min., 45-45-75% water – ACN, flow: 30 mL/min, column: Chromatorex 18 SMB100-5T 100×19 mm, 5 $\mu$m) to afford N,N-dimethyl-5-[2-[2-methyl-4-[5-(trifluoromethyl)-1,2,4-oxadiazol-3-

yl]phenyl]ethoxymethyl]isoxazole-3-carboxamide (58.0 mg, 137 $\mu$mol, 74.3% yield) as a brown solid. $^1$H NMR (500 MHz, dmso) $\delta_H$ 2.37 (s, 3H), 2.94 (t, 2H), 2.98 (s, 3H), 3.03 (s, 3H), 3.74 (t, 2H), 4.67 (s, 2H), 6.62 (s, 1H), 7.41 (d, 1H), 7.80 (d, 1H), 7.84 (s, 1H). LCMS(ESI): $[M+H]^+$ m/z: calcd 425.16; found 425.0.

#### Synthesis of N,N-dimethyl-5-[2-[[2-methyl-4-[5-(trifluoromethyl) - 1,2,4-oxadiazol-3-yl]phenyl]methoxy]ethyl]isoxazole-3-carboxamide (Compound 24)

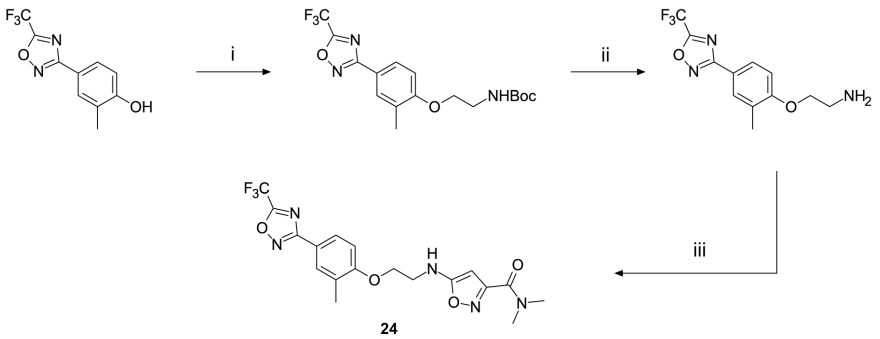


Scheme S4: Reagents and conditions: (i) tert-butyl 2,2-dioxooxathiazolidine-3-carboxylate, NaH, DMF, RT, 15h (ii) AcCl, MeOH, 0 °C to RT, 16h (iii) 5-chloro-N,N-dimethyl-isoxazole-3-carboxamide, $K_2CO_3$, DMF, 120 °C 48h

#### Step i: Data for tert-butyl N-[2-[2-methyl-4-[5-(trifluoromethyl)-1,2,4-oxadiazol-3-yl]phenoxy]ethyl]carbamate

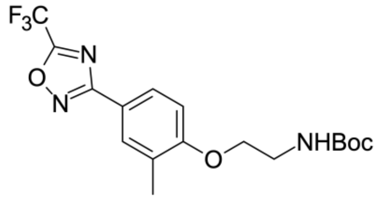


NaH (23.6 mg, 983 $\mu$mol, 60% dispersion in mineral oil) was added to a solution of 2-methyl-4-[5-(trifluoromethyl)-1,2,4-oxadiazol-3-yl]phenol (200 mg, 819 $\mu$mol) in DMF (5.0 mL) at room temperature. The resulting mixture was stirred at ambient temperature for 2 hr. Then, tert-butyl 2,2-dioxooxathiazolidine-3-carboxylate (219 mg, 983 $\mu$mol) was added in one portion to the mixture. The resulting mixture was stirred at room temperature for 15 hr. The reaction mixture was concentrated under reduced pressure. The residue was redissolved in EtOAc (30 mL) and washed with brine (3×30 mL). The combined organic layers were dried over anhydrous sodium sulfate and concentrated under reduced pressure to afford tert-butyl N-[2-[2-methyl-4-[5-(trifluoromethyl)-1,2,4-oxadiazol-3-yl]phenoxy]ethyl]carbamate (100 mg, 258 $\mu$mol, 31.5% yield) as a yellow solid which was used in the next step without further purification. $^1$H NMR (500 MHz, CDCl3) $\delta_H$ 1.46 (s, 9H), 2.29 (s, 3H), 3.54 - 3.65 (m, 2H), 4.10 (t, 2H), 4.94 (br. s, 1H), 6.90 (d, 1H), 7.88 – 7.95 (m, 2H). LCMS(ESI): [M+H-Boc]+ m/z: calcd 288.1; found 288.0.

#### Step ii: Data for 2-[2-methyl-4-[5-(trifluoromethyl)-1,2,4-oxadiazol-3-yl]phenoxy]ethanamine

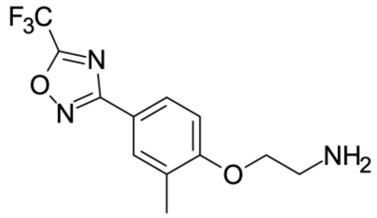


Acetyl chloride (30.4 mg, 387 $\mu$mol, 23.5 $\mu$L) was added dropwise to a stirred MeOH (5.0 mL) at 0 ºC. Tert-butyl N-[2-[2-methyl-4-[5-(trifluoromethyl)-1,2,4-oxadiazol-3-yl]phenoxy]ethyl]carbamate (100 mg, 258 $\mu$mol) was added to the obtained solution. The resulting mixture was stirred at room temperature for 15 hr. The reaction mixture was concentrated under reduced pressure to afford 2-[2-methyl-4-[5-(trifluoromethyl)-1,2,4-oxadiazol-3-yl]phenoxy]ethanamine (75.0 mg, 232 $\mu$mol, 89.8% yield, HCl) as a yellow solid which was used in the next step without further purification. LCMS(ESI): $[M+H]^+$ m/z: calcd 288.1; found 288.2.

#### Step iii: Data for N,N-dimethyl-5-[2-[2-methyl-4-[5-(trifluoromethyl)-1,2,4-oxadiazol-3-yl]phenoxy]ethylamino]isoxazole-3-carboxamide (Compound 24)

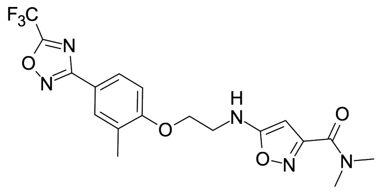

Potassium carbonate (96.1 mg, 695 $\mu$mol) and 5-chloro-N,N-dimethyl-isoxazole-3-carboxamide (40.5 mg, 232 $\mu$mol) were added to a solution of 2-[2-methyl-4-[5-(trifluoromethyl)-1,2,4-oxadiazol-3-yl]phenoxy]ethanamine (75.0 mg, 232 $\mu$mol, HCl) in DMF (4.0 mL). The resulting mixture was stirred at 120 °C for 48 hr. The reaction mixture was cooled to room temperature, filtered and subjected to HPLC (0-1-5 min., 30-30-60% water – ACN, flow: 30 mL/min, column: Chromatorex 18 SMB100-5T 100×19 mm, 5 $\mu$m) to afford N,N-dimethyl-5-[2-[2-methyl-4-[5-(trifluoromethyl)-1,2,4-oxadiazol-3-yl]phenoxy]ethylamino]isoxazole-3-carboxamide (1.50 mg, 3.53 $\mu$mol, 1.52% yield) as a yellow gum. $^1$H NMR (500 MHz, $CD_3OD$) $\delta_H$ 2.29 (s, 3H), 3.07 (s, 3H), 3.21 (s, 3H), 3.86 (t, 2H), 4.12 (t, 2H), 5.47 (s, 1H), 7.25 (d, 1H), 7.60 (dd, 1H), 7.69 (d, 1H). LCMS(ESI): $[M+H]^+$ m/z: calcd 426.15; found 426.2.

#### Synthesis of 5-[3-(5-cyanoindol-1-yl)propyl]-N,N-dimethyl-isoxazole-3-carboxamide (Compound 25) and N,N-dimethyl-5-[3-[5-[5-(trifluoromethyl)-1,2,4-oxadiazol-3-yl]indol-1-yl]propyl]isoxazole-3-carboxamide (Compound 26)

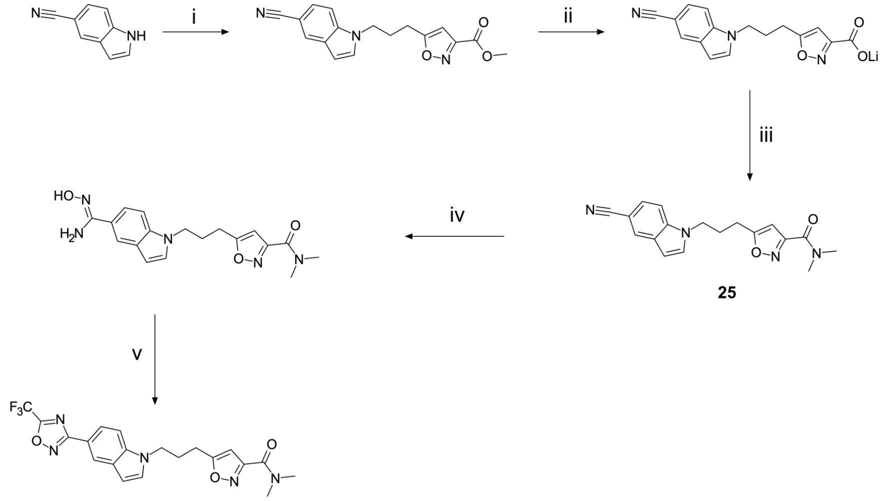


Scheme S5: Reagents and conditions: (i) methyl 5-(3-bromopropyl)isoxazole-3-carboxylate, NaH, DMF, 0 °C to 50 °C, 16h (ii) LiOH, $H_2O$, 0 °C to RT, 16h (iii) $Me_2NH$, HATU, DIPEA, DMF, RT, 16h (iv) hydroxylamine hydrochloride, $NaHCO_3$, EtOH, RT to 70 °C, 16h (v) TFAA, RT, 16h

#### Step i: Data for methyl 5-[3-(5-cyanopyrrolo[3,2-b]pyridin-1-yl)propyl] isoxazole-3-carboxylate

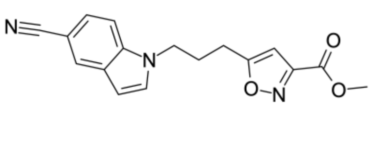

1H-indole-5-carbonitrile (249 mg, 1.39 mmol, HCl) was added to a stirred mixture of NaH (139 mg, 3.48 mmol, 60% dispersion in mineral oil) in DMF (20 mL) at 0 °C. After stirring for 30 min, methyl 5-(3-bromopropyl)isoxazole-3-carboxylate (449 mg, 1.81 mmol) was added

at 0 °C to the mixture. The resulting mixture was stirred at 50 °C for 16 hr. The reaction mixture was cooled to room temperature and diluted with water (30 mL). The resulting mixture was extracted with EtOAC (3×15 mL). The combined organic layers were washed with water (3×15 mL) and brine (15 mL), dried over anhydrous sodium sulfate and concentrated under reduced pressure. The residue was subjected to column chromatography ($SiO_2$, Hexane-MTBE 7:3) to afford methyl 5-[3-(5-cyanopyrrolo[3,2-b]pyridin-1-yl)propyl]isoxazole-3-carboxylate (600 mg, 1.55 mmol, 92.4% yield) as a yellow solid. $^1$H NMR (500 MHz, dmso) $\delta_H$ 2.18 (p, 2H), 2.81 (t, 2H), 3.87 (s, 3H), 4.33 (t, 2H), 6.61 (d, 1H), 6.68 (s, 1H), 7.49 (d, 1H), 7.62 (d, 1H), 7.69 (d, 1H), 8.08 (s, 1H). LCMS(ESI): $[M+H]^+$ m/z: calcd 310.12; found 310.2.

#### Step ii: Data for [5-[3-(5-cyanopyrrolo[3,2-b]pyridin-1-yl)propyl]isoxazole-3-carbonyl]oxylithium

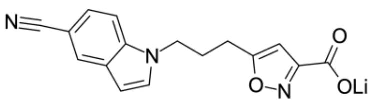


Water (200 $\mu$L) and LiOH·$H_2O$ (200 mg, 645 $\mu$mol) were added to a stirred solution of methyl 5-[3-(5-cyanopyrrolo[3,2-b]pyridin-1-yl)propyl]isoxazole-3-carboxylate (200 mg, 645 $\mu$mol) in THF (5.0 mL) at 0 °C. The resulting mixture was stirred at room temperature for 16 hr. The reaction mixture was concentrated under reduced pressure to afford [5-[3-(5-cyanopyrrolo[3,2-b]pyridin-1-yl)propyl]isoxazole-3-carbonyl]oxylithium (340 mg, 824 $\mu$mol, 72.8% yield) as a yellow gum which was used in the next step without further purification. LCMS(ESI): [M-Li]- m/z: calcd 294.3; found 294.0.

#### Step iii: Data for 5-[3-(5-cyanoindol-1-yl)propyl]-N,N-dimethyl-isoxazole-3-carboxamide (Compound 25)

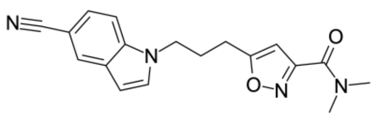

N-Methylmethanamine (54.0 mg, 662 $\mu$mol, 83.0 $\mu$L, HCl), HATU (302 mg, 794 $\mu$mol) and DIPEA (257 mg, 1.99 mmol, 346 $\mu$L) were added sequentially to a stirred solution of [5-[3-(5-cyanoindol-1-yl)propyl]isoxazole-3-carbonyl]oxylithium (199 mg, 662 $\mu$mol) in DMF (10 mL) at room temperature. The resulting mixture was stirred at room temperature for 16 hr. The reaction mixture was diluted with water (20 mL). The resulting mixture was extracted with EtOAc (2×15 mL). The combined organic layers were washed with an aqueous solution of $NH_4Cl$ (10 mL), an aqueous solution of $NaHCO_3$ (10 mL), water (2×10 mL) and brine (10 mL). The organic layer was dried over anhydrous sodium sulfate and concentrated under reduced pressure to afford 5-[3-(5-cyanoindol-1-yl)propyl]-N,N-dimethyl-isoxazole-3-carboxamide (180 mg, 452 $\mu$mol, 80.1% yield) as a yellow solid which was used in the next step without further purification. LCMS(ESI): $[M+H]^+$ m/z: calcd 323.15; found 323.2.

Note: purification of a similar sample under HPLC conditions (0-1-5 min., 40-40-65% water – MeOH, flow: 30 mL/min, column: Chromatorex 18 SMB100-5T 100×19 mm, 5 $\mu$m) provided a pure sample with the following NMR data: $^1$H NMR (500 MHz, $CD_3OD$) $\delta_H$ 2.28 (p, 2H), 2.81 (t, 2H), 3.09 (s, 3H), 3.16 (s, 3H), 4.34 (t, 2H), 6.31 (s, 1H), 6.61 (d, 1H), 7.41 – 7.46 (m, 2H), 7.57 (d, 1H), 7.98 (s, 1H).

#### Step iv: Data for 5-[3-[5-[(Z)-N'-hydroxycarbamimidoyl]indol-1-yl]propyl]-N,N-dimethyl-isoxazole-3-carboxamide

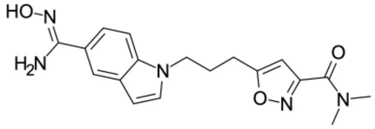

Hydroxylamine hydrochloride (65.8 mg, 946 $\mu$mol) and $NaHCO_3$ (88.3 mg, 1.05 mmol) were added respectively at room temperature to a stirred solution of 5-[3-(5-cyanoindol-1-yl)propyl]-N,N-dimethyl-isoxazole-3-carboxamide (167 mg, 526 $\mu$mol) in EtOH (15 mL). The resulting mixture was stirred at 70 °C for 16 hr. The reaction mixture was cooled to room temperature and filtered. The filtrate was concentrated under reduced pressure to afford 5-[3-[5-[(Z)-N'-hydroxycarbamimidoyl]indol-1-yl]propyl]-N,N-dimethyl-isoxazole-3-carboxamide (200 mg, 310 $\mu$mol, 55.4% yield) as a yellow gum which was used in the next step without further purification. LCMS(ESI): $[M+H]^+$ m/z: calcd 356.17; found 356.2.

#### Step iii: Data for N,N-dimethyl-5-[3-[5-[5-(trifluoromethyl)-1,2,4-oxadiazol-3-yl]indol-1-yl]propyl]isoxazole-3-carboxamide (Compound 26)

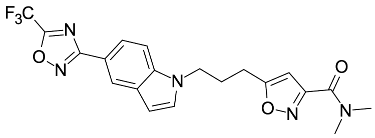


TFAA (150 mg, 716 $\mu$mol, 99.5 $\mu$L) was added to a stirred solution of 5-[3-[5-[(Z)-N'-hydroxycarbamimidoyl]indol-1-yl]propyl]-N,N-dimethyl-isoxazole-3-carboxamide (170 mg, 477 $\mu$mol) in THF (10 mL) at room temperature. The resulting mixture was stirred at room temperature for 16 hr. The reaction mixture was concentrated under reduced pressure. The residue was quenched with an aqueous solution of $NaHCO_3$ (10 mL). The aqueous layer was separated and extracted with MTBE (2×10 mL). The combined organic layers were washed with water (10 mL), brine (10 mL), dried over anhydrous sodium sulfate and concentrated under reduced pressure. The residue was subjected to reverse phase HPLC (0-1-5 min., 55-55-75% water – ACN, flow: 30 mL/min, column: Chromatorex 18 SMB100-5T 100×19 mm, 5 $\mu$m) to afford N,N-dimethyl-5-[3-[5-[5-(trifluoromethyl)-1,2,4-oxadiazol-3-yl]indol-1-yl]propyl]isoxazole-3-carboxamide (68.0 mg, 157 $\mu$mol, 51.6% yield) as an off-white gum. $^1$H NMR (500 MHz, dmso) $\delta_H$ 2.19 (p, 2H), 2.79 (t, 2H), 2.96 (s, 3H), 3.01 (s, 3H), 4.33 (t, 2H), 6.47 (s, 1H), 6.65 (d, 1H), 7.56 (d, 1H), 7.71 (d, 1H), 7.81 (d, 1H), 8.31 (s, 1H). LCMS(ESI): $[M+H]^+$ m/z: calcd 434.16; found 434.2.

**Synthesis of N,N-dimethyl-5-[3-[5-[5-(trifluoromethyl)-1,2,4-oxadiazol-3-yl]indazol-1-yl]propyl]isoxazole-3-carboxamide (Compound 27)**

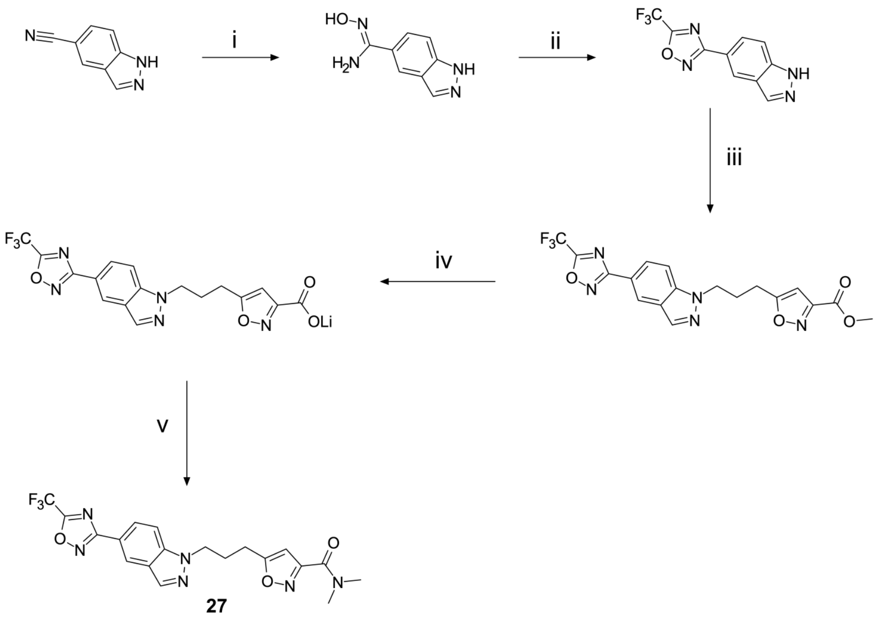

Scheme S6: Reagents and conditions: (i) hydroxylamine hydrochloride, $NaHCO_3$, MeOH, RT, 16h (ii) TFAA, THF, RT, 16h (iii) methyl 5-(3-bromopropyl)isoxazole-3-carboxylate, $K_2CO_3$, DMF, RT to 50 °C, 16h (iv) LiOH, $H_2O$, MeOH, RT, 16h (v) $Me_2NH$, HATU, DIPEA, DMF, RT to 50 °C, 16h

#### Step i: Data for N'-hydroxy-1H-indazole-5-carboxamidine

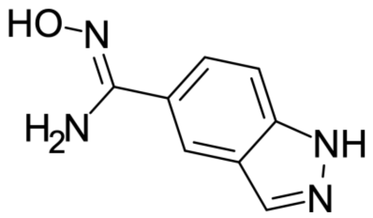

$NaHCO_3$ (1.47 g, 17.5 mmol) and hydroxylamine hydrochloride (485 mg, 6.99 mmol) were added to a stirred solution of 1H-indazole-5-carbonitrile (500 mg, 3.49 mmol) in MeOH (18 mL) at room temperature. The resulting mixture was stirred at room temperature overnight. The reaction mixture was filtered. The filtercake was washed with MeOH (2×5 mL). The combined filtrate was concentrated under reduced pressure to afford N'-hydroxy-1H-indazole-5-carboxamidine (600 mg, 2.72 mmol, 78.0% yield) as a brown solid which was used in the next step without further purification. LCMS(ESI): $[M+H]^+$ m/z: calcd 177.07; found 177.2.

#### Step ii: Data for 3-(1H-indazol-5-yl)-5-(trifluoromethyl)-1,2,4-oxadiazole

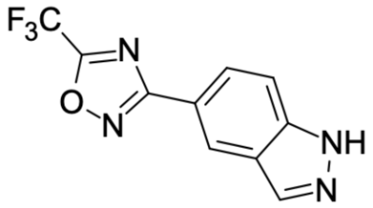

TFAA (1.45 g, 6.90 mmol, 960 $\mu$L) was added to a stirred solution of N'-hydroxy-1H-indazole-5-carboxamidine (600 mg, 2.72 mmol) in THF (15.0 mL) at room temperature. The resulting mixture was stirred at room temperature overnight. The reaction mixture was concentrated under reduced pressure. The residue was partitioned between EtOAc (20 mL) and water (10 mL). The aqueous layer was separated and extracted with EtOAc (10 mL). The combined organic layers were washed with water (10 mL) and brine (10 mL), dried over anhydrous sodium sulfate and concentrated under reduced pressure to afford 3-(1H-indazol-5-yl)-5-(trifluoromethyl)-1,2,4-oxadiazole (700 mg, 1.79 mmol, 65.7% yield) as a brown solid which was used in the next step without further purification. $^1$H NMR (500 MHz, dmso) $\delta_H$ 7.76 (d, 1H), 8.00 (d, 1H), 8.28 (s, 1H), 8.57 (s, 1H), 13.37 (br. s, 1H). LCMS(ESI): $[M+H]^+$ m/z: calcd 255.05; found 255.0.

#### Step iii: Data for methyl 5-[3-[5-[5-(trifluoromethyl)-1,2,4-oxadiazol-3-yl]indazol-1-yl]propyl]isoxazole-3-carboxylate

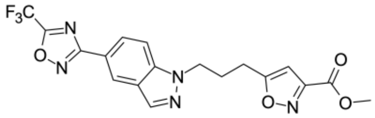

Methyl 5-(3-bromopropyl)isoxazole-3-carboxylate (533 mg, 2.15 mmol) and $K_2CO_3$ (619 mg, 4.48 mmol) were added to a stirred solution of 3-(1H-indazol-5-yl)-5-(trifluoromethyl)-1,2,4-oxadiazole (700 mg, 1.79 mmol) in DMF (7.0 mL) at room temperature. The resulting mixture was stirred at 50 °C overnight. The reaction mixture was cooled to room temperature and filtered. The filtrate was subjected to preparative HPLC (0-1-6 min., 50-50-70% water – ACN, flow: 30 mL/min, column: Chromatorex 18 SMB100-5T 100×19 mm, 5 $\mu$m) to afford methyl 5-[3-[5-[5-(trifluoromethyl)-1,2,4-oxadiazol-3-yl]indazol-1-yl]propyl]isoxazole-3-carboxylate (89.0 mg, 211 $\mu$mol, 11.8% yield) as a yellow solid. $^1$H NMR (500 MHz, $CD_3OD$) $\delta_H$ 2.41 (p, 2H), 2.92 (t, 2H), 3.95 (s, 3H), 4.62 (t, 2H), 6.58 (s, 1H), 7.85 (d, 1H), 8.14 (d, 1H), 8.29 (s, 1H), 8.62 (s, 1H). LCMS(ESI): $[M+H]^+$ m/z: calcd 422.11; found 422.2.

#### Step iv: Data for [5-[3-[5-[5-(trifluoromethyl)-1,2,4-oxadiazol-3-yl]indazol-1-yl]propyl]isoxazole-3-carbonyl]oxylithium

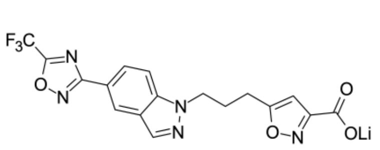


$LiOH \cdot H_2O$ (10.8 mg, 256 $\mu$mol) in water (1.0 mL) was added to a stirred solution of methyl 5-[3-[5-[5-(trifluoromethyl)-1,2,4-oxadiazol-3-yl]indazol-1-yl]propyl]isoxazole-3-carboxylate (90.0 mg, 214 $\mu$mol) in THF (1.0 mL) and MeOH (1.0 mL) at room temperature. The resulting mixture was stirred at room temperature overnight. The reaction mixture was concentrated under reduced pressure to afford [5-[3-[5-[5-(trifluoromethyl)-1,2,4-oxadiazol-3-yl]indazol-1-yl]propyl]isoxazole-3-carbonyl]oxylithium (100 mg, 26.6 $\mu$mol, 12.5% yield) as a brown solid which was used in the next step without further purification. LCMS(ESI): [M-Li+2H]+ m/z: calcd 408.09; found 408.0.

#### Step v: Data for N,N-dimethyl-5-[3-[5-[5-(trifluoromethyl)-1,2,4-oxadiazol-3-yl]indazol-1-yl]propyl]isoxazole-3-carboxamide (Compound 27)

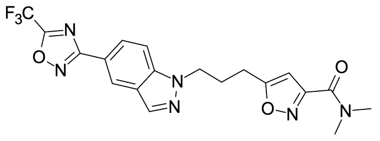


DIPEA (103 mg, 799 $\mu$mol, 139 $\mu$L) and HATU (202 mg, 532 $\mu$mol) were added to a stirred solution of [5-[3-[5-[5-(trifluoromethyl)-1,2,4-oxadiazol-3-yl]indazol-1-yl]propyl]isoxazole-3-carbonyl]oxylithium (100 mg, 26.6 $\mu$mol) and dimethylamine (32.6 mg, 399 $\mu$mol, 50.1 $\mu$L, HCl) in DMF (3.0 mL) at room temperature. The resulting mixture was stirred at 50 °C overnight. The reaction mixture was cooled to room temperature and subjected to preparative HPLC (0-5 min., 60-85% water – MeOH, flow: 30 mL/min, column: Chromatorex 18 SMB100-5T 100×19 mm, 5 $\mu$m) to afford N,N-dimethyl-5-[3-[5-[5-(trifluoromethyl)-1,2,4-oxadiazol-3-yl]indazol-1-yl]propyl]isoxazole-3-carboxamide (7.50 mg, 17.3 $\mu$mol, 64.9% yield) as a beige solid. $^1$H NMR (500 MHz, dmso) $\delta_H$ 2.26 (p, 2H), 2.81 (t, 2H), 2.96 (s, 3H), 3.01 (s, 3H), 4.55 (t, 2H), 6.45 (s, 1H), 7.90 (d, 1H), 8.03 (d, 1H), 8.29 (s, 1H), 8.55 (s, 1H). LCMS(ESI): $[M+H]^+$ m/z: calcd 435.15; found 435.0.

#### Data for N,N-dimethyl-5-[3-[5-[5-(trifluoromethyl)-1,2,4-oxadiazol-3-yl]pyrrolo[3,2-b]pyridin-1-yl]propyl]isoxazole-3-carboxamide (Compound 28)

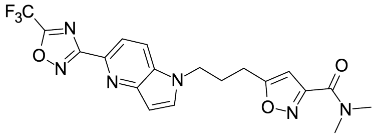


Compound 28 was made by a similar route to compound 26, starting with 1H-Pyrrolo[3,2-b]pyridine-5-carbonitrile, affording N,N-dimethyl-5-[3-[5-[5-(trifluoromethyl)-1,2,4-oxa diazol-3-yl]pyrrolo[3,2-b]pyridin-1-yl]propyl]isoxazole-3-carbox amide (56.0 mg, 129 $\mu$mol, 27.0% yield) as an off-white gum. $^1$H NMR (500 MHz, dmso) $\delta_H$ 2.21 (p, 2H), 2.79 (t, 2H), 2.96 (s, 3H), 3.01 (s, 3H), 4.36 (t, 2H), 6.46 (s, 1H), 6.75 (d, 1H), 7.90 – 7.96 (m, 2H), 8.18 (d, 1H). LCMS(ESI): $[M+H]^+$ m/z: calcd 435.15; found 435.2.

#### Data for N,N-dimethyl-5-[3-[5-[5-(trifluoromethyl)-1,2,4-oxadiazol-3-yl]pyrrolo[2,3-b]pyridin-1-yl]propyl]isoxazole-3-carboxamide (Compound 29)

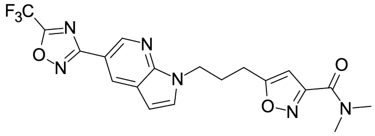


Compound 29 was made by a similar route to compound 26, starting with 1H-pyrrolo[2,3-b]pyridine-5-carbonitrile, affording N,N-dimethyl-5-[3-[5-[5-(trifluoromethyl)-1,2,4-oxadiazol-3-yl]pyrrolo[2,3-b]pyridin-1-yl]propyl]isoxazole-3-carboxamide

(150 mg, 345 $\mu$mol, 35.2% yield) as an off-white solid. $^{1}$H NMR (500 MHz, dmso) $\delta_{H}$ 2.24 (p, 2H), 2.80 (t, 2H), 2.97 (s, 3H), 3.02 (s, 3H), 4.40 (t, 2H), 6.47 (s, 1H), 6.68 (d, 1H), 7.78 (d, 1H), 8.64 (d, 1H), 8.90 (d, 1H). LCMS(ESI): $[M+H]^{+}$ m/z: calcd 435.15; found 435.2.

#### Data for N,N-dimethyl-5-[3-[5-[5-(trifluoromethyl)-1,2,4-oxadiazol-3-yl]pyrrolo[2,3-c]pyridin-1-yl]propyl]isoxazole-3-carboxamide (Compound 30)

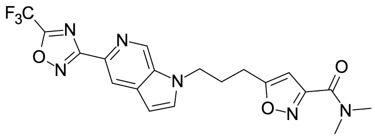


Compound 30 was made by a similar route to compound 26, starting with 1H-pyrrolo[2,3-c]pyridine-5-carbonitrile, affording N,N-dimethyl-5-[3-[5-[5-(trifluoromethyl)-1,2,4-oxadiazol-3-yl]pyrrolo[2,3-c]pyridin-1- yl]propyl]isoxazole-3-carboxamide (174 mg, 401 $\mu$mol, 35.7% yield) as a white solid. $^{1}$H NMR (500 MHz, dmso) $\delta_{H}$ 2.25 (p, 2H), 2.81 (t, 2H), 2.96 (s, 3H), 3.01 (s, 3H), 4.47 (t, 2H), 6.48 (s, 1H), 6.73 (d, 1H), 7.80 (d, 1H), 8.39 (s, 1H), 9.08 (s, 1H). LCMS(ESI): $[M+H]^{+}$ m/z: calcd 435.15; found 435.2.

#### Synthesis of N,N-dimethyl-5-[3-[4-[5-(trifluoromethyl)-1,2,4-oxadiazol-3-yl]phenoxy]phenyl]isoxazole-3-carboxamide (Compound 31)

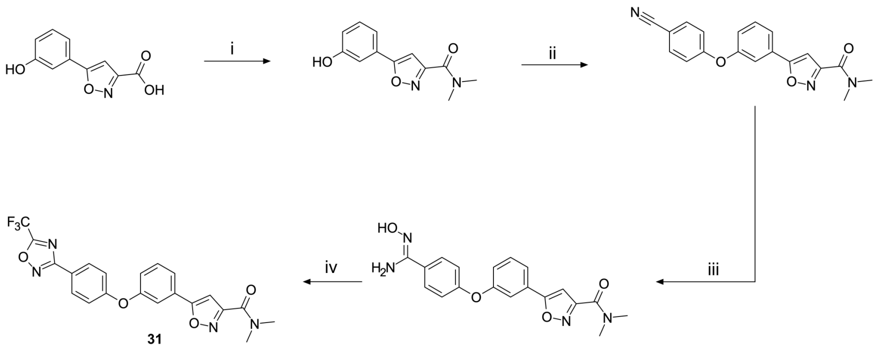

31

Scheme S7: Reagents and conditions: (i) $Me_2NH$, HATU, DIPEA, DMF, 50 °C, 16h (ii) 4-fluorobenzonitrile, $K_2CO_3$, 100 °C, 16h (iii) hydroxylamine hydrochloride, $NaHCO_3$, $^{i}PrOH$, 75 °C, 16h (iv) TFAA, pyridine, 50 °C, 16h

#### Step i: Data for 5-(3-hydroxyphenyl)-N,N-dimethyl-isoxazole-3-carboxamide

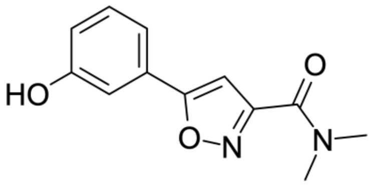

5-(3-Hydroxyphenyl)isoxazole-3-carboxylic acid (1.00 g, 4.87 mmol), HATU (2.22 g, 5.85 mmol) and N-methylmethanamine (1.99 g, 24.4 mmol, 3.06 mL) and DIPEA (6.30 g, 48.7 mmol) were mixed in dry DMF (30 mL). The resulting mixture was stirred at 50 °C overnight. The reaction mixture was cooled to room temperature, diluted with water (200 mL) and extracted with EtOAc (3×50 mL). The combined organic layers were washed with water (3×100 mL) and brine (100 mL), dried over anhydrous sodium sulfate and filtered. The filtrate was concentrated under reduced pressure to afford 5-(3-hydroxyphenyl)-N,N-dimethyl-isoxazole-3-carboxamide (620 mg, crude) as a yellow solid which was used in the next step without further purification. LCMS(ESI): $[M+H]^+$ m/z: calcd 233.09; found 233.0.

#### Step ii: Data for 5-[3-(4-cyanophenoxy)phenyl]-N,N-dimethyl-isoxazole-3-carboxamide

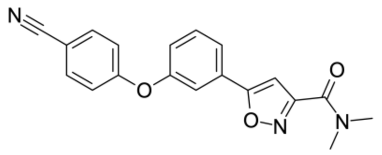

5-(3-Hydroxyphenyl)-N,N-dimethyl-isoxazole-3-carboxamide (620 mg, 2.67 mmol), $K_2CO_3$ (738 mg, 5.34 mmol) and 4-fluorobenzonitrile (388 mg, 3.20 mmol) were mixed in dry DMF (5 mL). The resulting mixture was stirred at 100 °C overnight. The mixture was cooled to room temperature and diluted with water (10 mL). The formed precipitate was collected by filtration, washed with water (5 mL) and air dried to afford 5-[3-(4-cyanophenoxy)phenyl]-N,N-dimethyl-isoxazole-3-carboxamide (288 mg, 734 $\mu$mol, 27.5% yield) as a brown solid which was used in the next step without further purification. $^1$H NMR (500 MHz, dmso) $\delta_H$ 3.03 (s, 3H), 3.11 (s, 3H), 7.19 (d, 2H), 7.32 (d, 1H), 7.36 (s, 1H), 7.65 (t, 1H), 7.73 (s, 1H), 7.83 (d, 1H), 7.88 (d, 2H). LCMS(ESI): $[M+H]^+$ m/z: calcd 334.12; found 334.0.

#### Step iii: Data for 5-[3-[4-[(Z)-N'-hydroxycarbamimidoyl]phenoxy]phenyl]-N,N-dimethyl-isoxazole-3-carboxamide

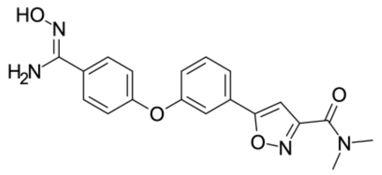

Hydroxylamine hydrochloride (62.5 mg, 900 $\mu$mol) and $NaHCO_3$ (151 mg, 1.80 mmol) were added to a solution of 5-[3-(4-cyanophenoxy)phenyl]-N,N-dimethyl-isoxazole-3-carboxamide (100 mg, 300 $\mu$mol) in i-PrOH (3.0 mL). The resulting mixture was stirred at 75 °C overnight. The reaction mixture was cooled to room temperature and filtered. The filtrate was concentrated under reduced pressure to afford 5-[3-[4-[(Z)-N'-hydroxycarbamimidoyl]phenoxy]phenyl]-N,N-dimethyl-isoxazole-3-carboxamide (80.0 mg, crude) as a yellow solid which was used in the next step without further purification. LCMS(ESI): $[M+H]^+$ m/z: calcd 367.14; found 367.0.

#### Step iv: Data for N,N-dimethyl-5-[3-[4-[5-(trifluoromethyl)-1,2,4-oxadiazol-3-yl]phenoxy]phenyl]isoxazole-3-carboxamide (Compound 31)

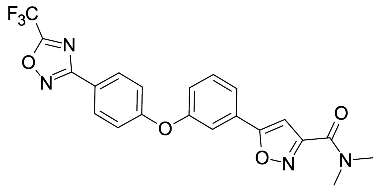

Trifluoroacetic anhydride (138 mg, 655 $\mu$mol, 91.0 $\mu$L) was added to a solution of 5-[3-[4-[(Z)-N'-hydroxycarbamimidoyl] phenoxy]phenyl]-N,N-dimethyl-isoxazole-3-carboxamide (80.0 mg, 218 $\mu$mol) in pyridine (1.0 mL). The resulting mixture was stirred at 50 °C overnight. The reaction mixture was cooled to room temperature and concentrated under reduced pressure. The residue was re-disolved with ACN (2.0 mL) and subjected to HPLC (0-1-6 min., 40-40-90% water – ACN, flow: 60 mL/min, column: XBridge OBD 100×30 mm, 5 $\mu$m) to afford N,N-dimethyl-5-[3-[4-[5-(trifluoromethyl)-1,2,4-oxadiazol-3-yl]phenoxy]phenyl]isoxazole-3-carboxamide (35.3 mg, 78.5 $\mu$mol, 35.9% yield) as a yellow oil. $^1$H NMR (500 MHz, dmso) $\delta_H$ 3.03 (s, 3H), 3.11 (s, 3H), 7.26 (d, 2H), 7.33 (d, 1H), 7.36 (s, 1H), 7.66 (t, 1H), 7.73 (s, 1H), 7.82 (d, 1H), 8.10 (d, 2H). LCMS(ESI): $[M+H]^+$ m/z: calcd 445.12; found 445.2.

#### Synthesis of N,N-dimethyl-5-[5-[2-methyl-4-[5-(trifluoromethyl)-1,2,4-oxadiazol-3-yl]phenoxy]-3-pyridyl]isoxazole-3-carboxamide (Compound 32)

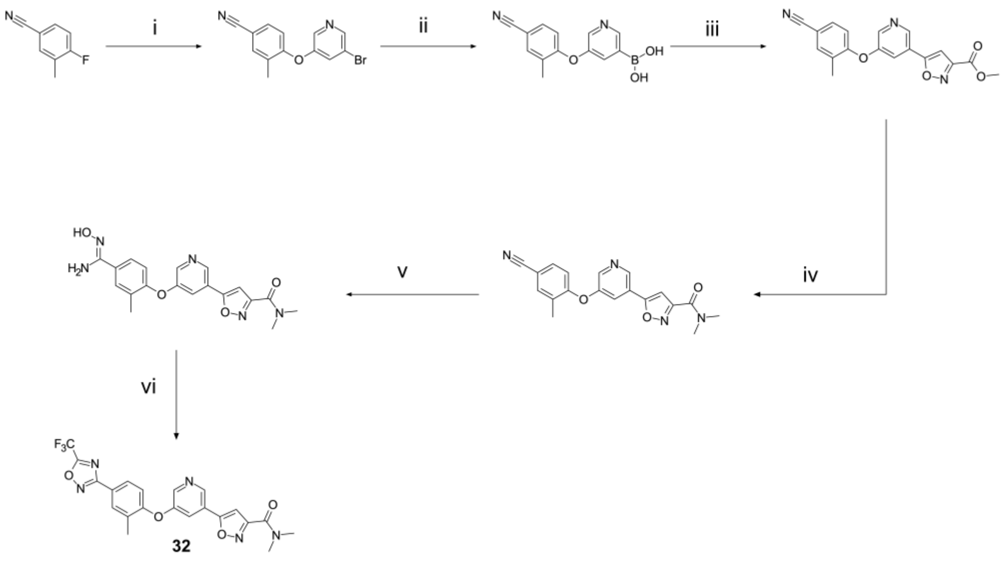

Scheme S8: Reagents and conditions: (i) 5-bromopyridin-3-ol, $K_2CO_3$, DMF, 115 °C, 15h (ii) bis(pinacolato)diboron, Pd(dppf)$Cl_2$, KOAc, dioxane, 90 °C, 15h (iii) methyl 5-bromoisoxazole-3-carboxylate, $Na_2CO_3$, Pd(dppf)$Cl_2$, $H_2O$, dioxane, 90 °C, 15h (iv) dimethylamine, $H_2O$, THF, 60 °C, 16h (v) hydroxylamine hydrochloride, EtOH, 70 °C, 15h (vi) TFAA, THF, RT, 15h

#### Step i: Data for 4-[(5-bromo-3-pyridyl)oxy]-3-methyl-benzonitrile

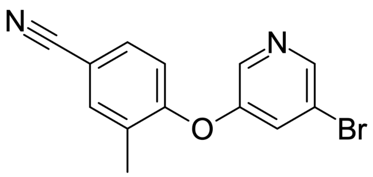

Potassium carbonate (1.59 g, 11.5 mmol) and 4-fluoro-3-methyl-benzonitrile (777 mg, 5.75 mmol) were added in one portion to a solution of 5-bromopyridin-3-ol (1.00 g, 5.75 mmol) in DMF (20 mL). The resulting mixture was stirred at 115 °C for 15 hr. The reaction mixture was cooled to room temperature, filtered and concentrated under reduced pressure. The residue was diluted with EtOAc (40 mL) the resulting mixture was washed with brine (3×20 mL). The obtained organic layer was dried over anhydrous sodium sulfate and concentrated under reduced pressure to afford 4-[(5-bromo-3-pyridyl)oxy]-3-methyl-benzonitrile (1.50 g, crude) as a yellow oil which was used in the next step without further purification. LCMS(ESI): $[M+H]^+$ m/z: calcd 289.0; found 289.0.

#### Step ii: Data for 5-(4-cyano-2-methylphenoxy)pyridin-3-ylboronic acid

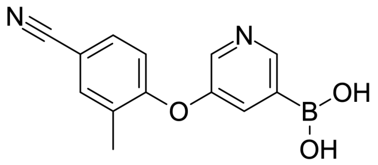


Potassium acetate (1.02 g, 10.4 mmol) was added to a solution of 4-[(5-bromo-3-pyridyl)oxy]-3-methyl-benzonitrile (1.50 g, 5.19 mmol) in dioxane (40 mL). The resulting mixture was stirred at ambient temperature for 5 min. 4,4,5,5-tetramethyl-2-(4,4,5,5-tetramethyl-1,3,2-dioxaborolan-2-yl)-1,3,2-dioxaborolane (1.32 g, 5.19 mmol) and bis(diphenylphosphino) ferrocene]dichloropalladium(II)-DCM (212 mg, 259 $\mu$mol) were added to the mixture under argon atmosphere. The resulting mixture was stirred with a reflux condenser under argon atmosphere at 90 °C for 15 hrs.. The resulting mixture was cooled to room temperature and concentrated under reduced pressure. Water (30 mL) was added to the residue and then resulting mixture was extracted with DCM (3×20 mL). The combined organic layers were dried over anhydrous sodium sulfate and filtered. The filtrate was concentrated under reduced pressure to afford 5-(4-cyano-2-methylphenoxy)pyridin-3-ylboronic acid (2.00 g, crude) as a brown solid which was used in the next step without further purification. LCMS(ESI): $[M+H]^+$ m/z: calcd 255.10; found 255.0.

#### Step iii: Data for methyl 5-[5-(4-cyano-2-methyl-phenoxy)-3 pyridyl] isoxazole-3-carboxylate

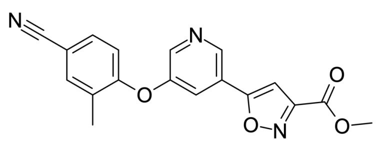

Sodium carbonate (631 mg, 5.95 mmol) was added to a solution of 5-(4-cyano-2-methylphenoxy)pyridin-3-ylboronic acid (1.00 g, crude) in dioxane (19.8 mL) and water (3.96 mL). The resulting mixture was stirred at ambient temperature for 5 min. Methyl 5-bromoisoxazole-3-carboxylate (613 mg, 2.97 mmol) and bis(diphenylphosphino) ferrocene]dichloropalladium(II)-DCM (122 mg, 149 $\mu$mol) were added to the mixture under argon atmosphere. The resulting mixture was stirred with a reflux condenser under argon atmosphere at 90 °C for 15 hrs. The reaction mixture was cooled to room temperature and concentrated under reduced pressure. The residue was subjected to HPLC (0-1-5 min., 35-35-60% water – ACN, flow: 30 mL/min, column: Chromatorex 18 SMB100-5T 100×19 mm, 5 $\mu$m) to afford methyl 5-[5-(4-cyano-2-methyl-phenoxy)-3-pyridyl]isoxazole-3-carboxylate (53.2 mg, 159 $\mu$mol, 5.33% yield) as a yellow solid which was used in the next step without further purification. LCMS(ESI): $[M+H]^+$ m/z: calcd 336.1; found 336.2.

#### Step iv: Data for 5-[5-(4-cyano-2-methyl-phenoxy)-3-pyridyl]-N,N-dimethyl-isoxazole-3-carboxamide

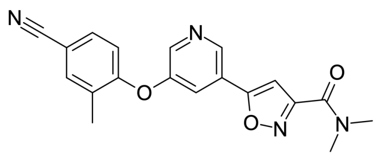

Dimethylamine solution (40% in water, 3.33 mL) was added in one portion to a solution of methyl 5-[5-(4-cyano-2-methyl-phenoxy)-3-pyridyl]isoxazole-3-carboxylate (30.0 mg, 89.5 $\mu$mol) in THF (700 $\mu$L). The resulting mixture was stirred at 65 °C for 16 hr. The mixture was cooled to room temperature and concentrated under reduced pressure to afford 5-[5-(4-cyano-2-methyl-phenoxy)-3-pyridyl]-N,N-dimethyl-isoxazole-3-carboxamide (30.0 mg, 86.1 $\mu$mol, 96.3% yield) as a yellow solid which was used in the next step without further purification. LCMS(ESI): $[M+H]^+$ m/z: calcd 349.13; found 349.0.

#### Step v: Data for 5-[5-[4-[(Z)-N'-hydroxycarbamimidoyl]-2-methyl-phenoxy]-3-pyridyl]-N,N-dimethyl-isoxazole-3-carboxamide

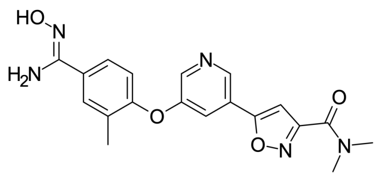


Hydroxylamine hydrochloride (18.0 mg, 258 $\mu$mol) was added in one portion to a solution of 5-[5-(4-cyano-2-methyl-phenoxy)-3-pyridyl]-N,N-dimethyl-isoxazole-3-carboxamide (50.0 mg, 144 $\mu$mol) in EtOH (10 mL). The resulting mixture was stirred at 70 °C for 15 hr. The reaction mixture was cooled to room temperature and concentrated under reduced pressure to afford 5-[5-[4-[(Z)-N'-hydroxycarbamimidoyl]-2-methyl-phenoxy]-3-pyridyl]-N,N-dimethyl-isoxazole-3-carboxamide (100 mg, crude) as a brown oil which was used in the next step without further purification. LCMS(ESI): $[M+H]^+$ m/z: calcd 382.15; found 382.2.

#### Step vi: Data for N,N-dimethyl-5-[5-[2-methyl-4-[5-(trifluoromethyl)-1,2,4-oxadiazol-3-yl]phenoxy]-3-pyridyl]isoxazole-3-carboxamide (Compound 32)

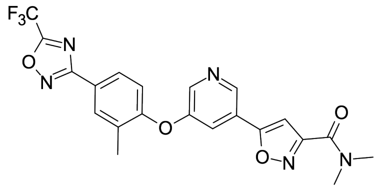


Trifluoroacetic anhydride (82.6 mg, 393 $\mu$mol, 54.5 $\mu$L) was added a stirred solution of 5-[5-[4-[(Z)-N'-hydroxycarbamimidoyl]-2-methyl-phenoxy]-3-pyridyl] -N,N-dimethyl-isoxazole-3-carboxamide (100 mg, 262 $\mu$mol) in THF (10 mL) at room temperature to. The resulting mixture was stirred at room temperature for 15 hr. The reaction mixture was concentrated under reduced pressure and quenched with an aqueous solution of $NaHCO_3$ (10 mL). The aqueous layer was separated and extracted with DCM (2×30 mL). The combined organic layers were washed with water (10 mL), brine (10 mL), dried over anhydrous sodium sulfate and concentrated under reduced pressure. The residue was subjected to HPLC (0-1-5 min., 45-45-70% water – ACN, flow: 30 mL/min, column: Chromatorex 18 SMB100-5T 100×19 mm, 5 $\mu$m)

to afford N,N-dimethyl-5-[5-[2-methyl-4-[5-(trifluoromethyl)-1,2,4-oxadiazol-3-yl]phenoxy]-3-pyridyl]isoxazole-3-carboxamide (12.8 mg, 27.9 $\mu$mol, 10.6% yield) as a beige solid. $^1$H NMR (500 MHz, $CD_3OD$) $\delta_H$ 2.40 (s, 3H), 3.13 (s, 3H), 3.24 (s, 3H), 7.16 (d, 1H), 7.20 (s, 1H), 7.91 (t, 1H), 8.02 (dd, 1H), 8.14 (s, 1H), 8.45 (d, 1H), 8.88 (d, 1H). LCMS(ESI): $[M+H]^+$ m/z: calcd 460.13; found 460.0.

#### Synthesis of N,N-dimethyl-5-(3-(2-methyl-4-(5-(trifluoromethyl)-1,2,4-oxadiazol-3-yl)phenoxy)pyrrolidin-1-yl)isoxazole-3-carboxamide (Enantiomers 33 and 34)

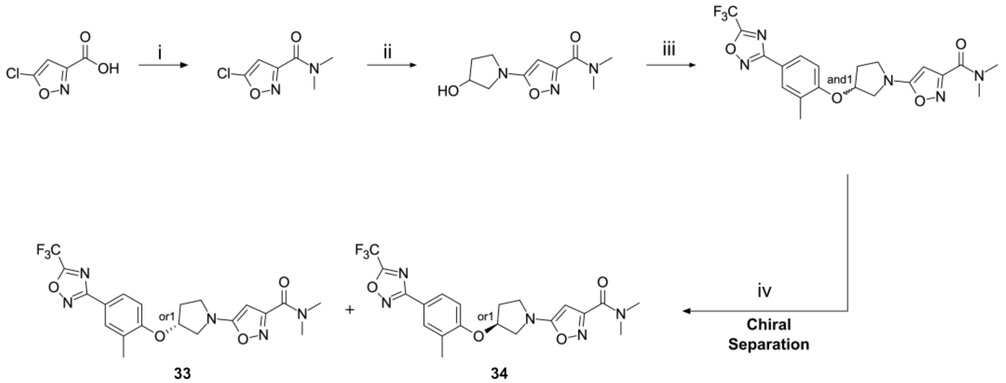


Scheme S9: Reagents and conditions: (i) CDI, N-methylmethanamine, triethylamine, RT, 16h (ii) pyrrolidin-3-ol, $K_2CO_3$, DMF, 90 °C, 16h (iii) 2-methyl-4-[5-(trifluoromethyl)-1,2,4-oxadiazol-3-yl]phenol, DIAD, $PPh_3$, THF, RT, 24h (iv) chiral HPLC

#### Step i: Data for 5-chloro-N,N-dimethyl-isoxazole-3-carboxamide

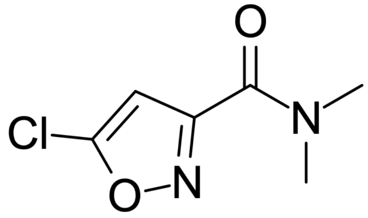

CDI (687 mg, 4.24 mmol) was added to a solution of 5-chloroisoxazole-3-carboxylic acid (500 mg, 3.39 mmol) in THF (5.0 mL) at room temperature. The resulting mixture was stirred at room temperature for 2 hr. $Et_3N$ (567 $\mu$L) and N-methylmethanamine (153 mg, 1.87 mmol, 235 $\mu$L, HCl) were added to the mixture. The resulting mixture was stirred at room temperature for 16 hr. The reaction mixture was concentrated under reduced pressure. The residue was subjected to reverse phase HPLC (0-1.3-5.3 min., 20-20-70% water – MeOH, +0.1% vol. of 25% aq. $NH_3$, flow: 30 mL/min, column: Chromatorex 18 SMB100-5T 100×19 mm, 5 $\mu$m) to afford 5-chloro-N,N-dimethyl-isoxazole-3-carboxamide (100 mg, 573 $\mu$mol, 16.9% yield) as a brown oil. LCMS(ESI): $[M+H]^+$ m/z: calcd 175.02; found 175.0.

#### Step ii: Data for 5-(3-hydroxypyrrolidin-1-yl)-N,N-dimethyl-isoxazole-3-carboxamide

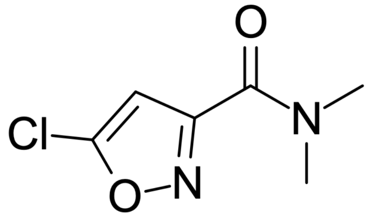


$K_2CO_3$ (158 mg, 1.15 mmol) was added to a stirred solution of 5-chloro-N,N-dimethyl-isoxazole-3-carboxamide (100 mg, 573 $\mu$mol) and pyrrolidin-3-ol (74.9 mg, 859 $\mu$mol, 71.4 $\mu$L) in DMF (4.0 mL) at room temperature. The resulting mixture was stirred at 90 °C for 16 hr. The reaction mixture was cooled to room temperature and filtered. The filtrate was concentrated under reduced pressure. The residue was subjected to reverse phase HPLC (0-1.3-5.3 min., 10-10-60% water – MeOH, flow: 30 mL/min, column: Chromatorex 18 SMB100-5T 100×19 mm, 5 $\mu$m) to afford 5-(3-hydroxypyrrolidin-1-yl)-N,N-dimethyl-isoxazole-3-carboxamide (76.0 mg, 337 $\mu$mol, 58.9% yield) as a yellow oil. $^1$H NMR (500 MHz, dmso) $\delta_H$ 1.85 – 1.92 (m, 1H), 1.96 – 2.05 (m, 1H), 2.95 (s, 3H), 3.05 (s, 3H), 3.22 (d, 1H), 3.38 – 3.49 (m, 3H), 4.36 – 4.40 (m, 1H), 5.16 (s, 1H). LCMS(ESI): $[M+H]^+$ m/z: calcd 226.13; found 226.2.

#### Step iii: Data for N,N-dimethyl-5-[3-[2-methyl-4-[5-(trifluoromethyl)-1,2,4-oxadiazol-3-yl]phenoxy]pyrrolidin-1-yl]isoxazole-3-carboxamide

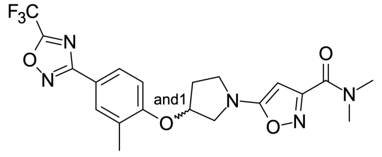


DIAD (75.1 mg, 371 $\mu$mol, 73.1 $\mu$L) and $PPh_3$ (97.4 mg, 371 $\mu$mol) were to a stirred solution of 5-(3-hydroxypyrrolidin-1-yl)-N,N-dimethyl-isoxazole-3-carboxamide (76.0 mg, 337 $\mu$mol) and 2-methyl-4-[5-(trifluoromethyl)-1,2,4-oxadiazol-3-yl]phenol (90.6 mg, 371 $\mu$mol) in THF (5.0 mL) added at room temperature. The resulting mixture was stirred at room temperature for 24 hr. The reaction mixture was concentrated under reduced pressure. The residue was subjected to reverse phase HPLC (0-1.3-6.3 min., 50-55-95% water – MeOH, flow: 30 mL/min, column: Chromatorex 18 SMB100-5T 100×19 mm, 5 $\mu$m) to afford N,N-dimethyl-5-[3-[2-methyl-4-[5-(trifluoromethyl)-1,2,4-oxadiazol-3-yl]phenoxy]pyrrolidin-1-yl]isoxazole-3-carboxamide (50.0 mg, 111 $\mu$mol, 32.8% yield) as a colorless gum. LCMS(ESI): $[M+H]^+$ m/z: calcd 452.17; found 452.0.

#### Step iv: Chiral resolution

N,N-Dimethyl-5-[3-[2-methyl-4-[5-(trifluoromethyl)-1,2,4-oxadiazol-3-yl]phenoxy]pyrrolidin-1-yl]isoxazole-3-carboxamide (50.0 mg, 114 $\mu$mol) was subjected to chiral HPLC (column: CHIRALPAK IA (250×20 mm, 5 $\mu$m); eluent: Hexane-IPA-MeOH, 50:25:25; flow: 13 mL/min) to afford **33** (14.0 mg, 31.9 $\mu$mol, 28.0% yield) and (**34** (15.0 mg, 34.1 $\mu$mol, 30.0% yield) as yellow oils.

#### Data for (R,or1)-N,N-dimethyl-5-(3-(2-methyl-4-(5-(trifluoromethyl)-1,2,4-oxadiazol-3-yl)phenoxy)pyrrolidin-1-yl)isoxazole-3-carboxamide (Compound 33)

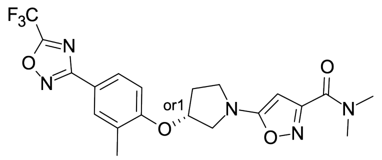


Analytical data: $^1$H NMR (500 MHz, dmso) $\delta_H$ 2.16 (s, 3H), 2.19 – 2.26 (m, 1H), 2.29 – 2.40 (m, 1H), 2.94 (s, 3H), 3.03 (s, 3H), 3.49 – 3.62 (m, 3H), 3.78 (dd, 1H), 5.26 (s, 1H), 5.28 – 5.32 (m, 1H), 7.24 (d, 1H), 7.86 (s, 1H), 7.89 (d, 1H).

LCMS(ESI): $[M+H]^+$ m/z: calcd 452.17; found 452.0. Analytical chiral HPLC (column: Chiralpak IA (250 × 4.6 mm, 5 μm; mobile phase: Hexane-IPA-MeOH, 50:25:25; flow 0.6 mL/min.): Rt 22.09 min.

#### Data for (S,or1)-N,N-dimethyl-5-(3-(2-methyl-4-(5-(trifluoromethyl)-1,2,4-oxadiazol-3-yl)phenoxy)pyrrolidin-1-yl)isoxazole-3-carboxamide (Compound 34)

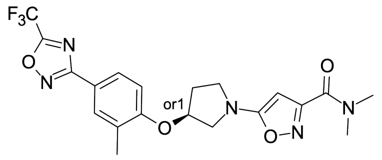

Analytical data: $^1$H NMR (500 MHz, dmso) $\delta_H$ 2.16 (s, 3H), 2.19 – 2.26 (m, 1H), 2.29 – 2.40 (m, 1H), 2.94 (s, 3H), 3.03 (s, 3H), 3.49 – 3.62 (m, 3H), 3.78 (dd, 1H), 5.26 (s, 1H), 5.28 – 5.32 (m, 1H), 7.24 (d, 1H), 7.86 (s, 1H), 7.89 (d, 1H). LCMS(ESI): $[M+H]^+$ m/z: calcd 452.17; found 452.0. Analytical chiral HPLC (column: Chiralpak IA (250 × 4.6 mm, 5 μm; mobile phase: Hexane-IPA-MeOH, 50:25:25; flow 0.6 mL/min.): Rt 34.50 min.

#### 3-[3-methyl-4-[3-(1,3,4-oxadiazol-2-yl)propoxy]phenyl]-5-(trifluoromethyl)-1,2,4-oxadiazole (Compound 35)

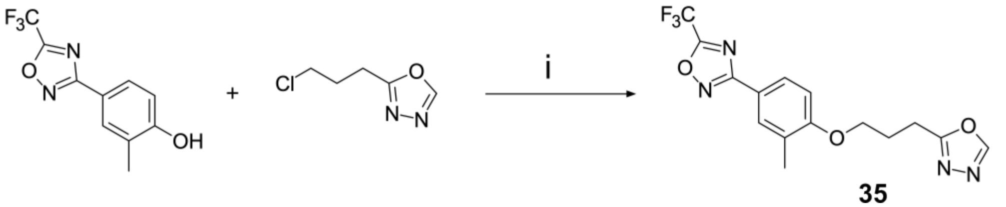

35

Scheme S10: Reagents and conditions: (i) $^t$BuOK, NaI, DMF, RT, 24h

#### Step i: Data for 3-[3-methyl-4-[3-(1,3,4-oxadiazol-2-yl)propoxy]phenyl]-5-(trifluoromethyl)-1,2,4-oxadiazole (Compound 35)

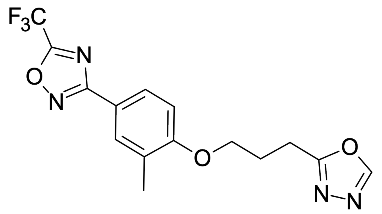


NaI (93.0 mg, 620 μmol), ᵗBuOK (83.5 mg, 744 μmol) and 2-(3-chloropropyl)-1,3,4-oxadiazole (100 mg, 682 μmol) were added sequentially to a solution of 2-methyl-4-[5-(trifluoromethyl)-1,2,4-oxadiazol-3-yl]phenol (151 mg, 620 μmol) in DMF (1.0 mL). The resulting mixture was stirred at room temperature for 24 hr. The reaction mixture was subjected to HPLC (0-1.2-6 min, 30-30-80% water+FA (0.1% vol.) - ACN+FA (0.1% vol.); flow: 30 mL/min, column: Chromatorex 18 SMB100-5T, 100×19 mm, 5 μm) to afford 3-[3-methyl-4-[3-(1,3,4-oxadiazol-2-yl)propoxy]phenyl]-5-(trifluoromethyl)-1,2,4-oxadiazole (2.20 mg, 5.90 μmol, 0.95% yield) as a brown solid. $^1$H NMR (500 MHz, CDCl3) $\delta_H$ 2.26 (s, 3H), 2.40 (p, 2H), 3.17 (t, 2H), 4.18 (t, 2H), 6.90 (d, 1H), 7.89 (s, 1H), 7.92 (d, 1H), 8.36 (s, 1H). LCMS(ESI): $[M+H]^+$ m/z: calcd 355.11; found 355.0.

#### Synthesis of 2-methyl-5-[3-[2-methyl-4-[5-(trifluoromethyl)-1,2,4-oxadiazol-3-yl]phenoxy]propyl]-1,3,4-oxadiazole (Compound 36)

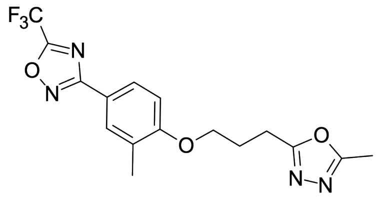

**36** was synthesised by a route similar to Compound **2**: DIAD (99.4 mg, 492 $\mu$mol) was added dropwise to a stirred solution of 3-(5-methyl-1,3,4-oxadiazol-2-yl)propan-1-ol (58.2 mg, 410 $\mu$mol), 2-methyl-4-[5-(trifluoromethyl)-1,2,4-oxadiazol-3-yl]phenol (100 mg, 410 $\mu$mol) and $PPh_3$ (129 mg, 492 $\mu$mol) in anhydrous THF (5.0 mL). The resulting mixture was stirred at room temperature for 16 hr. The reaction mixture was concentrated under reduced pressure. The residue was subjected to HPLC (0-1.5-6 min, 30-30-85% water+FA (0.1% vol.) - ACN+FA (0.1% vol.); flow: 30 mL/min, column: Chromatorex 18 SMB100-5T, 100×19 mm, 5 $\mu$m) to afford 2-methyl-5-[3-[2-methyl-4-[5-(trifluoromethyl)-1,2,4-oxadiazol-3-yl]phenoxy]propyl]-1,3,4-oxadiazole (45.0 mg, 116 $\mu$mol, 28.3% yield) as a yellow gum. $^1$H NMR (500 MHz, dmso) $\delta_H$ 2.15 – 2.24 (m, 5H), 2.42 (s, 3H), 3.01 (t, 2H), 4.16 (t, 2H), 7.13 (d, 1H), 7.83 (d, 1H), 7.87 (dd, 1H). LCMS(ESI): $[M+H]^+$ m/z: calcd 369.13; found 369.2.

#### Synthesis of 2-(difluoromethyl)-5-[3-[2-methyl-4-[5-(trifluoromethyl)-1,2,4-oxadiazol-3-yl]phenoxy]propyl]-1,3,4-oxadiazole (Compound 37)

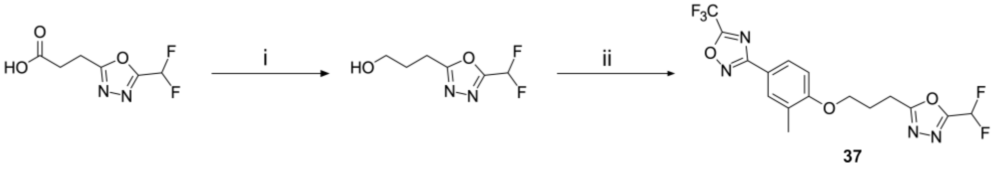

Scheme S11: Reagents and conditions: (i) borane dimethyl sulfide complex, THF, RT, 2h (ii) 2-methyl-4-[5-(trifluoromethyl)-1,2,4-oxadiazol-3-yl]phenol, $PPh_3$, DIAD, THF, RT, 24h

#### Step i: Data for 3-[5-(difluoromethyl)-1,3,4-oxadiazol-2-yl]propan-1-ol

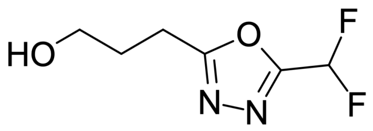


Borane dimethyl sulfide complex (158 mg, 2.08 mmol, 198 $\mu$L) was added in one portion to a stirred solution of 3-[5-(difluoromethyl)-1,3,4-oxadiazol-2-yl]propanoic acid (200 mg, 1.04 mmol) in THF (10 mL). The resulting mixture was stirred at room temperature for 2 hr. The reaction mixture was diluted with MeOH (2.0 mL), heated for 10 min and concentrated under reduced pressure. The residue was subjected to HPLC (0-1-5 min., 5-5-45% water – MeOH, flow: 30 mL/min, column: Chromatorex 18 SMB100-5T 100×19 mm, 5 $\mu$m) to afford 3-[5-(difluoromethyl)-1,3,4-oxadiazol-2-yl]propan-1-ol (24.6 mg, 138 $\mu$mol, 13.3% yield) as a white oil. LCMS(ESI): $[M+H]^+$ m/z: calcd 179.07; found 179.0.

#### Step ii: Data for 2-(difluoromethyl)-5-[3-[2-methyl-4-[5-(trifluoromethyl)-1,2,4-oxadiazol-3-yl]phenoxy]propyl]-1,3,4-oxadiazole (Compound 37)

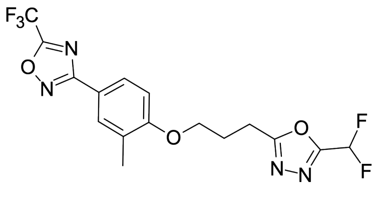


$PPh_3$ (43.5 mg, 166 $\mu$mol), DIAD (33.5 mg, 166 $\mu$mol) and 2-methyl-4-[5-(trifluoromethyl)-1,2,4-oxadiazol-3-yl]phenol (33.7 mg, 138 $\mu$mol) were added to a solution of 3-[5-(difluoromethyl)-1,3,4-oxadiazol-2-yl]propan-1-ol (24.6 mg, 138 $\mu$mol) in THF (5.0 mL). The resulting mixture was stirred at room temperature for 15 hr. The reaction mixture was subjected to HPLC (0-5 min., 40-90% water – ACN, flow: 30 mL/min, column: Chromatorex 18A31 SMB100-5T 100×19 mm, 5 $\mu$m) to afford 2-(difluoromethyl)-5-[3-[2-methyl-4-[5-(trifluoromethyl)-1,2,4-oxadiazol-3-yl]phenoxy]propyl]-1,3,4-oxadiazole (27.0 mg, 66.8 $\mu$mol, 48.4% yield) as a white solid. $^1$H NMR (500 MHz, dmso) $\delta_H$ 2.15 (s, 3H), 2.26 (p, 2H), 3.16 (t, 2H), 4.19 (t, 2H),

7.13 (d, 1H), 7.45 (t, 1H, CHF2), 7.83 (d, 1H), 7.85 – 7.89 (m, 1H). LCMS(ESI): $[M+H]^+$ m/z: calcd 405.11; found 405.2.

#### Synthesis of 2-[3-[2-methyl-4-[5-(trifluoromethyl)-1,2,4-oxadiazol-3-yl]phenoxy]propyl]-5-(trifluoromethyl)-1,3,4-oxadiazole (Compound 38)

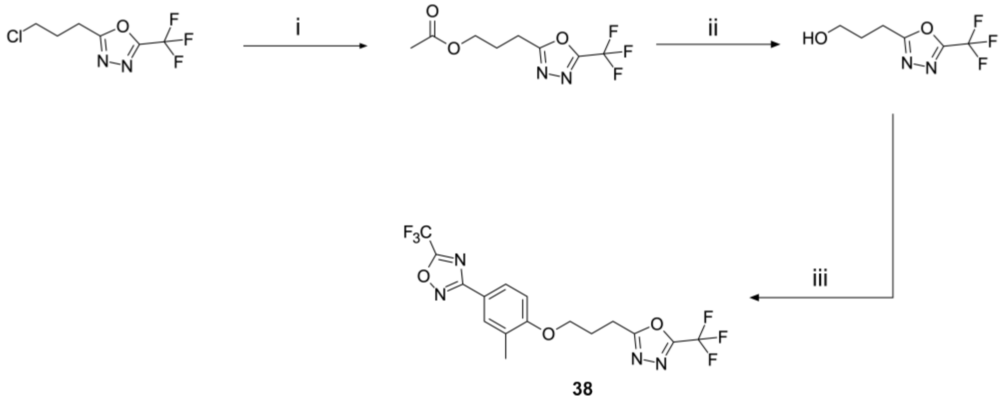


Scheme S12: Reagents and conditions: (i) KOAc, DMF, 60 °C, 16h (ii) NaOH, EtOH, $H_2O$, THF, 0 °C to RT, 16h (iii) 2-methyl-4-[5-(trifluoromethyl)-1,2,4-oxadiazol-3-yl]phenol, $PPh_3$, DIAD, THF, RT, 36h

#### Step i: Data for 3-[5-(trifluoromethyl)-1,3,4-oxadiazol-2-yl]propyl acetate

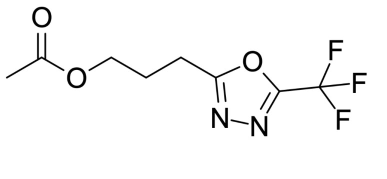

AcOK (1.14 g, 11.7 mmol) was added to a solution of 2-(3-chloropropyl)-5-(trifluoromethyl)-1,3,4-oxadiazole (500 mg, 2.33 mmol) in DMF (3.0 mL). The resulting mixture was stirred at 60 °C for 16 hr. The reaction mixture was cooled to room temperature, quenched with water (10 mL) and extracted with EtOAc (3×20 mL). The combined organic layers were washed with brine, dried over anhydrous sodium sulfate and concentrated under reduced pressure to afford 3-[5-(trifluoromethyl)-1,3,4-oxadiazol-2-

yl]propyl acetate (400 mg, 1.68 mmol, 72.1% yield) as a yellow oil which was used in the next step without further purification. LCMS(ESI): $[M+H]^+$ m/z: calcd 239.07; found 239.2.

#### Step ii: Data for 3-[5-(trifluoromethyl)-1,3,4-oxadiazol-2-yl]propan-1-ol

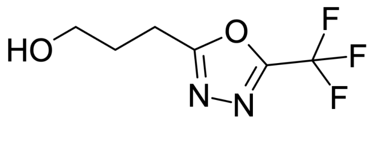


A solution of NaOH (92.4 mg, 2.31 mmol) in EtOH (2.0 mL) and water (500 $\mu$L) was added to a solution of 3-[5-(trifluoromethyl)-1,3,4-oxadiazol-2-yl]propyl acetate (500 mg, 2.10 mmol) in THF (2.0 mL) at 0 °C. The resulting mixture was stirred at room temperature overnight. The reaction mixture was concentrated under reduced pressure. The residue was extracted with EtOAc (2×10 mL). The combined organic layers were dried over anhydrous sodium sulfate and concentrated under reduced pressure to afford 3-[5-(trifluoromethyl)-1,3,4-oxadiazol-2-yl]propan-1-ol (150 mg, 765 $\mu$mol, 36.4% yield) as a yellow oil which was used in next step without further purification. LCMS(ESI): $[M+H]^+$ m/z: calcd 197.06; found 197.0.

#### Step iii: Data for 2-[3-[2-methyl-4-[5-(trifluoromethyl)-1,2,4-oxadiazol-3-yl]phenoxy]propyl]-5-(trifluoromethyl)-1,3,4-oxadiazole (Compound 38)

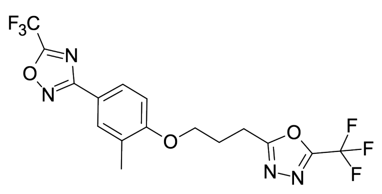


DIAD (186 mg, 918 $\mu$mol, 181 $\mu$L) was added dropwise to a stirred solution of 3-[5-(trifluoromethyl)-1,3,4-oxadiazol-2-yl]propan-1-ol (150 mg, 765 $\mu$mol), 2-methyl-4-[5-(trifluoromethyl)-1,2,4-oxadiazol-3-yl]phenol (187 mg, 765 $\mu$mol) and $PPh_3$ (241 mg, 918 $\mu$mol) in anhydrous THF (1.47 mL). The resulting mixture was stirred at room temperature for 36 hr. The reaction mixture was cooled to room temperature and concentrated under reduced pressure. The residue was subjected to HPLC (0-1.5-5.3 min., 50-50-80% water – ACN, +0.1% vol. of 25% aq. $NH_3$, flow: 30 mL/min, column: SunFire 100×19 mm, 5 $\mu$m) to afford 2-[3-[2-methyl-4-[5-(trifluoromethyl)-1,2,4-oxadiazol-3-yl]phenoxy]propyl]-5-(trifluoromethyl)-1,3,4-oxadiazole (2.00 mg, 4.74 $\mu$mol, 0.62% yield) as a colorless gum. LCMS(ESI): $[M+H]^+$ m/z: calcd 423.1; found 423.0.

#### Synthesis of 3-[3-methyl-4-[3-(5-methyl-4H-1,2,4-triazol-3-yl)propoxy] phenyl]-5-(trifluoromethyl)-1,2,4-oxadiazole (Compound 39)

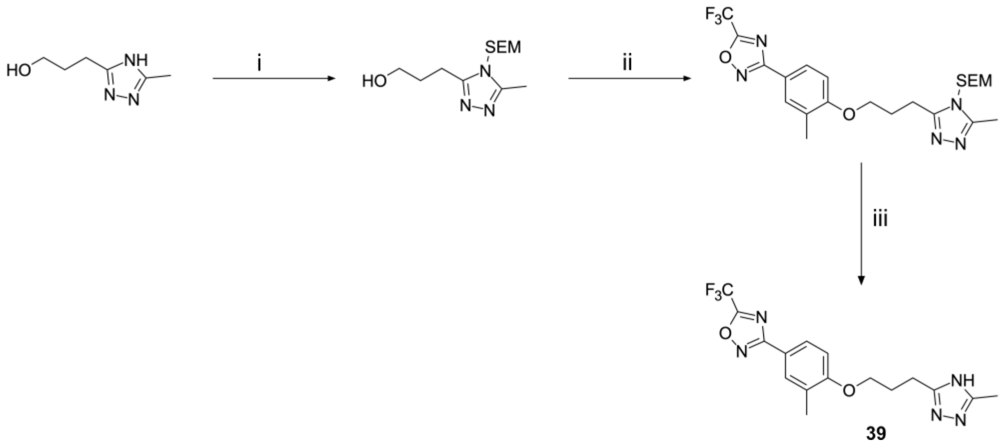


Scheme S13: Reagents and conditions: (i) SEM-Cl, $Et_3N$, DMF, RT, 16h (ii) 2-methyl-4-[5-(trifluoromethyl)-1,2,4-oxadiazol-3-yl]phenol, $PPh_3$, DIAD, THF, 0 °C to RT, 16h (iii) TFA, DCM, 0 °C to RT, 16h

#### Step i: Data for 3-[5-methyl-4-(2-trimethylsilylethoxymethyl)-1,2,4-triazol-3-yl]propan-1-ol

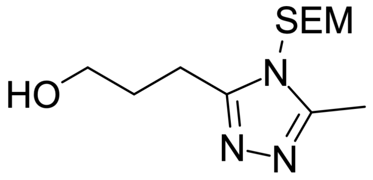


3-(5-Methyl-4H-1,2,4-triazol-3-yl)propan-1-ol (400 mg, 2.83 mmol), 2-(chloromethoxy)ethyl-trimethyl-silane (543 mg, 3.26 mmol) and $Et_3N$ (287 mg, 2.83 mmol, 395 $\mu$L) were mixed in DMF (10 mL). The resulting mixture was

stirred at room temperature for 16 hr. The reaction mixture was diluted with DCM (20 mL) and washed with water (20 mL). The organic layer was separated and concentrated under reduced pressure to afford 3-[5-methyl-4-(2-trimethylsilylethoxymethyl)-1,2,4-triazol-3-yl]propan-1-ol (300 mg, 674 $\mu$mol, 23.8% yield) as a yellow oil which was used in the next step without further purification. LCMS(ESI): $[M+H]^+$ m/z: calcd 272.18; found 272.2.

##### Step ii: Data for trimethyl-[2-[[3-methyl-5-[3-[2-methyl-4-[5-(trifluoromethyl)-1,2,4-oxadiazol-3-yl]phenoxy]propyl]-1,2,4-triazol-4-yl]methoxy]ethyl]silane

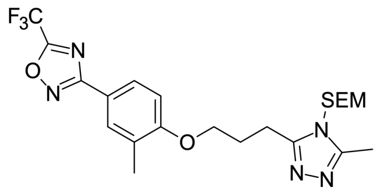


DIAD (72.4 mg, 358 $\mu$mol, 70.5 $\mu$L) was added dropwise to a stirred solution of 2-methyl-4-[5-(trifluoromethyl)-1,2,4-oxadiazol-3-yl]phenol (72.9 mg, 298 $\mu$mol), 3-[5-methyl-4-(2-trimethylsilylethoxymethyl)-1,2,4-triazol-3-yl]propan-1-ol (300 mg, 298 $\mu$mol) and $PPh_3$ (93.9 mg, 358 $\mu$mol) in anhydrous THF (10 mL) at 0 °C. The resulting mixture was stirred at room temperature for 16 hr. The resulting mixture was concentrated under reduced pressure. The residue was partitioned between EtOAc (25 mL) and water (7 mL). The organic layer was separated, dried over anhydrous sodium sulfate and concentrated under reduced pressure to afford trimethyl-[2-[[3-methyl-5-[3-[2-methyl-4-[5-(trifluoromethyl)-1,2,4-oxadiazol-3-yl]phenoxy]propyl]-1,2,4-triazol-4-yl]methoxy]ethyl]silane (600 mg, 265 $\mu$mol, 88.9% yield) as a brown oil which was used in next step without further purification. LCMS(ESI): $[M+H]^+$ m/z: calcd 498.22; found 498.0.

#### Step iii: Data for 3-[3-methyl-4-[3-(5-methyl-4H-1,2,4-triazol-3-yl)propoxy] phenyl]-5-(trifluoromethyl)-1,2,4-oxadiazole (Compound 39)

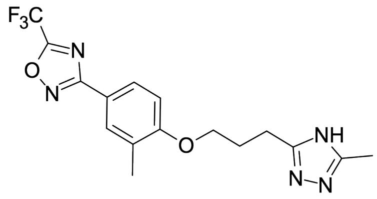

TFA (182 mg, 1.59 mmol, 122 $\mu$L) was added to a solution of trimethyl-[2-[[3-methyl-5-[3-[2-methyl-4-[5-(trifluoromethyl)-1,2,4-oxadiazol-3-yl]phenoxy]propyl]-1,2,4-triazol-4-yl]methoxy]ethyl]silane (600 mg, 265 $\mu$mol) in DCM (10 mL) at 0 °C. The resulting mixture was stirred at room temperature for 16 hr. The reaction mixture was concentrated under reduced pressure. The residue was subjected to HPLC (0-1.3-5.3 min., 40-40-65% water – ACN, flow: 30 mL/min, column: Chromatorex 18 SMB100-5T 100×19 mm, 5 $\mu$m) to afford 3-[3-methyl-4-[3-(5-methyl-4H-1,2,4-triazol-3-yl)propoxy]phenyl]-5-(trifluoromethyl)-1,2,4-oxadiazole (22.5 mg, 58.2 $\mu$mol, 21.9% yield) as a white solid. $^1$H NMR (500 MHz, dmso) $\delta_H$ 2.05 - 2.26 (m, 8H), 2.69 – 2.87 (m, 2H), 4.11 (t, 2H), 7.11 (d, 1H), 7.82 – 7.88 (m, 2H), 13.15 – 2.27 (m, 1H). LCMS(ESI): $[M+H]^+$ m/z: calcd 368.15; found 368.2.

#### Synthesis of 3-[3-methyl-4-[3-(1H-pyrazol-4-yl)propoxy]phenyl] -5-(trifluoromethyl)-1,2,4-oxadiazole (Compound 40)

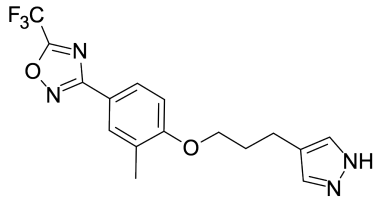

**40** was synthesised by a route similar to Compound **2**: DIAD (96.2 mg, 476 $\mu$mol, 93.6 $\mu$L), $PPh_3$ (125 mg, 476 $\mu$mol) and 2-methyl-4-[5-(trifluoromethyl)-1,2,4-oxadiazol-3-yl]phenol (96.8 mg, 396 $\mu$mol) were added to a solution of 3-(1H-pyrazol-4-yl)propan-1-ol (50.0 mg, 396 $\mu$mol) in THF (5.0 mL). The resulting mixture was stirred at room temperature for 14 hr. The reaction mixture was subjected to HPLC (0-1-5 min., 55-55-80% water – ACN, flow: 30 mL/min, column:

Chromatorex 18 SMB100-5T 100×19 mm, 5 μm) to afford 3-[3-methyl-4-[3-(1H-pyrazol-4-yl)propoxy]phenyl]-5-(trifluoromethyl)-1,2,4-oxadiazole (38.3 mg, 109 μmol, 27.4% yield) as a beige solid. $^1$H NMR (500 MHz, dmso) $\delta_H$ 1.96 – 2.06 (m, 2H), 2.24 (s, 3H), 2.61 (t, 2H), 4.06 (t, 2H), 7.11 (d, 1H), 7.32 (br. s, 1H), 7.52 (br. s, 1H), 7.82 – 7.89 (m, 2H), 12.52 (br. s, 1H). LCMS(ESI): $[M+H]^+$ m/z: calcd 353.14; found 353.2.

#### Synthesis of 3-[3-methyl-4-[3-(1-methylpyrazol-4-yl)propoxy]phenyl]-5-(trifluoromethyl)-1,2,4-oxadiazole (Compound 41)

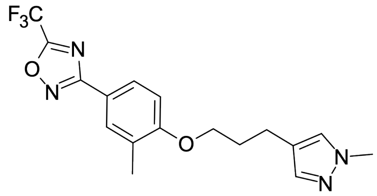

**41** was synthesised by a route similar to Compound **2**: 2-Methyl-4-[5-(trifluoromethyl)-1,2,4-oxadiazol-3-yl]phenol (50.0 mg, 205 μmol) was mixed with 3-(1-methylpyrazol-4-yl)propan-1-ol (28.7 mg, 205 μmol) and $PPh_3$ (80.6 mg, 307 μmol) in dry THF (3.0 mL), after that DEAD (53.5 mg, 307 μmol) was added dropwise to the solution at 0 °C. The resulting mixture was allowed to warm up to room temperature and stirred overnight. The reaction mixture was subjected to HPLC (0-1-6 min., 40-40-90% water – ACN, flow: 60 mL/min, column: XBridge C18 OBD 100×30 mm, 5 μm) to afford 3-[3-methyl-4-[3-(1-methylpyrazol-4-yl)propoxy]phenyl]-5-(trifluoromethyl)-1,2,4-oxadiazole (65.4 mg, 179 μmol, 87.2% yield) as a colorless oil. $^1$H NMR (500 MHz, dmso) $\delta_H$ 1.93 – 2.08 (m, 2H), 2.23 (s, 3H), 2.57 (t, 2H), 3.74 (s, 3H), 4.06 (t, 2H), 7.11 (d, 1H), 7.24 (s, 1H), 7.48 (s, 1H), 7.82 – 7.88 (m, 2H). LCMS(ESI): $[M+H]^+$ m/z: calcd 367.16; found 367.0.

#### Synthesis of 3-[3-methyl-4-[3-(2-pyridyl)propoxy]phenyl]-5-(trifluoromethyl)-1,2,4-oxadiazole)-1,2,4-oxadiazole (Compound 42)

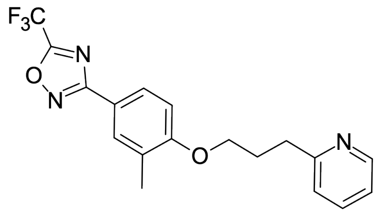


**42** was synthesised by a route similar to Compound **2**: $PPh_3$ (107 mg, 410 $\mu$mol) and DIAD (82.8 mg, 410 $\mu$mol) were added to a stirred solution of 2-methyl-4-[5-(trifluoromethyl)-1,2,4-oxadiazol-3-yl]phenol (100 mg, 410 $\mu$mol) and 3-(2-pyridyl)propan-1-ol (51.1 mg, 372 $\mu$mol) in THF (4.0 mL) at room temperature. The resulting mixture was stirred at room temperature for 24 hr. The reaction mixture was concentrated under reduced pressure. The residue was subjected to HPLC (0-1.3-5.3 min., 60-60-85% water – ACN, flow: 30 mL/min, column: Chromatorex 18 SMB100-5T 100×19 mm, 5 $\mu$m) to afford 3-[3-methyl-4-[3-(2-pyridyl)propoxy]phenyl]-5-(trifluoromethyl)-1,2,4-oxadiazole (6.60 mg, 18.2 $\mu$mol, 4.88% yield) as a white solid. $^{1}$H NMR (500 MHz, dmso) $\delta_H$ 2.13 – 2.20 (m, 2H), 2.21 (s, 3H), 2.92 (t, 2H), 4.11 (t, 2H), 7.11 (d, 1H), 7.19 (dd, 1H), 7.28 (d, 1H), 7.65 – 7.72 (m, 1H), 7.81 – 7.88 (m, 2H), 8.48 (dd, 1H). LCMS(ESI): $[M+H]^+$ m/z: calcd 364.15; found 364.2.

#### Synthesis of 3-[3-methyl-4-[3-(3-pyridyl)propoxy]phenyl]-5-(trifluoromethyl)-1,2,4-oxadiazole (Compound 43)

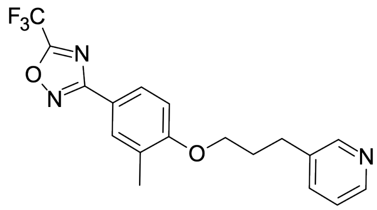


**43** was synthesised by a route similar to Compound **2**: DIAD (95.2 mg, 471 $\mu$mol, 92.7 $\mu$L), $PPh_3$ (124 mg, 471 $\mu$mol) and 3-(3-pyridyl)propan-1-ol (64.6 mg, 471 $\mu$mol) were added to a solution of 2-methyl-4-[5-(trifluoromethyl)-1,2,4-oxadiazol-3-yl]phenol (100 mg, 410 $\mu$mol) in THF (15.0 mL) at 0 °C. The resulting mixture was stirred at room temperature for 16 hr. The reaction mixture was concentrated under reduced pressure. The residue was subjected to HPLC (0-1-5 min, 35-35-55% water+FA (0.1% vol.) - ACN+FA (0.1% vol.); flow: 30 mL/min, column: Chromatorex 18 SMB100-5T, 100×19 mm, 5 $\mu$m) to afford 3-[3-methyl-4-[3-(3-pyridyl)propoxy]phenyl]-5-(trifluoromethyl)-1,2,4-oxadiazole (20.0 mg, 55.1 $\mu$mol, 13.4% yield) as an off-white solid. $^1$H NMR (500 MHz, dmso) $\delta_H$ 1.96 – 2.06 (m, 2H), 2.22 (s, 3H), 2.80 (t, 2H), 4.07 (t, 2H), 7.11 (d, 1H), 7.30 (dd, 1H), 7.63 – 7.69 (m, 1H), 7.82 – 7.87 (m, 2H), 8.39 (dd, 1H), 8.45 (d, 1H). LCMS(ESI): $[M+H]^+$ m/z: calcd 364.15; found 364.2.

#### Synthesis of 3-[3-methyl-4-[3-(4-pyridyl)propoxy]phenyl]-5-(trifluoromethyl)-1,2,4-oxadiazole (Compound 44)

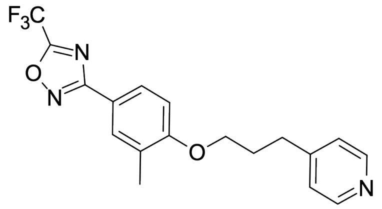


**44** was synthesised by a route similar to Compound **2**: 2-Methyl-4-[5-(trifluoromethyl)-1,2,4-oxadiazol-3-yl]phenol (50.0 mg, 205 $\mu$mol) was mixed with 3-(4-pyridyl)propan-1-ol (28.1 mg, 205 $\mu$mol) and $PPh_3$ (80.6 mg, 307 $\mu$mol) in dry THF (3.0 mL), after that DEAD (53.5 mg, 307 $\mu$mol) was added dropwise to the solution at 0 °C. The resulting mixture was allowed to warm up to room temperature and stirred overnight. The reaction mixture was subjected to HPLC (0-1-6 min., 40-40-90% water – ACN, flow: 60 mL/min, column: XBridge C18 OBD 100×30 mm, 5 $\mu$m) to afford 3-[3-methyl-4-[3-(4-pyridyl)propoxy]phenyl]-5-(trifluoromethyl)-1,2,4-oxadiazole (45.0 mg, 124 $\mu$mol, 60.5% yield) as a beige solid. $^1$H NMR (500 MHz, dmso) $\delta_H$ 2.10 (p, 2H), 2.21 (s, 3H), 2.79 (t, 2H), 4.07 (t, 2H), 7.11 (d, 1H), 7.26 (d, 2H), 7.82 – 7.87 (m, 2H), 8.44 (d, 2H). LCMS(ESI): $[M+H]^+$ m/z: calcd 364.15; found 364.0.

#### Synthesis of 3-[3-methyl-4-[3-(1-phenyltetrazol-5-yl)propoxy]phenyl]-5-(trifluoromethyl)-1,2,4-oxadiazole (Compound 45)

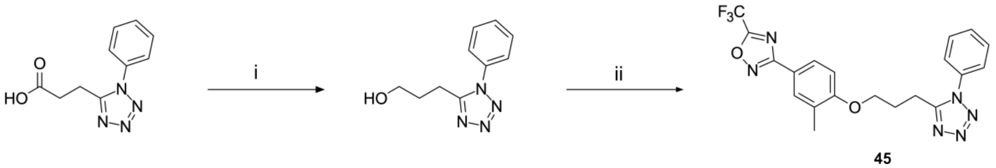


Scheme S14: Reagents and conditions: (i) borane dimethyl sulfide complex, THF, 10 °C to RT, 16h (ii) $PPh_3$, DIAD, 2-methyl-4-[5-(trifluoromethyl)-1,2,4-oxadiazol-3-yl]phenol, THF, RT, 16h

#### Step i: Data for 3-(1-phenyltetrazol-5-yl)propan-1-ol

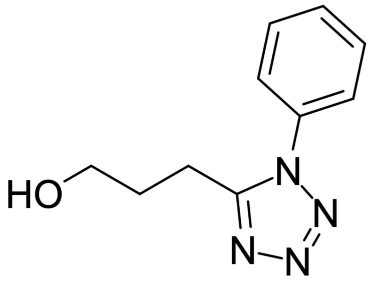


Borane dimethyl sulfide complex (69.6 mg, 917 $\mu$mol, 86.9 $\mu$L) was added to a pre-cooled to 0 °C stirred solution of 3-(1-phenyltetrazol-5-yl)propanoic acid (100 mg, 458 $\mu$mol) in THF (1.0 mL) keeping temperature below 10 °C. The resulting mixture was allowed to warm up to room temperature and stirred for 16 hr. The reaction mixture was concentrated under reduced pressure. The residue was dissolved in MeOH (2.0 mL) and concentrated under reduced pressure again to afford 3-(1-phenyltetrazol-5-yl)propan-1-ol (100 mg, crude) as a yellow oil which was used in the next step without further purification. LCMS(ESI): $[M+H]^+$ m/z: calcd 205.11; found 205.2.

#### Step ii: Data for 3-[3-methyl-4-[3-(1-phenyltetrazol-5-yl)propoxy]phenyl]-5-(trifluoromethyl)-1,2,4-oxadiazole (Compound 45)

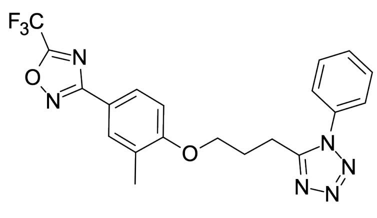

DIAD (218 mg, 1.08 mmol) was added dropwise to a stirred solution of 3-(1-phenyltetrazol-5-yl)propan-1-ol (200 mg, 979 $\mu$mol), 2-methyl-4-[5-(trifluoromethyl)-1,2,4-oxadiazol-3-yl]phenol (191 mg, 783 $\mu$mol) and $PPh_3$ (283 mg, 1.08 mmol) in anhydrous THF (1.5 mL). The resulting mixture was stirred at room temperature for 16 hr. The reaction mixture was concentrated under reduced pressure. The residue was subjected to HPLC (0-55 min., 40-80% water – ACN, flow: 30 mL/min, column: PHENYL SMB100-5T 100×19 mm, 5 $\mu$m) to afford 3-[3-methyl-4-[3-(1-phenyltetrazol-5-yl)propoxy]phenyl]-5-(trifluoromethyl)-1,2,4-oxadiazole (41.0 mg, 95.3 $\mu$mol, 9.73% yield) as a light-yellow solid. $^1$H NMR (500 MHz, dmso) $\delta_H$ 2.06 (s, 3H), 2.19 (p, 2H), 3.10 (t, 2H), 4.10 (t, 2H), 7.06 (d, 1H), 7.56 – 7.66 (m, 5H), 7.80 (s, 1H), 7.84 (d, 1H). LCMS(ESI): $[M+H]^+$ m/z: calcd 431.16; found 431.2.

#### Synthesis of 5-[3-[(3-ethoxy-1,2-benzoxazol-6-yl)oxy]propyl]-N,N-dimethyl-isoxazole-3-carboxamide (Compound 46)

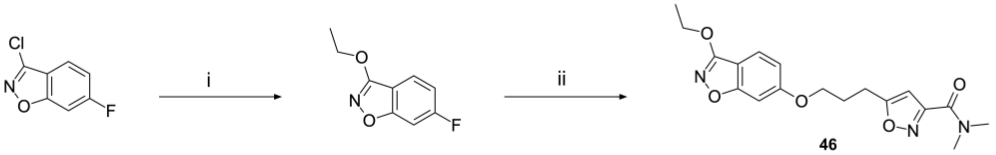

Scheme S15: Reagents and conditions: (i) NaH, EtOH, DMF, RT, 16h (ii) 5-(3-Hydroxypropyl)-N,N-dimethyl-isoxazole-3-carboxamide, NaH, DMF, 0 °C to RT, 16h

#### Step i: Data for 3-ethoxy-6-fluoro-1,2-benzoxazole

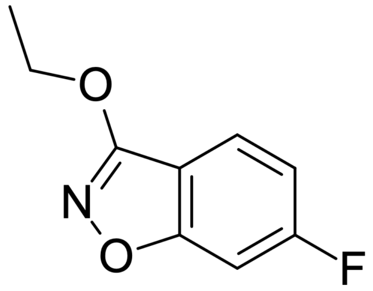

NaH (14.0 mg, 583 μmol, 60% dispersion in mineral oil) was added portionwise to an ice-cooled mixture of 3-Chloro-6-fluoro-1,2-benzoxazole (100 mg, 583 μmol) and EtOH (26.9 mg, 583 μmol, 34.0 μL) in DMF (2.0 mL). The resulting mixture was stirred at room temperature for 16 hr. The reaction mixture was cooled to room temperature to afford a solution of 3-ethoxy-6-fluoro-1,2-benzoxazole (100 mg, 254 μmol, 46% purity by LCMS) in DMF which was used in the next step without work up.

#### Step ii: Data for 5-[3-[(3-ethoxy-1,2-benzoxazol-6-yl)oxy]propyl]-N,N-dimethyl-isoxazole-3-carboxamide (Compound 46)

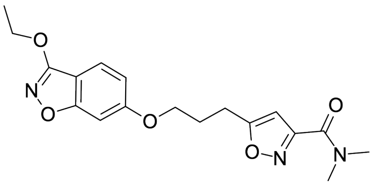

5-(3-Hydroxypropyl)-N,N-dimethyl-isoxazole-3-carboxamide (109 mg, 552 μmol) and NaH (13.3 mg, 552 μmol, 60% dispersion in mineral oil) were added sequentially to a stirred solution

of 3-ethoxy-6-fluoro-1,2-benzoxazole (100 mg, 552 $\mu$mol) in DMF (2.0 mL) at 0 °C. The resulting mixture was stirred at room temperature for 16 hr. The reaction mixture was partitioned between EtOAc (20 mL) and water (5.0 mL). The organic layer was separated, washed with water (3×5 mL) and brine (5 mL), dried over anhydrous sodium sulfate and concentrated under reduced pressure. The residue was subjected to HPLC (0-1.3-6 min., 35-35-45% water – ACN, +0.1% vol. of 25% aq. $NH_3$, flow: 30 mL/min, column: XBridge BEH C18 100×19 mm, 5 $\mu$m) to afford a mixture of two regioisomers which was further subjected to chiral HPLC (column: Chromatorex EP2 (100×19 mm, 5 $\mu$m), eluent: 95:2.5:2.5 Hexane:IPA:MeOH, flow: 12.0 mL/min) to afford 5-[3-[(3-ethoxy-1,2-benzoxazol-6-yl)oxy]propyl]-N,N-dimethyl-isoxazole-3-carboxamide (3.40 mg, 9.27 $\mu$mol, 1.68% yield) as a light-yellow solid. $^1$H NMR (600 MHz, dmso) $\delta_H$ 1.41 (t, 3H), 2.14 (p, 2H), 2.94 – 3.00 (m, 5H), 3.04 (s, 3H), 4.12 (t, 2H), 4.39 (q, 2H), 6.52 (s, 1H), 6.90 (dd, 1H), 7.16 (d, 1H), 7.54 (d, 1H). LCMS(ESI): $[M+H]^+$ m/z: calcd 360.17; found 360.0.

#### Synthesis of 5-[3-[(3-ethoxy-5-methyl-1,2-benzoxazol-6-yl)oxy]propyl]-N,N-dimethyl-isoxazole-3-carboxamide (Compound 47)

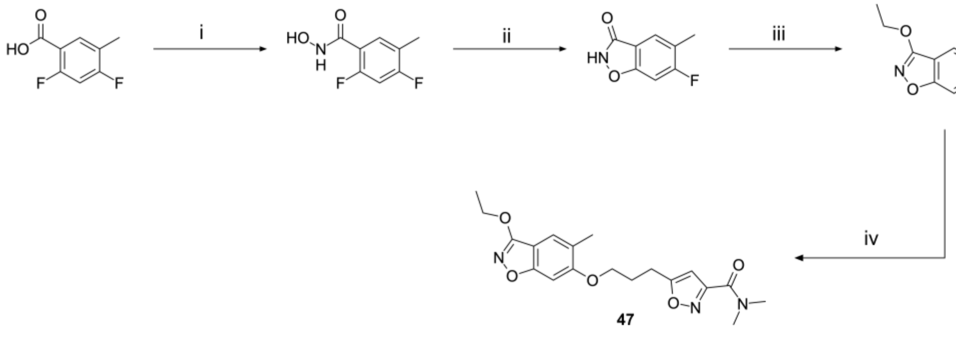


Scheme S16: Reagents and conditions: (i) hydroxylamine, oxalyl chloride, DCM, DMF, 0 °C to RT, 16h (ii) KOH, $^n$BuOH, 130 °C, 4h (iii) $Ag_2O$, iodoethane, $CHCl_3$, RT to reflux, 16h (iv) 5-(3-hydroxypropyl)-N,N-dimethyl-isoxazole-3-carboxamide, NaH, DMF, 100 °C, 16h

#### Step i: Data for 2,4-difluoro-5-methyl-benzenecarbohydroxamic acid

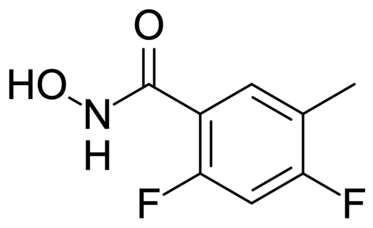


A catalytic amount of DMF and oxalyl chloride (3.48 g, 27.5 mmol, 2.39 mL) were added sequentially to a stirred solution of 2,4-difluoro-5-methyl-benzoic acid (3.15 g, 18.3 mmol) in DCM (50 mL) at 0 °C. The resulting mixture was allowed to warm up to room temperature and stirred overnight. The reaction mixture was concentrated under reduced pressure. The residue was redissolved in a minimum amount of EtOAc. This solution was added dropwise to a vigorously stirring mixture of hydroxylamine (2.54 g, 36.6 mmol, 1.52 mL, HCl), $K_2CO_3$ (10.1 g, 73.2 mmol), water (80 mL) and EtOAc (160 mL) at 0 °C. The aqueous layer was separated and extracted with EtOAc (80 mL). The combined organic layers were dried over anhydrous sodium sulfate and concentrated under reduced pressure to afford 2,4-difluoro-5-methyl-benzenecarbohydroxamic acid (3.00 g, 16.0 mmol, 87.6% yield) as a brown solid which was used in the next step without further purification. $^1$H NMR (500 MHz, dmso) $\delta_H$ 2.21 (s, 3H), 7.26 (t, 1H), 7.49 (t, 1H), 9.19 (s, 1H), 10.91 (s, 1H). LCMS(ESI): $[M+H]^+$ m/z: calcd 188.05; found 188.0.

#### Step ii: Data for 6-fluoro-5-methyl-1,2-benzoxazol-3-one

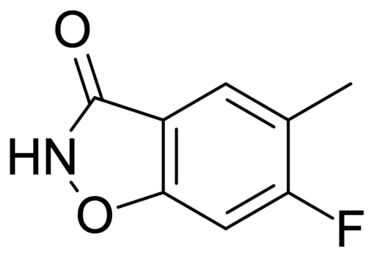


KOH (1.08 g, 19.2 mmol) was added to a stirred solution of 2,4-difluoro-5-methyl-benzenecarbohydroxamic acid (600 mg, 3.21 mmol) in $^n$BuOH (6.0 mL) at room temperature. The resulting mixture was stirred at 130 °C for 4 hr. The reaction mixture was cooled to room temperature and concentrated under reduced pressure. The residue was diluted with water (10 mL), acidified with hydrochloric acid and extracted with EtOAc (20 mL × 2). The combined organic layers were washed with brine (10 mL), dried over anhydrous sodium sulfate and concentrated under reduced pressure. The residue was subjected to column chromatography (ISCO®: CombiFlash; 40 g $SiO_2$, gradient $CHCl_3$/MTBE with MTBE from 0 95%, flow rate = 40 mL/min, Rf = 8-10 CV.) to afford 6-fluoro-5-methyl-1,2-benzoxazol-3-one (200 mg, 1.20 mmol, 37.3% yield) as a yellow solid. $^1$H NMR (500 MHz, dmso) $\delta_H$ 2.30 (s, 3H), 7.46 (d, 1H), 7.64 (d, 1H), 12.34 (br. s, 1H). LCMS(ESI): $[M+H]^+$ m/z: calcd 168.05; found 168.0.

#### Step iii: Data for 3-ethoxy-6-fluoro-5-methyl-1,2-benzoxazole

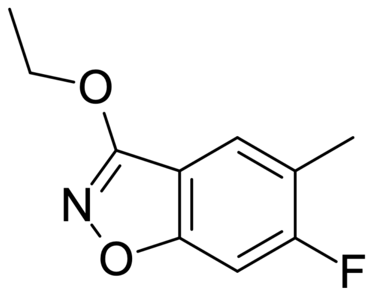


$Ag_2O$ (555 mg, 2.39 mmol) and iodoethane (187 mg, 1.20 mmol, 96.2 $\mu$L) were added sequentially to a stirred solution of 6-fluoro-5-methyl-1,2-benzoxazol-3-one (200 mg, 1.20 mmol) in $CHCl_3$ (50 mL) at room temperature. The resulting mixture was stirred under reflux overnight. The reaction mixture was cooled to room temperature and filtered through a thin pad of silica gel. The filtercake was washed several times with $CHCl_3$. The filtrate was concentrated under reduced pressure. The residue was subjected to reverse phase HPLC (0-1-5 min., 35-35-80% water – ACN, flow: 30 mL/min, column: Chromatorex 18 SMB100-5T 100×19 mm, 5 $\mu$m) to afford 3-ethoxy-6-fluoro-5-methyl-1,2-benzoxazole (65.0 mg, 333 $\mu$mol, 27.8% yield) as a yellow solid. LCMS(ESI): $[M+H]^+$ m/z: calcd 196.09; found 196.0.

#### Step iv: Data for 5-[3-[(3-ethoxy-5-methyl-1,2-benzoxazol-6-yl)oxy]propyl]-N,N-dimethyl-isoxazole-3-carboxamide (Compound 47)

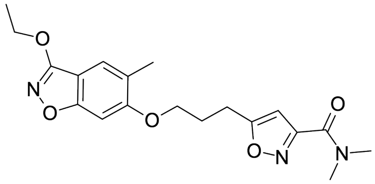


NaH (6.45 mg, 161 $\mu$mol, 60% dispersion in mineral oil) was added to a stirred solution of 5-(3-hydroxypropyl)-N,N-dimethyl-isoxazole-3-carboxamide (32.0 mg, 161 $\mu$mol)

in DMF (3.0 mL) at room temperature. The mixture was stirred for 15 min then 3-ethoxy-6-fluoro-5-methyl-1,2-benzoxazole (30.0 mg, 154 $\mu$mol) was added. The resulting mixture was stirred at 100 °C overnight. The reaction mixture was cooled to room temperature and subjected to preparative HPLC (0-1-7 min., 30-35-60% water+FA (0.1% vol.) - ACN+FA (0.1% vol.); flow: 30 mL/min, column: Chromatorex 18 SMB100-5T 100×19 mm, 5 $\mu$m) to afford 5-[3-[(3-ethoxy-5-methyl-1,2-benzoxazol-6-yl)oxy]propyl]-N,N-dimethyl-isoxazole-3-carboxamide (16.5 mg, 44.2 $\mu$mol, 28.8% yield) as a white solid. $^1$H NMR (500 MHz, dmso) $\delta_H$ 1.40 (t, 3H), 2.13 – 2.22 (m, 5H), 2.97 (s, 3H), 3.00 (t, 2H), 3.03 (s, 3H), 4.11 (t, 2H), 4.38 (q, 2H), 6.52 (s, 1H), 7.14 (s, 1H), 7.41 (s, 1H). LCMS(ESI): $[M+H]^+$ m/z: calcd 374.19; found 374.2.

#### Synthesis of 5-[3-[(3-ethoxy-7-methyl-1,2-benzoxazol-6-yl)oxy]propyl]-N,N-dimethyl-isoxazole-3-carboxamide (Compound 48)

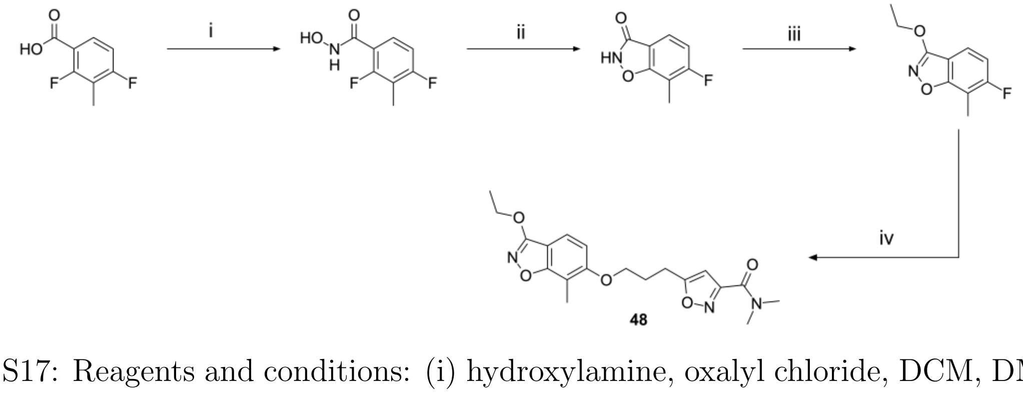


Scheme S17: Reagents and conditions: (i) hydroxylamine, oxalyl chloride, DCM, DMF, 0 °C to RT, 16h (ii) KOH, $^n$BuOH, 130 °C, 4h (iii) $Ag_2O$, iodoethane, $CHCl_3$, RT to reflux, 16h (iv) 5-(3-hydroxypropyl)-N,N-dimethyl-isoxazole-3-carboxamide, NaH, DMF, 100 °C, 16h

#### Step i: Data for 2,4-difluoro-3-methyl-benzenecarbohydroxamic acid

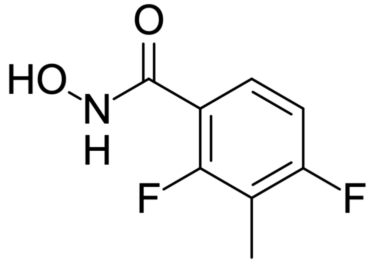

A catalytic amount of DMF and oxalyl chloride (553 mg, 4.36 mmol, 380 $\mu$L) were added sequentially to a stirred solution of 2,4-difluoro-3-methyl-benzoic acid (500 mg, 2.90 mmol) in DCM (10 mL) at 0 °C. The resulting mixture was allowed to warm up to room temperature and stirred overnight. The reaction mixture was concentrated under reduced pressure. The residue was redissolved in a minimum amount of EtOAc. This solution was added dropwise to a vigorously stirring mixture of hydroxylamine (404 mg, 5.81 mmol, HCl), $K_2CO_3$ (1.61 g, 11.6 mmol), water (15 mL) and EtOAc (30 mL) at 0 °C. The aqueous layer was separated and extracted with EtOAc (40 mL). The organic layer was dried over anhydrous sodium sulfate and concentrated under reduced pressure to afford 2,4-difluoro-3-methyl-benzenecarbohydroxamic acid (460 mg, 2.46 mmol, 84.6% yield) as a red solid which was used in the next step without further purification. $^1$H NMR (500 MHz, dmso) $\delta_H$ 2.16 (s, 3H), 7.12 (t, 1H), 7.40 (q, 1H), 9.20 (s, 1H), 10.95 (s, 1H). LCMS(ESI): $[M+H]^+$ m/z: calcd 188.05; found 188.0.

#### Step ii: Data for 6-fluoro-7-methyl-1,2-benzoxazol-3-one

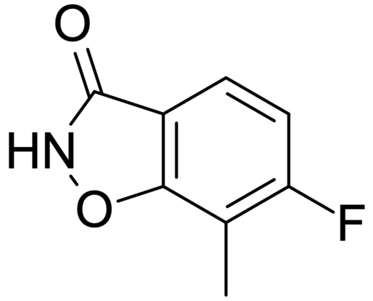

KOH (828 mg, 14.8 mmol) was added to a stirred solution of 2,4-difluoro-3-methyl-benzenecarbohydroxamic acid (460 mg, 2.46 mmol) in n-BuOH (10 mL) at room temperature. The resulting mixture was stirred at 100 °C for 4 hr. The reaction mixture was cooled to room temperature and concentrated under reduced pressure. The residue was diluted with water (10 mL), acidified with hydrochloric acid and extracted with EtOAc (20 mL × 2). The combined organic layers were washed with brine (10 mL), dried over anhydrous sodium sulfate and concentrated under reduced pressure to afford 6-fluoro-7-methyl-1,2-benzoxazol-3-one (400 mg, 1.68 mmol, 68.2% yield) as a brown solid which was used in the next step without further purification. LCMS(ESI): [M-H]- m/z: calcd 166.03; found 166.0.

#### Step iii: Data for 3-ethoxy-6-fluoro-7-methyl-1,2-benzoxazole

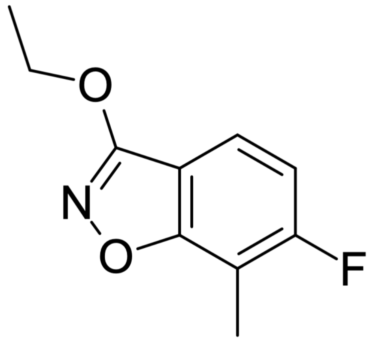


$Ag_2O$ (776 mg, 3.35 mmol) and iodoethane (26 mg, 1.68 mmol, 135 $\mu$L) were added to a stirred solution of 6-fluoro-7-methyl-1,2-benzoxazol-3-one (400 mg, 1.68 mmol) in $CHCl_3$ (25 mL) at room temperature. The resulting mixture was stirred under reflux overnight. The reaction mixture was cooled to room temperature and filtered through a thin pad of silica gel. The filtercake was washed several times with $CHCl_3$. The filtrate was concentrated under reduced pressure. The residue was subjected to reverse phase HPLC (0-1-5 min., 35-35-80% water – ACN, flow: 30 mL/min, column: Chromatorex 18 SMB100-5T 100×19 mm, 5 $\mu$m) to afford 3-ethoxy-6-fluoro-7-methyl-1,2-benzoxazole (77.0 mg, 394 $\mu$mol, 23.6% yield) as a yellow solid. $^1$H NMR (500 MHz, dmso) $\delta_H$ 1.51 (t, 3H), 2.39 (s, 3H), 4.47 (q, 2H), 6.98 (t, 1H), 7.39 (dd, 1H). LCMS(ESI): $[M+H]^+$ m/z: calcd 196.09; found 196.0.

#### Step iv: Data for 5-[3-[(3-ethoxy-7-methyl-1,2-benzoxazol-6-yl)oxy]propyl]-N,N-dimethyl-isoxazole-3-carboxamide (Compound 48)

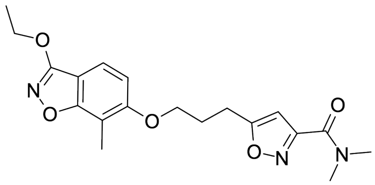


NaH (6.45 mg, 161 $\mu$mol, 60% dispersion in mineral oil) was added to a stirred solution of 5-(3-hydroxypropyl)-N,N-dimethyl-isoxazole-3-carboxamide (32.0 mg, 161 $\mu$mol) in DMF (3.0 mL) at room temperature. The mixture was stirred for 15 min then 3-ethoxy-6-fluoro-7-methyl-1,2-

benzoxazole (30.0 mg, 154 $\mu$mol) was added. The resulting mixture was stirred at 100 °C overnight. The reaction mixture was cooled to room temperature and subjected to preparative HPLC (0-6 min., 30-60% water – ACN, flow: 30 mL/min, column: Chromatorex 18 SMB100-5T 100×19 mm, 5 $\mu$m) to afford 5-[3-[(3-ethoxy-7-methyl-1,2-benzoxazol-6-yl)oxy]propyl]-N,N-dimethyl-isoxazole-3-carboxamide (5.40 mg, 14.5 $\mu$mol, 9.41% yield) as a beige solid. $^1$H NMR (500 MHz, $CD_3OD$) $\delta_H$ 1.48 (t, 3H), 2.22 – 2.31 (m, 5H), 3.05 – 3.12 (m, 5H), 3.17 (s, 3H), 4.17 (t, 2H), 4.42 (q, 2H), 6.41 (s, 1H), 6.99 (d, 1H), 7.40 (d, 1H). LCMS(ESI): $[M+H]^+$ m/z: calcd 374.19; found 374.2.

#### Synthesis of 5-[3-[[3-(cyclopropylmethoxy)-1,2-benzoxazol-6-yl]oxy]propyl]-N,N-dimethyl-isoxazole-3-carboxamide (Compound 49)

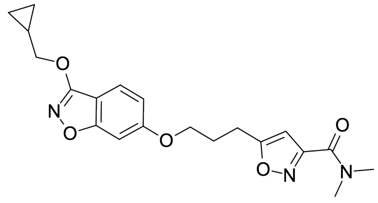

**49** was synthesised by a route similar to Compound **46**: 5-(3-Hydroxypropyl)-N,N-dimethyl-isoxazole-3-carboxamide (332 mg, 1.67 mmol) was added to the previously obtained solution of 6-fluoro-3-isobutoxy-1,2-3-(cyclopropylmethoxy)-6-fluoro-1,2-benzoxazole (100 mg, 0.48 mmol) in DMF (2 mL). The obtained solution was cooled to 0 °C. To the resulting mixture NaH (19 mg, 0.48 mmol, 60% dispersion in mineral oil) was added. The reaction mixture was stirred at ambient temperature for 16 hrs. The resulting mixture was diluted with water (5 mL) and extracted with ethyl acetate (10 mL). The organic layer was washed with water (3 × 1 mL), dried over anhydrous sodium sulphate and concentrated. The residue was subjected to HPLC (0-1.3-5 min. 35-35-75%, water – ACN +0.1% vol. of 25% aq. $NH_3$, flow: 30 mL/min; column: XBridge C18 100×19mm,) to afford 15.1 mg of a mixture of two regioisomers. The resulting mixture was submitted to chiral HPLC (Column: CHIRALPAK IC 250×21 mm, 5 $\mu$m, mobile phase: Hexane-IPA-MeOH, 50:25:25, flow: 12 mL/min) to afford 5-[3-[[3-(cyclopropylmethoxy)-

1,2-benzoxazol-6-yl]oxy]propyl]-N,N-dimethyl-isoxazole-3-carboxamide (9.5 mg, 23.4 $\mu$mol, 4.85% yield) as a white solid. $^1$H NMR (500 MHz, $CD_3OD$) $\delta_H$ 0.40 – 0.44 (m, 2H), 0.65 – 0.68 (m, 2H), 1.35 – 1.43 (m, 1H), 2.24 (p, 2H), 3.06 (t, 2H), 3.10 (s, 3H), 3.18 (s, 3H), 4.13 (t, 2H), 4.19 (d, 2H), 6.41 (s, 1H), 6.91 (dd, 1H), 6.99 (d, 1H), 7.52 (d, 1H). LCMS(ESI): $[M+H]^+$ m/z: calcd 386.18; found 386.0.

#### Synthesis of 5-(3-((3-isobutoxybenzo[d]isoxazol-6-yl)oxy)propyl)-N,N-dimethylisoxazole-3-carboxamide (50) & 5-(3-((6-isobutoxybenzo[d]isoxazol-3-yl)oxy)propyl)-N,N-dimethylisoxazole-3-carboxamide (51)

#### (Compound 50)

#### (Compound 51)

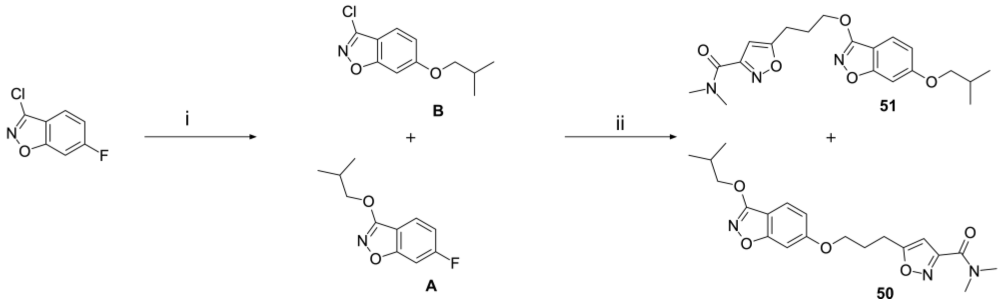


Scheme S18: Reagents and conditions: (i) NaH, EtOH, DMF, 0 °C to RT, 16h (ii) 5-(3-Hydroxypropyl)-N,N-dimethyl-isoxazole-3-carboxamide, NaH, DMF, 0 °C to RT, 16h

Compound **50** was synthesised by a route similar to Compound **46**. In this case, the isomeric product: Compound **51** was also isolated.

#### Step i: Data for 6-fluoro-3-isobutoxybenzo[d]isoxazole A and 3-chloro-6-isobutoxybenzo[d]isoxazole B

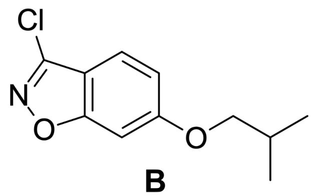

**B**

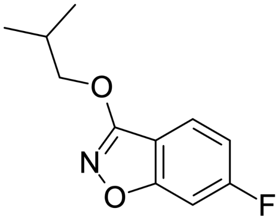

**A**

3-Chloro-6-fluoro-1,2-benzoxazole (400 mg, 2.33 mmol) and 2-methylpropan-1-ol (190 mg, 2.56 mmol) was dissolved in DMF (8 mL) at 0 ºC. NaH (100 mg, 2.5 mmol, 60% dispersion in mineral oil) was added portionwise at 0°C. The reaction mixture was stirred for 16 hrs. at ambient temperature. The obtained solution was used in the next step without work up. 6-fluoro-3-isobutoxy-1,2-benzoxazole (**A**): LCMS(ESI): $[M+H]^+$ m/z: calcd 210.10; found 210.0. 3-chloro-6-isobutoxybenzo[d]isoxazole (**B**): LCMS(ESI): $[M+H]^+$ m/z: calcd 226.07; found 226.0.

##### Step ii: Data for 5-(3-((3-isobutoxybenzo[d]isoxazol-6-yl)oxy)propyl)-N,N-dimethylisoxazole-3-carboxamide (Compound 50) and 3-chloro-6-isobutoxybenzo[d]isoxazole (Compound 51)

5-(3-Hydroxypropyl)-N,N-dimethyl-isoxazole-3-carboxamide (332 mg, 1.67 mmol) was added to the previously obtained solution of 6-fluoro-3-isobutoxy-1,2-benzoxazole (0.35 g, 1.67 mmol) in DMF (8 mL). To the resulting mixture NaH (67 mg, 1.67 mmol, 60% dispersion in mineral oil) was added at 0 ºC. The reaction mixture was stirred at ambient temperature for 16 hrs. The resulting mixture was diluted with water (10 mL) and extracted with ethyl acetate (20 mL). The organic layer was washed with water (3 × 1 mL), dried over anhydrous sodium sulphate and concentrated. The residue was subjected to HPLC (0-1.3-6 min. 35-35-75%, water - ACN, flow: 30 mL/min; column: XBridge C18 100×19mm,) to afford a mixture of two regioisomers. The resulting mixture was submitted to chiral HPLC (Column: CHIRALPAK IC 250×21 mm, 5 $\mu$m, mobile phase: IPA-MeOH, 50:50, flow: 10 mL/min) to afford 5-[3-[(3-isobutoxy-1,2-benzoxazol-6-yl)oxy]propyl]-N,N-dimethyl-isoxazole-3-carboxamide (**50**, 8.5 mg, 21.9 $\mu$mol, 1.31% yield) and 5-(3-((6-isobutoxybenzo[d]isoxazol-3-yl)oxy)propyl)-N,N-dimethylisoxazole-3-carboxamide (**51**, 8.4 mg, 21.7 $\mu$mol, 1.30% yield).

Compound **50**

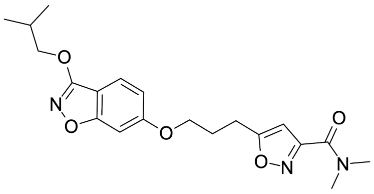


Analytical data: $^1$H NMR (500 MHz, $CD_3OD$) $\delta_H$ 1.07 (d, 6H), 2.15 – 2.22 (m, 1H), 2.22 – 2.28 (m, 2H), 3.07 (t, 2H), 3.10 (s, 3H), 3.19 (s, 3H), 4.11 – 4.17 (m, 4H), 6.41 (s, 1H), 6.92 (d, 1H), 7.00 (s, 1H), 7.51 (d, 1H). LCMS(ESI): $[M+H]^+$ m/z: calcd 388.21; found 388.2. Analytical chiral HPLC (column: Chiralpak IC (250 $\times$ 4.6 mm, 5 $\mu$m; mobile phase: IPA-MeOH, 50:50; flow 0.6 mL/min.): Rt 18.37 min.

Compound **51**

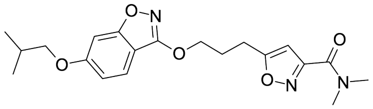


Analytical data: $^1$H NMR (500 MHz, $CD_3OD$) $\delta_H$ 1.06 (d, 6H), 2.07 – 2.14 (m, 1H), 2.32 (p, 2H), 3.08 (t, 2H), 3.10 (s, 3H), 3.18 (s, 3H), 3.82 (d, 2H), 4.46 (t, 2H), 6.44 (s, 1H), 6.91 (dd, 1H), 6.97 (d, 1H), 7.49 (d, 1H). LCMS(ESI): $[M+H]^+$ m/z: calcd 388.21; found 388.2. Analytical chiral HPLC (column: Chiralpak IC (250 $\times$ 4.6 mm, 5 $\mu$m; mobile phase: IPA-MeOH, 50:50; flow 0.6 mL/min.): Rt 21.71 min.

#### General Procedure A: Parallel Synthesis of Phenol Ethers by Mitsunobu Coupling

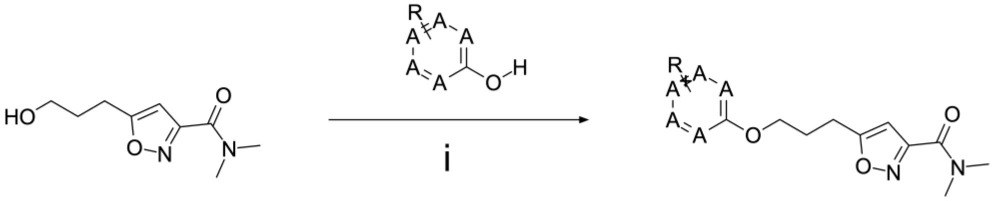


Scheme S19: Reagents and conditions: (i) hydroxy aryl compound, ADDP, $PBu_3$, dioxane, RT, 16h

#### Data for 5-(3-(3-Fluorophenoxy)propyl)-N,N-dimethylisoxazole-3-carboxamide (Compound 52)

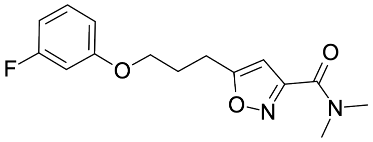


Following General Procedure A using 3-fluorophenol afforded 12.9 mg of 5-(3-(3-fluorophenoxy)propyl)-N,N-dimethylisoxazole-3-carboxamide as a beige solid (44.1 $\mu$mol, 25.8% yield). HPLC method: 1.3-7.3 min., 30-50% water – ACN, +0.1% vol. of 25% aq. $NH_3$, flow: 30 mL/min + 4 mL/min (organic phase), column: XBridge BEH C18 5$\mu$m 130A, 100 ×19mm, 5$\mu$m. $^1$H NMR (500 MHz, dmso) $\delta_H$ 2.09 (p, 2H), 2.95 (t, 2H), 2.98 (s, 3H), 3.03 (s, 3H), 4.03 (t, 2H), 6.51 (s, 1H), 6.70 – 6.82 (m, 3H), 7.29 (q, 1H). LCMS(ESI): $[M+H]^+$ m/z: calcd 293.15; found 293.2.

#### Data for 5-(3-((1-Fluoro-5,6,7,8-tetrahydronaphthalen-2-yl)oxy)propyl)-N,N-dimethylisoxazole-3-carboxamide (Compound 53)

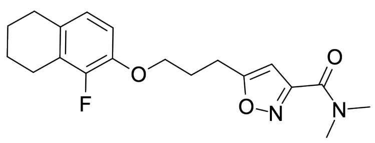

Following General Procedure A using 1-fluoro-5,6,7,8-tetrahydronaphthalen-2-ol afforded 12.0 mg of 5-(3-((1-fluoro-5,6,7,8-tetrahydronaphthalen-2-yl)oxy)propyl) -N,N-dimethylisoxazole-3-carboxamide as a white solid (34.6 $\mu$mol, 24.0% yield). HPLC method: 1.3-7.3 min., 40-65% water – ACN, +0.1% vol. of 25% aq. $NH_3$, flow: 30 mL/min + 4 mL/min (organic phase), column: XBridge BEH C18 5$\mu$m 130A, 100 ×19mm, 5$\mu$m. $^1$H NMR (500 MHz, dmso) $\delta_H$ 1.62 – 1.72 (m, 4H), 2.08 (p, 2H), 2.57 – 2.67 (m, 4H), 2.94 (t, 2H), 2.97 (s, 3H), 3.03 (s, 3H), 4.02 (t, 2H), 6.50 (s, 1H), 6.79 (d, 1H), 6.88 (t, 1H). LCMS(ESI): $[M+H]^+$ m/z: calcd 347.21; found 347.2.

#### Data for 5-(3-(4-Fluorophenoxy)propyl)-N,N-dimethylisoxazole-3-carboxamide (Compound 54)

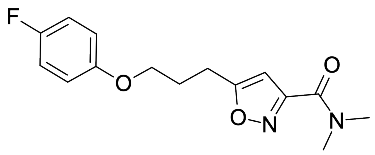

Following General Porcedure A using 4-fluorophenol afforded 15.1 mg of 5-(3-(4-fluorophenoxy)propyl)-N,N-dimethylisoxazole-3-carboxamide as a beige solid (52.0 $\mu$mol, 30.2% yield). HPLC method: 1.3-7.3 min., 30-55% water – ACN, +0.1% vol. of 25% aq. $NH_3$, flow: 30 mL/min + 4 mL/min (organic phase), column: XBridge BEH C18 5$\mu$m 130A, 100 ×19mm, 5$\mu$m. $^1$H NMR (500 MHz, dmso) $\delta_H$ 2.08 (p, 2H), 2.95 (t, 2H), 2.98 (s, 3H), 3.03 (s, 3H), 3.98 (t, 2H), 6.50 (s, 1H), 6.88 – 6.96 (m, 2H), 7.09 (t, 2H). LCMS(ESI): $[M+H]^+$ m/z: calcd 293.15; found 293.2.

#### Data for 5-(3-(4-(prop-2-yn-1-yl)phenoxy)propyl)-N,N-Dimethylisoxazole-3-carboxamide (Compound 55)

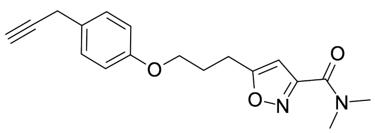

Following General Procedure A using 4-(prop-2-yn-1-yl)phenol afforded 19.5 mg of 5-(3-(4-(prop-2-yn-1-yl)phenoxy)propyl)-N,N-Dimethylisoxazole-3-carboxamide as a yellow solid (62.4 $\mu$mol, 39.0% yield). HPLC method: 1.3-7.3 min., 30-50% water – ACN, +0.1% vol. of 25% aq. $NH_3$, flow: 30 mL/min + 4 mL/min (organic phase), column: XBridge BEH C18 5$\mu$m 130A, 100 ×19mm, 5$\mu$m. $^1$H NMR (500 MHz, dmso) $\delta_H$ 2.09 (p, 2H), 2.95 (t, 2H), 2.97 (s, 3H), 3.00 (s, 1H), 3.03 (s, 3H), 3.52 (s, 2H), 3.99 (t, 2H), 6.50 (s, 1H), 6.87 (d, 2H), 7.20 (d, 2H). LCMS(ESI): $[M+H]^+$ m/z: calcd 313.18; found 313.2.

#### Data for 5-(3-(4-(But-3-yn-1-yl)phenoxy)propyl)-N,N-dimethylisoxazole-3-carboxamide (Compound 56)

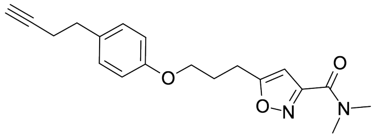

Following General Procedure A using 4-(But-3-yn-1-yl)phenol afforded 14.1 mg of 5-(3-(4-(but-3-yn-1-yl)phenoxy)propyl)-N,N-dimethylisoxazole-3-carboxamide as a yellow gum (43.2 $\mu$mol, 28.2% yield). HPLC method: 1.3-7.3 min., 30-50% water – ACN, +0.1% vol. of 25% aq. $NH_3$, flow: 30 mL/min + 4 mL/min (organic phase), column: XBridge BEH C18 5$\mu$m 130A, 100 ×19mm, 5$\mu$m. $^1$H NMR (500 MHz, dmso) $\delta_H$ 2.08 (p, 2H), 2.35 – 2.40 (m, 2H), 2.66 (t, 2H), 2.74 (t, 1H), 2.94 (t, 2H), 2.97 (s, 3H), 3.03 (s, 3H), 3.98 (t, 2H), 6.50 (s, 1H), 6.82 (d, 2H), 7.13 (d, 2H). LCMS(ESI): $[M+H]^+$ m/z: calcd 327.30; found 327.2.

#### Synthesis of (E)-N-ethoxy-1-[4-[2-[1-(6-methylpyridazin-3-yl)-4-piperidyl] ethoxy]phenyl]methanimine (Compound S-57 (BTA-188))

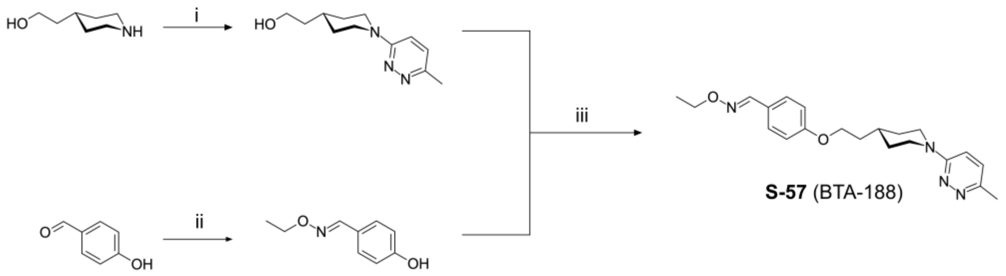

Scheme S20: Reagents and conditions: (i) 3-chloro-6-methyl-pyridazine, $Na_2CO_3$, DMAA, 150 °C, 16h (ii) O-ethylhydroxylamine, $NaHCO_3$, dioxane, reflux, 16h (iii) $PPh_3$, DIAD, THF, 16h

#### Step i: Data for 2-[1-(6-methylpyridazin-3-yl)-4-piperidyl]ethanol

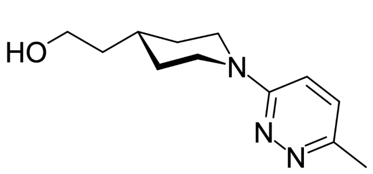

$Na_2CO_3$ (1.28 g, 12.1 mmol) and 2-(4-piperidyl)ethanol (1.00 g, 6.04 mmol, HCl) were added to a stirred solution of 3-chloro-6-methyl-pyridazine (776 mg, 6.04 mmol) in DMAA (5.0 mL) at room temperature. The resulting mixture was stirred at 150 °C overnight. The resulting mixture was cooled to room temperature and filtered through a thin pad of silica. The filtrate was subjected to HPLC (0-1-5 min., 10-10-50% water – MeOH, +0.1% vol. of 25% aq. $NH_3$, flow: 30 mL/min, column: XBridge C18 100×19 mm, 5 $\mu$m) to afford 2-[1-(6-methylpyridazin-3-yl)-4-piperidyl]ethanol (570 mg, 2.58 mmol, 42.7% yield) as a brown solid. LCMS(ESI): $[M+H]^+$ m/z: calcd 222.19; found 222.2.

#### Step ii: Data for 4-[(E)-ethoxyiminomethyl]phenol

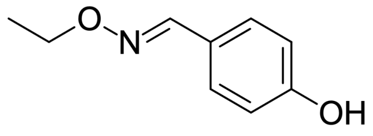

O-ethylhydroxylamine (3.99 g, 40.9 mmol, HCl) and $NaHCO_3$ (3.44 g, 40.9 mmol) were added to a stirred solution of 4-hydroxybenzaldehyde (1.00 g, 8.19 mmol, 886 $\mu$L) in

dioxane (20 mL) at room temperature. The resulting mixture was stirred under reflux overnight. The reaction mixture was cooled to room temperature and concentrated under reduced pressure. The residue was diluted in EtOAc (50 mL) and washed with water (10 mL). The organic layer was separated, dried over anhydrous sodium sulfate and concentrated under reduced pressure to afford 4-[(E)-ethoxyiminomethyl]phenol (1.30 g, 7.87 mmol, 96.1% yield) as a beige solid which was used in the next step without further purification. LCMS(ESI): $[M+H]^+$ m/z: calcd 166.09; found 166.2.

#### Step iii: Data for (E)-N-ethoxy-1-[4-[2-[1-(6-methylpyridazin-3-yl)-4-piperidyl]ethoxy]phenyl]methanimine (Compound S-57 (BTA-188))

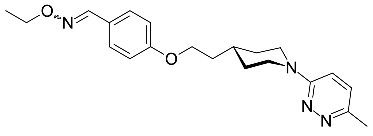


2-[1-(6-Methylpyridazin-3-yl)-4-piperidyl]ethanol (100 mg, 452 $\mu$mol), 4-[(E)-ethoxyiminomethyl]phenol (82.1 mg, 497 $\mu$mol) and $PPh_3$ (130 mg, 497 $\mu$mol) were dissolved in THF (5.0 mL) at room temperature. To the obtained solution DIAD (101 mg, 497 $\mu$mol, 98 $\mu$L) was added dropwise at room temperature. The resulting mixture was stirred at room temperature overnight. The reaction mixture was concentrated under reduced pressure. The residue was subjected to HPLC (0-1-5 min., 45-45-65% water – ACN, +0.1% vol. of 25% aq. $NH_3$, flow: 30 mL/min, column: XBridge C18 100×19 mm, 5 $\mu$m) to afford (E)-N-ethoxy-1-[4-[2-[1-(6-methylpyridazin-3-yl)-4-piperidyl]ethoxy]phenyl]methanimine (32.0 mg, 86.9 $\mu$mol, 19.2% yield) as a beige solid. $^1$H NMR (500 MHz, dmso) $\delta_H$ 1.16 – 1.26 (m, 5H), 1.66 – 1.72 (m, 2H), 1.73 – 1.82 (m, 3H), 2.41 (s, 3H), 2.84 (t, 2H), 4.07 (t, 2H), 4.11 (q, 2H), 4.29 (d, 2H), 6.98 (d, 2H), 7.21 (q, 2H), 7.53 (d, 2H), 8.14 (s, 1H). LCMS(ESI): $[M+H]^+$ m/z: calcd 369.27; found 369.2. The data were in agreement with those published.[S1]

#### Synthesis of 1-(2-amino-4-pyridyl)-3-[5-[4-(5-methyl-1,2,4-oxadiazol-3-yl)phenoxy]pentyl]imidazolidin-2-one (Compound S-59 (NLD-22))

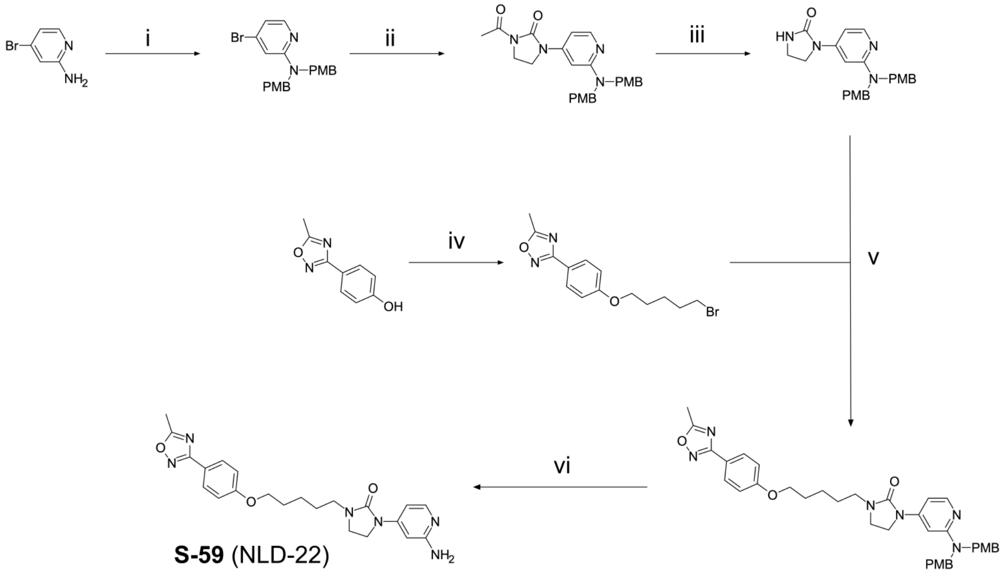

Scheme S21: Reagents and conditions: (i) 1-(chloromethyl)-4-methoxy-benzene, NaH, DMF, 0 °C to RT, 16h (ii) 1-acetylimidazolidin-2-one, $Cs_2CO_3$, $Pd(dba)_2$, Xantphos, toluene, 100 °C, 16h (iii) $K_2CO_3$, MeOH, RT, 16h (iv) 5-bromopentan-1-ol, $PPh_3$, DIAD, THF, 0 °C to RT, 16h (v) NaH, DMF, 0 °C to RT, 16h (vi) TFA, DCM, 0 °C to RT, 16h

#### Step i: Data for 4-bromo-N,N-bis[(4-methoxyphenyl)methyl]pyridin-2-amine

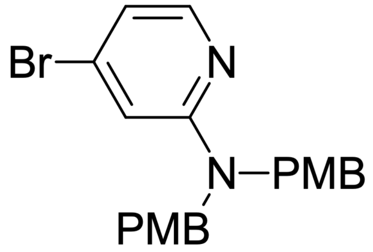

4-Bromopyridin-2-amine (1.00 g, 5.78 mmol) was added to a stirred solution of NaH (925 mg, 23.1 mmol, 60% dispersion in mineral oil) in DMF (20 mL) at 0 °C. The resulting mixture was stirred at ambient temperature for 30 min. The mixture was cooled to 0 °C then 1-(chloromethyl)-4-methoxy-benzene (2.26 g, 14.5 mmol) was added. The resulting mixture was stirred at room temperature for 16 hr. The reaction mixture was quenched by addition of water (25 mL). The resulting mixture was extracted with EtOAC (3×15 mL). The combined organic layers were washed with water (3×15 mL) and brine (15 mL), dried over anhydrous sodium sulfate and concentrated under reduced pressure to afford 4-bromo-N,N-bis[(4-methoxyphenyl)methyl]pyridin-2-amine (1.70 g, 4.11 mmol, 71.2% yield) as a yellow solid which was used in the next step without further purification. LCMS(ESI): $[M+H]^+$ m/z: calcd 413.09; found 413.0.

#### Step ii: Data for 1-acetyl-3-[2-[bis[(4-methoxyphenyl)methyl]amino]-4-pyridyl]imidazolidin-2-one

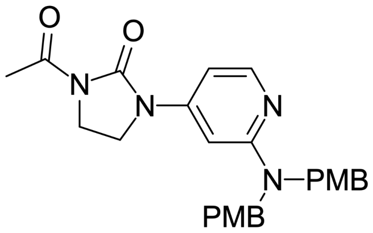


$Cs_2CO_3$ (591 mg, 1.81 mmol), 4-bromo-N,N-bis [(4-methoxyphenyl)methyl]pyridin-2-amine (500 mg, 1.21 mmol) and 1-acetylimidazolidin-2-one (465 mg, 3.63 mmol) were mixed in toluene (20 mL). The resulting suspension was degassed and backfilled with argon. To the resulting mixture $Pd(dba)_2$ (69.6 mg, 121 $\mu$mol) and Xantphos (70.00 mg, 121 $\mu$mol) were added sequentially. The reaction mixture was stirred at 100 °C for 16 hr. The reaction mixture was cooled to room temperature and filtered. The filtrate was concentrated under reduced pressure. The residue was subjected to column chromatography ($SiO_2$, eluent DCM - MeOH in ratio 95:5) to afford 1-acetyl-3-[2-[bis[(4-methoxyphenyl)methyl]amino]-4-pyridyl]imidazolidin-2-one (200 mg, 434 $\mu$mol, 35.9% yield) as a yellow solid. LCMS(ESI): $[M+H]^+$ m/z: calcd 461.22; found 461.2.

#### Step iii: Data for 1-[2-[bis[(4-methoxyphenyl)methyl]amino]-4-pyridyl] imidazolidin-2-one

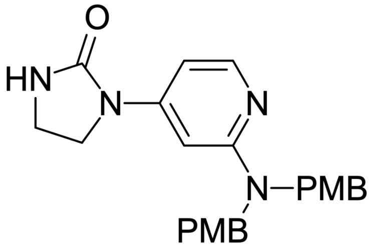


$K_2CO_3$ (180 mg, 1.30 mmol) was added to a stirred solution of 1-acetyl-3-[2-[bis[(4-methoxyphenyl)methyl]amino]-4-pyridyl]imidazolidin-2-one (200 mg, 434 $\mu$mol) in MeOH (15 mL) at room temperature. The resulting mixture was stirred at room temperature for 16 hr. The reaction mixture was diluted in water (30 mL) and filtered. The precipitate was washed with water and dried to afford 1-[2-[bis[(4-methoxyphenyl)methyl]amino]-4-pyridyl]imidazolidin-2-one (180 mg, 430 $\mu$mol, 99.0% yield) as a white solid which was used in the next step without further purification. LCMS(ESI): $[M+H]^+$ m/z: calcd 419.21; found 419.2

#### Step iv: Data for 3-[4-(5-bromopentoxy)phenyl]-5-methyl-1,2,4-oxadiazole

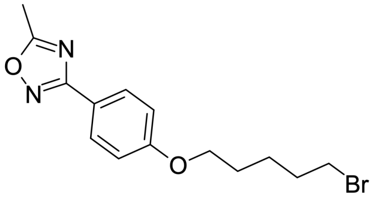


4-(5-Methyl-1,2,4-oxadiazol-3-yl)phenol (0.5 g, 2.84 mmol), 5-bromopentan-1-ol (545 mg, 3.26 mmol) and $PPh_3$ (856 mg, 3.26 mmol) were mixed in THF (20 mL) at room temperature. The mixture was cooled to 0 °C. DIAD (660 mg, 3.26 mmol, 643 $\mu$L) was added to the obtained mixture. The resulting reaction mixture was stirred under ambient conditions for 16 hr. The reaction mixture was concentrated under reduced pressure. The obtained residue was subjected to flash column chromatography ($SiO_2$, eluent Hex - EOAc in ratio 8:2) to afford 3-[4-(5-bromopentoxy)phenyl]-5-methyl-1,2,4-oxadiazole (0.60 g, 1.85 mmol, 65.0% yield) as a yellow gum. LCMS(ESI): $[M+H]^+$ m/z: calcd 325.06; found 325.0.

#### Step v: Data for 1-[2-[bis[(4-methoxyphenyl)methyl]amino]-4-pyridyl]-3-[5-[4-(5-methyl-1,2,4-oxadiazol-3-yl)phenoxy]pentyl]imidazolidin-2-one

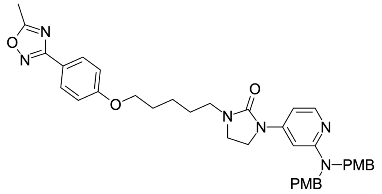

1-[2-[Bis[(4-methoxyphenyl)methyl]amino]-4-pyridyl]imidazolidin-2-one (90.0 mg, 215 $\mu$mol) was added to a stirred solution of NaH (12.9 mg, 323 $\mu$mol, 60% dispersion in oil) in DMF (10 mL) at 0 °C. The resulting mixture was stirred at ambient temperature for 30 min. The mixture was cooled to 0 °C then 3-[4-(5-bromopentoxy)phenyl]-5-methyl-1,2,4-oxadiazole (70.0 mg, 215 $\mu$mol) was added to the mixture. The reaction mixture was stirred at room temperature for 16 hr. The reaction mixture was diluted with water (20 mL) and extracted with EtOAc (2×15 mL). The combined organic layers were washed with water (2×10 mL) and brine (10 mL), dried over anhydrous sodium sulfate and concentrated under reduced pressure to afford 1-[2-[bis[(4-methoxyphenyl)methyl]amino]-4-pyridyl]-3-[5-[4-(5-methyl-1,2,4-oxadiazol-3-yl)phenoxy]pentyl]imidazolidin-2-one (100 mg, 151 $\mu$mol, 70.2% yield) as a yellow gum which was used in the next step without further purification. LCMS(ESI): $[M+H]^+$ m/z: calcd 663.33; found 663.2.

#### Step vi: Data for -(2-amino-4-pyridyl)-3-[5-[4-(5-methyl-1,2,4-oxadiazol-3-yl)phenoxy]pentyl]imidazolidin-2-one (Compound S-59 (NLD-22))

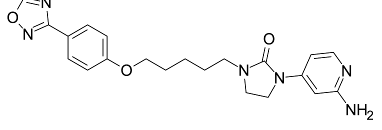

1-[2-[Bis[(4-methoxyphenyl)methyl]amino]-4-pyridyl]-3-[5-[4-(5-methyl-1,2,4-oxadiazol-3-yl)phenoxy]pentyl]imidazolidin-2-one (200 mg, 302 $\mu$mol) was dissolved in DCM (10 mL) and the resulting solution was cooled to 0 °C. TFA (344 mg, 3.02 mmol, 231 $\mu$L) was added to the stirred solution. The resulting mixture was stirred at room temperature for 16 hr. The reaction mixture was concentrated under reduced pressure. The residue was subjected to HPLC (0-1-5 min., 40-40-80% water – MeOH, +0.1% vol. of 25% aq. $NH_3$, flow: 30 mL/min, column: XBridge C18 100×19 mm, 5 $\mu$m) to afford 1-(2-amino-4-pyridyl)-3-[5-[4-(5-methyl-1,2,4-oxadiazol-3-yl)phenoxy]pentyl]imidazolidin-2-one (26.0 mg, 61.5 $\mu$mol, 20.4% yield) as a white solid. $^1$H NMR (500 MHz, dmso) $\delta_H$ 1.39 – 1.46 (m, 2H), 1.53 – 1.61 (m, 2H), 1.74 – 1.81 (m, 2H), 2.63 (s, 3H), 3.21 (t, 2H), 3.42 – 3.48 (m, 2H), 3.68 – 3.73 (m, 2H), 4.05 (t, 2H), 5.71 (s, 2H), 6.55 (d, 1H), 6.81 (dd, 1H), 7.09 (d, 2H), 7.24 (d, 1H), 7.91 (d, 2H). LCMS(ESI): $[M+H]^+$ m/z: calcd 423.24; found 423.2. The data were in agreement with those published.[S2]

#### Synthesis of 6-phenyl-N3-[4-(trifluoromethyl)phenyl]-1H-pyrazolo[3,4-d]pyrimidine-3,4-diamine (Compound S-60 (OBR-5-340))

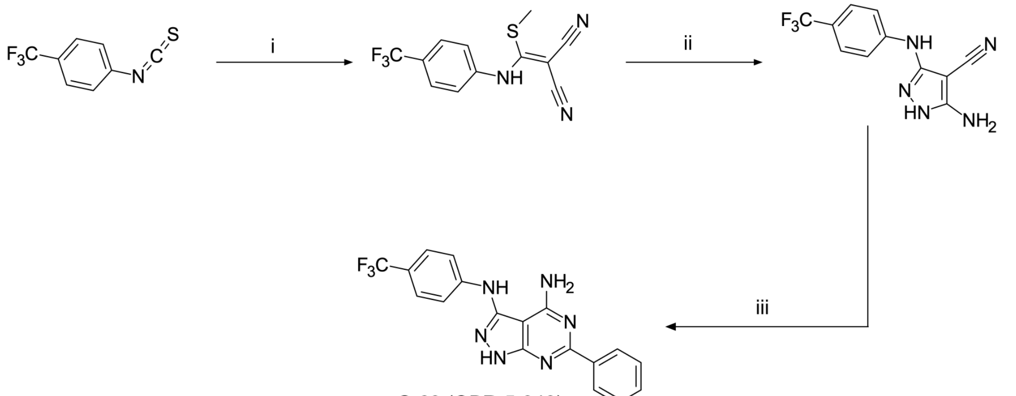


**S-60** (OBR-5-340)

Scheme S22: Reagents and conditions: (i) methyl iodide, malonitrile, NaH, DMF, RT, 16h (ii) hydrazine hydrate, EtOH, RT, reflux, 3h (iii) benzamidine, NaOAc, neat, 180 °C, 30 mins

#### Step i: Data for 2-[methylsulfanyl-[4-(trifluoromethyl)anilino]methylene] propanedinitrile

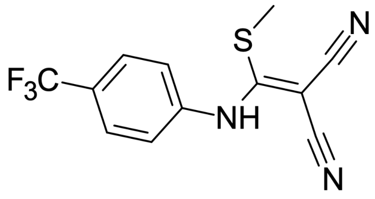

NaH (594 mg, 14.8 mmol, 60% dispersion in mineral oil) was added portionwise to a stirred solution of propanedinitrile (943 mg, 14.3 mmol) in DMF (50 mL) at room temperature. The resulting mixture was stirred at ambient temperature for 30 min. A solution of 1-isothiocyanato-4-(trifluoromethyl)benzene (2.90 g, 14.3 mmol) in DMF (5.0 mL) was added dropwise to the mixture. The resulting mixture was stirred at ambient temperature for 1 hr. Iodomethane (2.03 g, 14.3 mmol, 889 $\mu$L) was added dropwise at room temperature to the mixture. The resulting mixture was stirred at room temperature overnight. The reaction mixture was poured into water (400 mL), the obtained precipitate was filtered, washed with water (2×50 mL) and air dried to afford 2-[methylsulfanyl-[4-(trifluoromethyl)anilino]methylene]propanedinitrile (3.60 g, 10.9 mmol, 76.6% yield) as a yellow solid which was used in the next step without further purification. LCMS(ESI): $[M+H]^+$ m/z: calcd 284.05; found 284.0.

#### Step ii: Data for 6-phenyl-N3-[4-(trifluoromethyl)phenyl]-1H-pyrazolo[3,4-d]pyrimidine-3,4-diamine

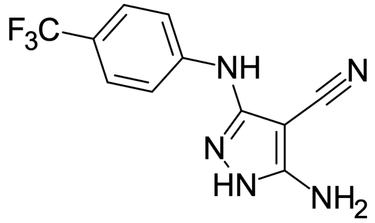

Hydrazine hydrate (3.50 g, 69.9 mmol) was added to a stirred solution of 2-[methylsulfanyl-[4-(trifluoromethyl) anilino]methylene]propanedinitrile (3.60 g, 12.7 mmol) in EtOH (50 mL) at room temperature. The resulting mixture was stirred under reflux for 3 hr. The reaction mixture was cooled to room temperature and diluted with water (50 mL). The obtained precipitate was filtered, washed with water (2×10 mL) and air dried to afford 3-amino-5-[4-(trifluoromethyl)anilino]-1H-pyrazole-4-carbonitrile (2.40 g, 8.98 mmol,

70.7% yield) as a beige solid which was used in the next step without further purification. LCMS(ESI): $[M+H]^+$ m/z: calcd 268.08; found 268.0.

#### Step iii: Data for 3-amino-5-[4-(trifluoromethyl)anilino]-1H-pyrazole-4-carbonitrile (Compound S-60 (OBR-5-340))

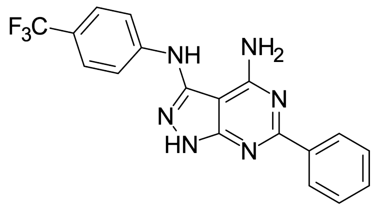


A mixture of 5-amino-3-[4-(trifluoromethyl)anilino]-1H-pyrazole-4-carbonitrile (800 mg, 2.99 mmol), benzamidine (1.17 g, 7.48 mmol, HCl) and NaOAc (614 mg, 7.48 mmol) was heated in neat at 180 °C for 30 min. The reaction mixture was cooled to room temperature and subjected to HPLC (0-1-5 min., 65-65-90% water – MeOH, +0.1% vol. of 25% aq. $NH_3$, flow: 30 mL/min, column: XBridge C18 100×19 mm, 5 $\mu$m) to afford 6-phenyl-N3-[4-(trifluoromethyl)phenyl]-1H-pyrazolo[3,4-d]pyrimidine-3,4-diamine (60.0 mg, 162 $\mu$mol, 5.41% yield) as a white solid. $^1$H NMR (500 MHz, dmso) $\delta_H$ 7.45 – 7.51 (m, 3H), 7.56 (br., s, 2H), 7.61 (d, 2H), 7.80 (d, 2H), 8.34 – 8.40 (m, 2H), 8.86 (s, 1H), 12.75 (br., s, 1H). LCMS(ESI): $[M+H]^+$ m/z: calcd 371.14; found 371.2.The data were in agreement with those published.[S3]

### Methodology for Antiviral and ADME Assays

#### EV-D68 Antiviral Screening Assay in RD Cells

For the full protocol, please visit the following entry on protocols.io:

**DOI** dx.doi.org/10.17504/protocols.io.yxmvmywybv3p/v1

RD cells (2k per well) were seeded in 96 Well Black/Clear Bottom Plates and incubated overnight. On the following day, cell density was determined using Trypan Blue. Infection media containing DMSO was prepared, and deep 96-well plates were filled with 800 $\mu$L per well, except for column 12. Diluted compounds were added to the designated wells. After a 2.5-hour incubation at 37°C with 5% CO2, the test plate was infected with EV-D68 (strain US/MO/14-18949, ATCC/BEI Resources) at an MOI of 1. After 24 hours of infection, media was aspirated, and cells were fixed with 4% Formaldehyde + PBS.

Following fixation, formaldehyde was carefully removed and the monolayers were washed once with PBS. PBS was discarded, and cells were incubated with 0.1% Triton X-100 for 20 minutes. The monolayer was washed three times with PBS. Cells were then incubated with 0.3% BSA/PBS for 30 minutes. The primary antibody, Anti-Enterovirus D68 VP1 (GTX132313), was diluted in 0.3% BSA/PBS and added to the wells. Following a 1-hour incubation, cells were washed three times with PBS. The secondary antibody, goat anti-rabbit IgG (H+L) Alexa Fluor 488 (ab150077), diluted in 0.3% BSA/PBS, was added to the cells and incubated for an additional hour, protected from light. The monolayer was washed three times with PBS and counterstained with DAPI. The final PBS wash was left on the cells for imaging.

Imaging was performed using the Cytation 1 system, Viral infection and cell viability were assessed using fluorescence imaging to quantify Alexa Fluor 488 and DAPI signals. Images were analyzed using Gen5 software to calculate infection rates and determine antiviral efficacy.

Rupintrivir was used as a positive antiviral control. RD cell cytotoxicity was determined by MTT assays on uninfected cells.

#### EV Panel Antiviral Screening Assays

For the full protocol, please visit the following entry on protocols.io:

**DOI** dx.doi.org/10.17504/protocols.io.5qpvo9jddv4o/v1

Cell monolayers were prepared in 96-well plates and exposed to eight serial half-log10 concentrations of test compounds, along with infected and uninfected controls, and a known active drug. After virus inoculation and incubation until >80% CPE was observed in virus controls, cell viability was quantified using neutral red staining and spectrophotometric analysis at 540 nm.

#### ADME Assays - LogD

For the full protocol, please visit the following entry on protocols.io:

**DOI** dx.doi.org/10.17504/protocols.io.e6nvw14kdlmk/v1

LogD was measured at pH 7.4 using the shake-flask method, optimised for high-throughput experimentation.[S4]

#### ADME Assays - KSOL

For the full protocol, please visit the following entry on protocols.io:

**DOI** dx.doi.org/10.17504/protocols.io.j8nlk8y41l5r/v1

Kinetic aqueous solubility was measured using the shake-flask method.

#### ADME Assays - MDCK Permeability

For the full protocol, please visit the following entry on protocols.io:

**DOI** dx.doi.org/10.17504/protocols.io.n2bvjne6ngk5/v1

Membrane permeability was measured by assessing the flux of compound across monolayers of Madine Darby Canine Kidney II (MDCKII) cells transfected with the human Multidrug Resistance 1 gene (MDR1), expressing the efflux transporter protein: P-gp.[S5–S11] For broader screening, the rate of apical to basolateral (A→B) permeability was determined. In some cases, the B→A direction was measured, in order to obtain the efflux ratio (ER = B→A / A→B).

#### ADME Assays - Human and Mouse Microsomal Stability

For the full protocol, please visit the following entry on protocols.io:

**DOI** dx.doi.org/10.17504/protocols.io.5qpvokdb9l4o/v1

Test compounds in 96-well plate format were assessed for their susceptibility to metabolism by liver enzymes, primarily cytochrome P450 (CYP) enzymes. Using NADPH supplemented human liver microsomes (HLM) or mouse liver microsomes (MLM).

Microsomal incubations are carried out in 5 aliquots of 30 $\mu$L (one for each time point). The liver microsomal incubation medium comprises phosphate buffer (100 mM, pH 7.4), $MgCl_2$ (3.3 mM), NADPH (3 mM), glucose-6-phosphate (5.3 mM), glucose-6-phosphate dehydrogenase (0.67 units/mL) with 0.415 mg of liver microsomal protein per mL. In the negative control reactions, the NADPH-cofactor system is substituted with phosphate buffer. Test compounds (2 $\mu$M, final acetonitrile concentration 1.6%) are incubated with microsomes at 37°C, shaking at 100 rpm. Five time points over 40 minutes are analysed (0, 7, 15, 25, and 40 min). The reactions are stopped by adding 5 volumes of acetonitrile with internal standard to incubation aliquots, followed by protein sedimentation by centrifuging at 5500 rpm for 5 minutes. Supernatants are analyzed using the HPLC-MS/MS system coupled with a tandem mass spectrometer. The elimination constant ($k_{el}$), half-life ($t_{1/2}$), and intrinsic clearance ($Cl_{int}$) are determined in a plot of ln(AUC) versus time, using linear regression analysis:

$$k_{\mathrm{el}} = -slope \tag{1}$$

$$t_{1/2} = \frac{\ln 2}{k_{\mathrm{el}}} \tag{2}$$

$$Cl_{\mathrm{int}} = \frac{\ln 2}{t_{1/2}} * \frac{Incubation\, Volume(\mu L)}{Protein\, In\, Incubation(mg)} \tag{3}$$

### Antiviral and ADME profiling of Published Capsid-Targeting EV-D68 Antivirals

Table S1: Measured EV-D68 antiviral activity (RD cells) and and *in-vitro* ADME profile of published enterovirus capsid-interacting antiviral compounds. $EC_{50}$ and $CC_{50}$ values are stated as the geometric mean.

| Identifier | Synonym | Ref | Structure | $EC_{50}[CC_{50}]$ ($\mu$M) | LogD (pH 7.4) | KSOL ($\mu$M) | MDCKII-MDR1 $P_{app}A\rightarrow B$ ($10^{-6}$ cm $s^{-1}$) | $Cl_{int}$ HLM , MLM ($\mu$L $min^{-1}$ $mg^{-1}$) |
|---|---|---|---|---|---|---|---|---|
| **1** | Pleconaril | S12–S14 | 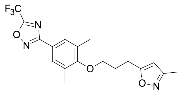 | <0.0295[>50] | 4.2 | 18 | 1.4 | 12 , 22 |
| **2** | CP-11526092 | S15,S16 | 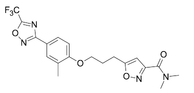 | 0.043[>42] | ≥4.5 | 9 | 2.3 | 11 , 24 |
| **3** | Vapendavir | S17–S20 | 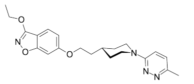 | 0.170[>50] | 3 | 16 | 2.4 | 42 , 95 |
| **S-57** | BTA-188 | S1 | 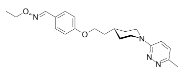 | >2.24[>50] | 4.4 | 8 | 7.2 | 18 , 38 |
| **S-58** | Pirodavir | S21 | 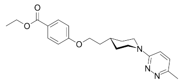 | 0.277[>50] | 4.1 | 9 | 9 | 708 , 580 |
| **S-59** | NLD-22 | S2 | 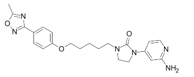 | <5.57[>25.6] | 2.5 | 10 | 1.9 | 33 , 30 |
| **S-60** | OBR-5-340 | S3 | 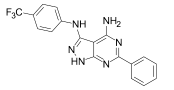 | 7.16[>50] | ≥4.5 | 2 | 1.8 | <10 , 11 |

### Published EV Capsid Structural Data

#### List of Ligand-Bound Crystal and Cryo-EM Structures of VP1

Table S2: List of known published enterovirus capsid structures with a small molecule ligand bound to VP1. For further structural work relating to numerous WIN compounds and how they bind to the HRV-14 capsid VP1, see the seminal work of Rossmann et al.[S22] XRD = X-Ray Diffraction, CryoEM = Cryo Electron Microscopy, EV = Enterovirus, POV = Poliovirus, HRV = Human Rhinovirus, Cox = Coxsackievirus.

| Compound | Method | PDB ID | Organism | Resolution (Å) |
|---|---|---|---|---|
| 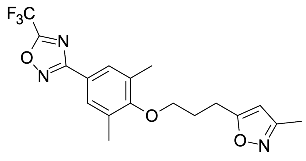 **1** (pleconaril) | XRD | 1C8M | HRV16 | 2.80 |
| | XRD | 1NA1 | HRV14 | 3.30 |
| | XRD | 1NCQ | HRV14 | 2.50 |
| | XRD | 1NCR | HRV16 | 2.70 |
| | XRD | 1ND3 | HRV16 | 2.80 |
| | XRD | 4WM7 | EV-D68 | 2.32 |
| | CryoEM | 7OZK | EV-70 | 2.31 |
| | CryoEM | 7TAG | EV-D68 | 2.70 |
| | CryoEM | 8AYY | POV-3 (VLP) | 2.60 |
| 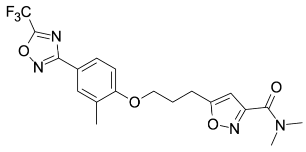 **2** (CP-11526092)) | CryoEM | 7TAF | EV-D68 | 2.00 |

Table S2: (Continued)

| Compound | Method | PDB ID | Organism | Resolution (Å) |
|---|---|---|---|---|
| 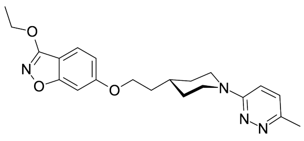 **3** (Vapendavir) | XRD | 3VDD | HRV2 | 3.20 |
| | XRD | 1D4M | CoxA9 | 2.9 |
| | XRD | 1PIV | POV-3 | 2.9 |
| 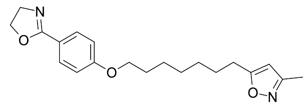 | XRD | 2R04 | HRV-B14 | 3.0 |
| **S-61 (Disoxaril)** | XRD | 3ZFF | EV-A71 | 4.40 |
| | XRD | 3ZFG | EV-A71 | 3.2 |
| | CryoEM | 7OZL | EV-70 | 2.74 |
| 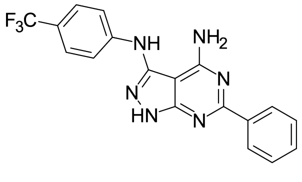 **S60** (OBR-5-340) | CryoEM | 6SK5 | HRV-B5 | 3.60 |
| 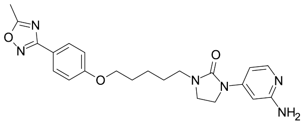 **S-59** (NLD-22) | CryoEM | 6LQD | EV-A71 | 3.26 |

Table S2: (Continued)

| Compound | Method | PDB ID | Organism | Resolution (Å) |
|---|---|---|---|---|
| 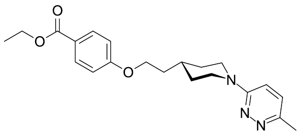 **S-58** (Pirodavir) | XRD | 1PO2 | POV-1 | 2.9 |
| | XRD | 1VBC | POV-3 | 2.8 |
| 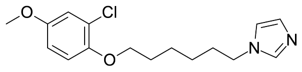 **S-62** (SCH 38057) | XRD | 1HRI | HRV-B14 | 3.00 |

#### 3D Overlays

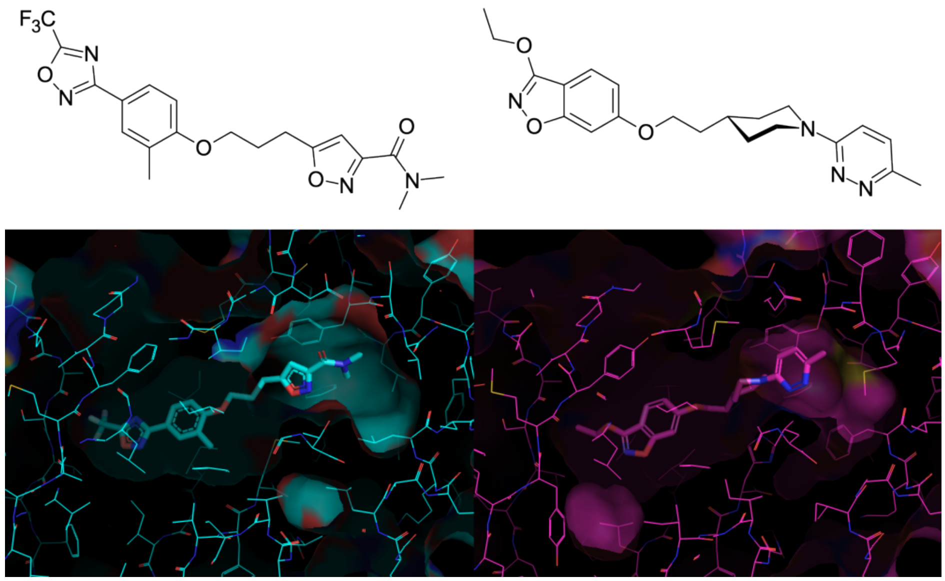

Figure S1: Aligned protein structures of EV-D68 (cyan, 7TAF)[S16] and HRV-A2 (magenta, 3VDD)[S20] capsid VP1 bound to CP-11526092 and vapendavir, respectively.

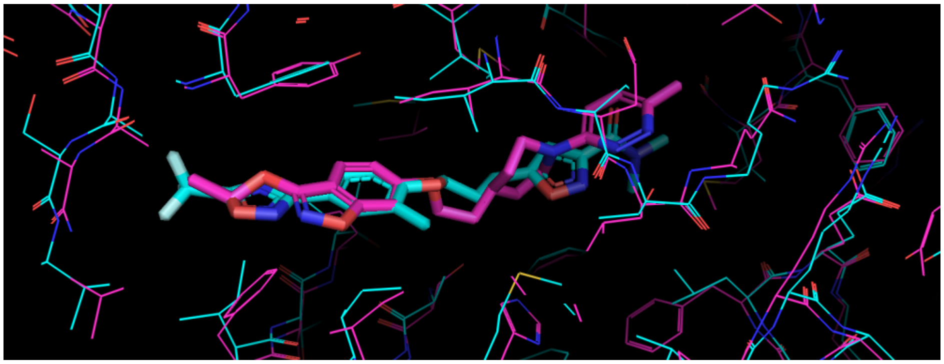

Figure S2: Overlay of 7TAF (cyan) and 3VDD (magenta), showing the relative orientation of the bound ligand with respect to the capsid VP1 hydrophobic pocket. Analysis such as this guided the design of vapendavir/pleconaril hybrid compounds seen in Table 5 of the main text.

### References

(S19) Aviragen Antiviral Drug Vapendavir Fails Phase IIb Study in Asthma Patients. https://www.genengnews.com/news/aviragen-antiviral-drug-vapendavir-fails-phase-iib-study-in-asthma-patients/.

(S20) Feil, S. C.; Hamilton, S.; Krippner, G. Y.; Lin, B.; Luttick, A.; McConnell, D. B.; Nearn, R.; Parker, M. W.; Ryan, J.; Stanislawski, P. C.; Tucker, S. P.; Watson, K. G.; Morton, C. J. An Orally Available 3-Ethoxybenzisoxazole Capsid Binder with Clinical Activity against Human Rhinovirus. *ACS Medicinal Chemistry Letters* **2012**, *3*, 303–307.

(S21) Andries, K.; Dewindt, B.; Snoeks, J.; Willebrords, R.; Van Eemeren, K.; Stokbroekx, R.; Janssen, P. A. In Vitro Activity of Pirodavir (R 77975), a Substituted Phenoxy-Pyridazinamine with Broad-Spectrum Antipicornaviral Activity. *Antimicrobial Agents and Chemotherapy* **1992**, *36*, 100–107.

(S22) Badger, J.; Minor, I.; Oliveira, M. A.; Smith, T. J.; Rossmann, M. G. Structural Analysis of Antiviral Agents That Interact with the Capsid of Human Rhinoviruses. *Proteins: Structure, Function, and Bioinformatics* **1989**, *6*, 1–19.
